# Hallmark molecular and pathological features of POLG disease are recapitulated in cerebral organoids

**DOI:** 10.1101/2023.09.16.558087

**Authors:** Anbin Chen, Tsering Yangzom, Yu Hong, Bjørn Christian Lundberg, Gareth John Sullivan, Charalampos Tzoulis, Laurence A. Bindoff, Kristina Xiao Liang

**Affiliations:** Department of Clinical Medicine (K1), University of Bergen, Bergen, Norway; Neuro-SysMed, Center of Excellence for Clinical Research in Neurological Diseases, Haukeland University Hospital, Bergen, Norway; Department of Neurosurgery, Xinhua Hospital, Shanghai Jiaotong University School of Medicine, Shanghai, China; Center for Diagnosis and Treatment of Cranial Nerve Diseases, Shanghai Jiaotong University, Shanghai, China; Centre for International Health, University of Bergen, Bergen, Norway; Department of Biomedicine, University of Bergen, Bergen, Norway; Department of Molecular Medicine, Institute of Basic Medical Sciences, University of Oslo, Oslo, Norway; Institute of Immunology, Oslo University Hospital, Oslo, Norway; Hybrid Technology Hub Centre of Excellence, Institute of Basic Medical Sciences, University of Oslo, Oslo, Norway; Department of Pediatric Research, Oslo University Hospital, Oslo, Norway; National Advisory Unit for Congenital Metabolic Diseases, Oslo University Hospital, Oslo, Norway

**Author notes:** **Correspondence to:** Kristina Xiao Liang, Department of Clinical Medicine (K1), University of Bergen, Jonas Lies vei 87, P. O. Box 7804, 5021 Bergen, Norway. These authors contributed equally to this work as corresponding authors.

**Keywords:** iPSC, cortical organoids, POLG, mitochondrial function, neuron

## Abstract

In our research, we developed a 3D brain organoid model to study POLG-related encephalopathy, a mitochondrial disease stemming from *POLG* gene mutations. We utilized induced pluripotent stem cells (iPSCs) derived from patients with these mutations to generate cortical organoids, which exhibited typical POLG disease features, such as altered morphology, neuronal loss, and mtDNA depletion. We also identified significant dysregulation in pathways crucial for neuronal development and function, alongside upregulated NOTCH and JAK-STAT signaling pathways. Metformin treatment ameliorated many of these abnormalities, except for the persistent affliction of inhibitory DA GLU neurons. This novel model effectively mirrors both the molecular and pathological attributes of POLG disease, providing a valuable tool for mechanistic understanding and therapeutic screening for POLG-related disorders and other conditions characterized by compromised neuronal mtDNA maintenance and complex I deficiency.

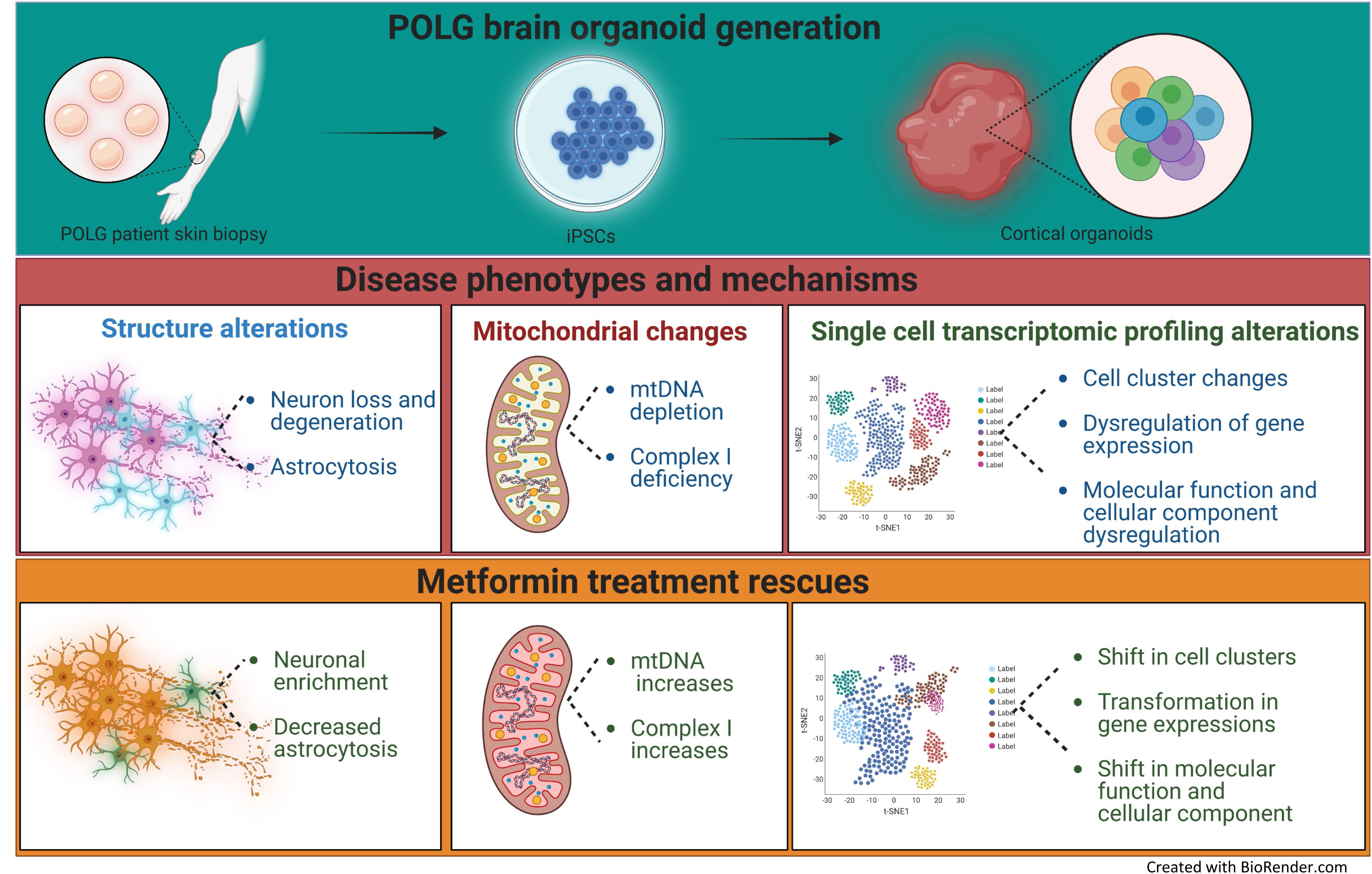

**Highlights:**

- We have successfully developed cortical organoid model that represents POLG-related disease.
- This model effectively replicates both histological and molecular signatures seen in the brains of patients.
- The cortical organoid model displays a range of features common in POLG-related disease, including neurodegeneration, mtDNA depletion, and neuronal complex I deficiency.
- The use of metformin supplementation in this model improved mitochondria protein and reduced cell death.

## Introduction

Mitochondrial diseases represent the largest class of inborn errors of metabolism and comprise mainly monogenic disorders that disrupt the process of oxidative phosphorylation (OXPHOS). One of the most common mitochondrial diseases is caused by mutations in the *POLG* gene encoding the catalytic subunit of the mitochondrial DNA polymerase gamma (Pol γ), which is responsible for replication and repair of mtDNA. Deleterious *POLG* mutations impair mtDNA maintenance and result in quantitative and qualitative mtDNA defects, which in turn cause mitochondrial respiratory chain deficits - mainly affecting complex I and, to a lesser degree, complex IV [1, 2]. Clinically, POLG diseases manifest a broad phenotypical spectrum ranging from pure myopathies [3], to juvenile syndromes with progressive spinocerebellar ataxia and epilepsy [4–7] and devastating infantile hepatic encephalopathy [8]. Moreover, the downstream molecular consequences of *POLG* mutations - defective mtDNA maintenance and complex I deficiency - are also seen in idiopathic forms of age-related neurodegeneration including Parkinson’s disease (PD) and Alzheimer’s disease (AD) [9–11]. Disease caused by *POLG* mutations is, therefore, an excellent model for exploring the pathogenic mechanisms by which impaired mtDNA maintenance and complex I deficiency cause neuronal disease, as well as for screening and assessing potential therapies targeting these processes.

The pathophysiological mechanisms underlying neurodegeneration in mitochondrial disease are poorly understood and there are no effective neuroprotective therapies able to delay or arrest disease progression. Advances in this field require the development of model systems that recapitulate the molecular semiology and cellular phenotypes characterizing human disease. Advances in stem cell technology now allow us to generate complex models using patient-derived induced pluripotent stem cells (iPSCs), reprogrammed into neurons and other brain cell-types. We previously generated iPSCs from POLG patients and differentiated them into neural stem cells (NSCs) and dopaminergic neurons [12, 13]. While this traditional 2D iPSC model (monolayer culture) is a powerful and reproducible tool, it does not reflect the complexity and high-level organization seen at the level of human brain tissue, thus limiting its mechanistic relevance for human disease [14, 15]. This limitation can be mitigated by resorting to brain organoids, i.e., 3D structures composed of multiple cell types that can self-organize to recapitulate embryonic and tissue development in vitro. Brain organoids have been shown to outperform 2D cell culture methods in reflecting the functional, structural, and geometric characteristics of brain tissue [16–18]. Moreover, since brain organoids can be generated in large numbers, they also have advantages over animal models in drug screening studies. Given that drugs ameliorating neurological disorders can be effective in animal models but fail in clinical trials [19, 20], this emphasizes the need for human cell-based systems to evaluate drug efficacy.

Metformin, a drug primarily used for the management of type 2 diabetes, has demonstrated potential therapeutic benefits in a variety of other conditions including cancer, cardiovascular diseases and neurodegenerative disorders [21]. Its proposed neuroprotective effects can be attributed to several mechanisms: primarily, metformin activates AMP-activated protein kinase (AMPK), a key regulator of cellular energy homeostasis, which in turn can enhance mitochondrial function and cell survival - critical factors in conditions such as POLG-related disorders characterized by mitochondrial dysfunction [22]. In addition, metformin possesses anti-inflammatory properties, exerted through the inhibition of the NF-κB signaling pathway [23], which might help in managing the inflammatory aspects of neurodegenerative diseases. Another potential benefit of metformin in this context is its ability to stimulate autophagy, a cellular recycling process that maintains neuronal health by degrading damaged proteins and organelles [24]. This effect could potentially contribute to the clearance of disease-associated pathogenic proteins and improve neuronal function. Nonetheless, additional research and clinical trials are necessary to validate the effectiveness and safety of metformin in treating neurodegenerative disorders, particularly those resulting from mitochondrial dysfunction such as POLG-related diseases.

We present a novel iPSC-derived cortical organoid model using cells derived from two patients with POLG encephalopathy, one harboring the homozygous c2243 G>C and one carrying compound heterozygous c.1399 G>A and c2243 G>C mutations (resulting in amino acid changes p. A467T and p. W748S, respectively). Our patient-derived organoids exhibited striking morphological abnormalities, respiratory complex I deficiency, and mtDNA depletion, accurately recapitulating the key histological and molecular features observed in POLG disease in the human brain. Notably, we found that supplementation with metformin effectively mitigated these phenotypes, offering a potential therapeutic avenue for POLG-related disorders.

## Materials and Methods

### Ethics Approval

The project was approved by the Western Norway Committee for Ethics in Health Research (REK nr. 2012/919).

### Generation and maintenance of iPSCs

Skin fibroblasts were obtained from punch biopsies of two patients carrying *POLG* mutations. One patient was homozygous for the c.2243G>C; p.W748S mutation (WS5A), and the other was compound heterozygous for the c.1399G>A/c.2243G>C; p.A467T/W748S mutations (CP2A). The patient identifiers WS5A and CP2A were previously used in our published work [7, 25]. Isogenic control for homozygous for the c.2243G>C; p.W748S mutation (WS5A), along with controls who were matched for age and gender, were included in the study. This group also encompassed iPSCs derived from Detroit 551 fibroblasts (ATCC CCL 110TM), CRL2097 fibroblasts (ATCC CRL-2097™), and AG05836. To generate iPSCs, the skin fibroblasts were reprogrammed using retroviral or Sendai virus vectors containing the coding sequences of human OCT4, SOX2, KLF4, and c-MYC. The iPSCs were maintained under feeder-free conditions as previously described [13, 26–29]. All iPSC lines, including the patient-derived lines and control lines, were maintained following established protocols [27]. Regular monitoring for mycoplasma contamination was performed using the Myco Alert™ Mycoplasma Detection Kit (Lonza, #LT07-218) to ensure the integrity of the cell lines.

### Karyotype analyses

Human G banding karyotyping was performed using the protocol as reported previously [27].

### Generation of cortical organoid

To generate cortical organoids from iPSCs, we followed a previously described protocol [30, 31]. Initially, feeder-free iPSCs were cultured in E8 medium for a minimum of 7 days prior to differentiation. The iPSC colonies were dissociated using Accutase (Life Technologies, #A11105-01) in a 1:1 mixture with phosphate-buffered saline (PBS), incubated at 37°C for 10 minutes, and then centrifuged at 100 × g for 3 minutes. The resulting single cells were diluted in neural induction media (Supplementary Table 1) and approximately 9,000 viable cells were seeded into each well of 96-well ultra-low attachment tissue culture plates (S-BIO, #MS-9096UZ) in 150 μl of neural induction media. The plates were kept in suspension and rotated at 85 rpm for 24 hours, with the addition of 50 μM Y-27632 Rock Inhibitor (Biotechne Tocris, #1254) to promote the formation of EBs. To minimize undirected differentiation, we employed dual SMAD inhibition and canonical Wnt inhibition during the initial phase. On day 2, half of the media in each well was replaced with human neural induction media containing 50 μM ROCK inhibitor. On days 4, 6, and 8, 100 μl of the medium was replaced with 150 μl of neural induction media without the ROCK inhibitor. After 10 days, the EBs were transferred to 6-well ultra-low attachment tissue culture plates (Corning, #3471) and cultured in neural differentiation media (Supplementary Table 2) without vitamin A for the next 8 days using an orbital shaker. This step aimed to induce the formation of cortical organoids. On day 18, the organoids were further matured in neural differentiation media supplemented with vitamin A (Supplementary Table 3). Media changes were performed every 3-4 days, and the addition of Brain Derived Neurotrophic Factor (BDNF, Peprotech, #450-02) and ascorbic acid (Sigma-Aldrich, #92902-500G) facilitated long-term neural maturation.

### Metformin treatment of cortical organoids

For phenotype rescue experiments, organoids were treated with 250 μM metformin (Sigma-Aldrich, #317240) from day 6 of differentiation. The medium was changed every two days and the organoids were grown for 3 months. An equivalent amount of vehicle of Dimethylsulfoxide (DMSO, Sigma-Aldrich, #153087) was added to grow untreated organoids.

### Snap freezing and embedding

Each human cerebral organoid was fixed in 4% paraformaldehyde (PFA, Thermo Fisher Scientific, #28908) in PBS overnight at 4°C, dehydrated with 30% sucrose in PBS. The samples were then embedded in gelatin solution (Sigma-Aldrich, #G1393) and snap frozen in boiled liquid nitrogen. The samples were then embedded in optimal cutting temperature (O.C.T) compound (Thermo Fisher Scientific, #23-730-57). Cryostat sections (15 µm) were cut and mounted Superfrost™ adhesion slide (Thermo Fisher Scientific, #J1800AMNZ).

### Immunofluorescence staining

Mounted sections were incubated for 1 hour at room temperature with and blocked using blocking buffer containing (Sigma-Aldrich, #G9023) and 0.1% (v/v) Triton X100 (Sigma-Aldrich, #9036-19-5), and then incubated with primary antibodies diluted in blocking solution overnight at 4°C. The following primary antibodies were used for immunostaining: Nanog (rabbit, 1:100; Abcam, #ab80892), Oct4 (rabbit, 1:100; Abcam, #ab19857), SSEA4 (mouse, 1:200; Abcam, #ab16287,), NeuN (rabbit, 1:500; Cell Signaling, #24307S), MAP2 (rabbit, 1:1000; Abcam, #ab5392), β-tubulin III (TUJ1) (mouse, 1:1000; Abcam, #ab78078), Synaptophysin (mouse, 1:500; Proteintech, #17785-1-AP), PSD-95 (rabbit, 1:500; Proteintech, #20665-1-AP), SOX2 (rabbit, 1:100; Abcam, #ab97959), NDUFB10 (rabbit, 1:350; Abcam 196019), TFAM (mouse, 1:1000; Abcam #ab119684), TOMM20 (mouse, 1:350; Abcam, #ab56783), GFAP (chicken, 1:500; Abcam #ab4674), DCX (mouse, 1:200; Abcam #ab135349), SATB2 (rabbit, 1:400; Abcam #ab4674), CTIP2 (rat, 1:500; Abcam, #ab18465), OLIG2 (rabbit, 1:500; Abcam, #ab42453). Alexa Fluor Dyes (Invitrogen, #A11008, #A21449, #A11012, #A21141, #A11042, #A21236) were used at 1:800 dilution as secondary antibodies. Slides were mounted using ProLong^TM^ diamond antifade mounting medium (SouthernBiotech, #0100-20), and analyzed using the Leica TCS SP8 STED 3X (Leica microsystems).

### Fluorescence imaging analysis

Immunofluorescence images were quantitatively measured using Image J software (Image J 1.52a; Wayne Rasband National Institutes of Health, USA). Six to ten regions within cortical layers were randomly selected for quantitative evaluation. To measure fluorescence intensity, we converted single-channel images to 8-bit images using Image J. To eliminate errors caused by manually selecting thresholds for different photos, we used default thresholds. Next, we chose the default algorithm, and parameters. Finally, the fluorescence intensity was measured by the mean gray value (Mean) with the following formula: Mean = Integrated Density (IntDen)/Area. Data were analyzed and plotted with GraphPad Prism 8.0.2 software (GraphPad Software, Inc). Details of statistical tests and p-values are described in Supplementary Figure 8.

### Single cell RNA sequencing (scRNA-seq) and data analysis

#### Organoid dissociation and single cell isolation

To collect the organoids, they were removed from the culture medium and washed with 1x PBS (Invitrogen, #10010-23). The organoids were then finely cut into pieces of 1-2 mm using ophthalmic scissors. Next, the organoid pieces were digested in 2 ml of CellLive™ Tissue Dissociation Solution (Singleron Biotechnologies, #1190062) at 37°C for 15 minutes in a 15-ml centrifuge tube (Sarstedt, #62.5544.003), with continuous agitation on a thermal shaker. The degree of dissociation was periodically checked under a light microscope. Following digestion, the suspension was filtered through a 40-µm sterile strainer (Greiner#542040). The cells were then centrifuged at 350 x g for 5 minutes at 4°C, and the resulting cell pellets were resuspended in 1 ml of PBS. To assess cell viability and count, the cells were stained with a 0.4% w/v solution of Trypan Blue (Gibco, #15250-061). The cell number and viability were determined using a hemocytometer under a light microscope.

### ScRNA-seq library preparation

scRNA-seq libraries were prepared using the GEXSCOPE™ Single Cell RNAseq Library Kit (Singleron Biotechnologies, #4161031) following the manufacturer’s instructions. Briefly, the single-cell suspension was adjusted to a concentration of 3x105 cells/ml with PBS and loaded onto an SD microfluidic chip to capture 6,000 cells. Paramagnetic beads conjugated to oligo(dT) probes with unique molecular identifiers (UMIs) and barcodes were added, and the cells were lysed. The polyadenylated mRNA bound to the beads was extracted, reverse transcribed into cDNA, and amplified by PCR. The resulting cDNA was fragmented and ligated to indexed Illumina adapters. The final amplified library’s fragment size distribution was analyzed using an Agilent Fragment Analyzer.

### Library sequencing

The library concentration was calculated using the Qubit 4.0 fluorometer and the libraries were pooled in an equimolar fashion. The single cell libraries were sequenced on an Illumina NovaSeq 6,000 using a 2x150-bp approach to a final depth of 90 GB per library. The reads were demultiplexed according to the multiplexing index sequencing on Illumina’s Base Cloud platform.

### Transcriptome data pre-processing

The scRNA-seq data was pre-processed using the CeleScope® software (v.1.3.0; www.github.com/singleron-RD/CeleScope; Singleron Biotechnologies GmbH) to generate raw data with default parameters. Low quality reads were discarded, and the remaining sequences were mapped to the human reference genome GRCh38 using STAR (https://github.com/alexdobin/STAR). Gene annotation was performed using Ensembl 92. The assignment of reads to genes was done using featureCount (https://subread.sourceforge.net/), resulting in a count matrix file that contained the number of Unique Molecular Identifiers (UMIs) for each gene within each cell.

Subsequent analysis was carried out using the scanpy package in Python. Quality control metrics, such as the number of genes detected per cell (nFeature_RNA) and the percentage of mitochondrial UMIs (percent_mt), were extracted from the gene count matrix using the calculate_qc_metrics function. To remove non-viable cells, cells with a high percentage of mitochondrial counts (>20%) were filtered out. Cells with a high number of detected genes (>5000), indicative of potential doublets, were also excluded. Additionally, cell debris characterized by a low number of detected genes (<200) were removed from the dataset.

### Sample integration, dimensionality reduction and cell clustering

The downstream bioinformatic analysis was performed by combining three samples using the *concatenate* function from the *anndata* package in *Python.* Data were log-normalized such that the total count for each cell was 10,000. Highly variable genes (HVGs) were identified using dispersion-based methods with mean between 0.1 and 8 and dispersion of 0.5 and above using the function *highly_variable_genes* from *scanpy*. Principal component analysis (PCA) was performed on expressions from HVGs. The top 17 principal components were retrained from 50 by checking explained variance ratio of PCA instances. We used *neighbors’* function in *scanpy* to calculate the neighborhood graph with 20 neighbors and 17 PC. The Uniform Manifold Approximation and Projection (UMAP) algorithm was used to calculate the reduced dimensions (Supplementary Figure 1) from neighborhood graph with minimum distance of 0.5, spread scale of 1 and 200 maximum iterations. The cell clusters were identified using the unsupervised network-based *leiden* algorithm [32] provided in the *scanpy* package with a resolution of 0.5 on the top 17 principal components.

### Cell type annotation

To identify distinct cell populations based on shared and unique patterns of gene expression, we performed dimensionality reduction and unsupervised cell clustering using single cell multi-resolution marker-based annotation scMRMA [33] (https://github.com/JiaLiVUMC/scMRMA) with cutoff of significant p value smaller than 0.05 for the fisher test for the significant cell type enrichment from 20 nearest neighbors. Cell type marker genes were obtained from the PanglaoDB database (https://panglaodb.se/).

### Differential gene expression (DEG) analysis

The marker genes of the individual clusters were found by using the differential gene expression analysis algorithm scanpy.tl.rank_genes_groups in Python with the following parameters sc.tl.rank_genes_groups (adata, groupby=’leiden’, method=’wilcoxon’, corr_method=’bonferroni’, use_raw=False). A p-value of 0.05 was used to keep the DEGs that were significant. The top 9 DEGs were shown for each cluster in the figures.

### Gene enrichment analysis

Gene enrichment analysis was performed with the package *gseapy* in Python with the databases “GO_Biological_Process_2021”, “GO_Molecular_Function_2021”, “GO_Cellular_Component_2021”, “Reactome_2016”, “KEGG_2016” and “KEA_2015”. The information on the software used for scRNA-seq analysis was listed in Supplementary Table 4

### Statistical analysis

The data were presented as mean ± standard deviation (SD) for samples with a minimum size of three. The normality of the data distribution was assessed using the Shapiro-Wilk test. Outliers were identified using the ROUT method. For variables with a non-normal distribution, statistical significance was determined using the Mann-Whitney U-test, while variables with a normal distribution were analyzed using a two-sided Student’s t-test. Graphs and statistical analyses were performed using GraphPad Prism 8.0.2 software (GraphPad Software, Inc). A p-value of ≤ 0.05 was considered statistically significant. Detailed information on the specific statistical tests conducted and corresponding p-values can be found in Supplementary Table 5.

## Results

### Generation of cortical organoids from iPSCs

We generated iPSCs from the skin fibroblasts of two patients with *POLG* mutations: one patient carrying homozygous c.2243 G > C, p.W748S and another patient with compound heterozygous c.1399 G > A, p.A467T and c.2243 G > C, p.W748S. We also included skin fibroblasts from two neurologically healthy individuals as controls, as previously described in our previous report [27]. The patient’s symptoms included progressive spinocerebellar ataxia, peripheral neuropathy, migraine-like headaches, and extraocular myopathy, leading to progressive external ophthalmoplegia, as reported in the previous study [7, 34]. The reprogrammed patient iPSCs exhibited standard karyotyping compared to the original fibroblasts, alongside positive expression of SOX2, OCT4, and NANOG (Supplementary Figure 2 and 3).

Cortical organoids were generated by inducing neural differentiation of iPSCs through the formation of EBs in suspension culture. This was achieved using a neural induction protocol that involved dual SMAD inhibition and canonical Wnt inhibition (Figure 1A) [31]. This led to the creation of cortical region brain tissues, termed as cortical organoids; this took 2 days for EB formation, 10 days for neural identity appearance and 20-30 days for definitive brain region formation (Figure 1B). On day 50, the organoids matured into large, complex heterogeneous tissues up to 3 mm in diameter (Figure 1C) that survived for 4-5 months when kept in a spinning rotator.

**Figure 1:**
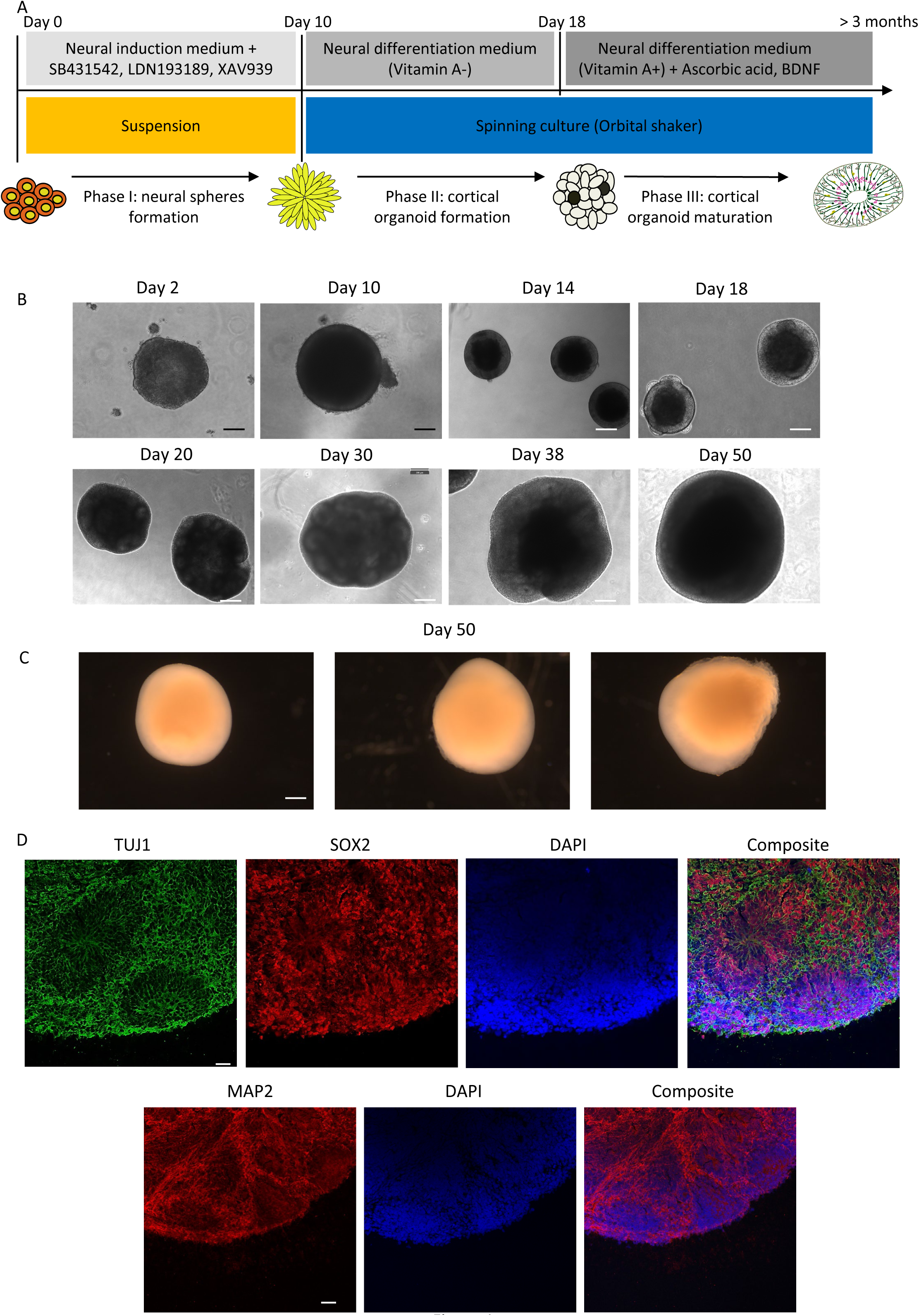
Generation of cortical organoids from iPSCs. (A) The differentiation protocol consisted of three phases. In Phase I, neural induction and neural sphere formation were achieved by generating embryoid bodies in stationary suspension 3D culture using the dual SMAD inhibition and canonical Wnt inhibition approach. In Phase II, cortical organoid formation was initiated by transferring the cells into spinning culture using an orbital shaker and culturing them in neural differentiation medium without vitamin A to promote regionalization factors and cortical organization. In Phase III, cortical organoid maturation was facilitated by maintaining the organoids in neural differentiation medium supplemented with vitamin A, BDNF, and ascorbic acid for long-term neural maturation. (B) Representative phase contrast images were captured at various time points during the differentiation process, including day 2, 10, 14, 18, 20, 30, 38, and 50. These images displayed the changing cell morphology throughout the differentiation process. The black scale bar represents 100 µm, and the white scale bar represents 300 µm. (C) The morphology of three individual organoids on day 50 of differentiation was examined. The image displayed the diversity in size and structure among the organoids. The scale bar represents 400 µm. (D) Immunofluorescent imaging was performed on cryosectioned organoids on day 30 of differentiation. The staining revealed the presence of the newborn neural marker TUJ1 (green), neural progenitor marker SOX2 (red), and mature neural marker MAP2 (red). Nuclei were stained with DAPI (blue). The scale bar represents 100 µm.

At different stages of development, we noted different organoid characteristics. At the early stage (30 days), small organoids expressed the newborn neural marker Tubulin beta III (TUJ1), ventricular zone (VZ) marker SOX2, as well as the mature neural marker MAP2 (Figure 1D and 2A). By 60 days (intermediate stage), organoids showed more pronounced SOX2 expression (Figure 2B). By the late stage (70 days), large neural tubes expressing SOX2 and MAP2 were observed in the outer layers (Figure 2C). An increase in SOX2 levels was noted from day 30 to day 60 and day 70, while MAP2 levels remained stable (Figure 2D and E).

**Figure 2:**
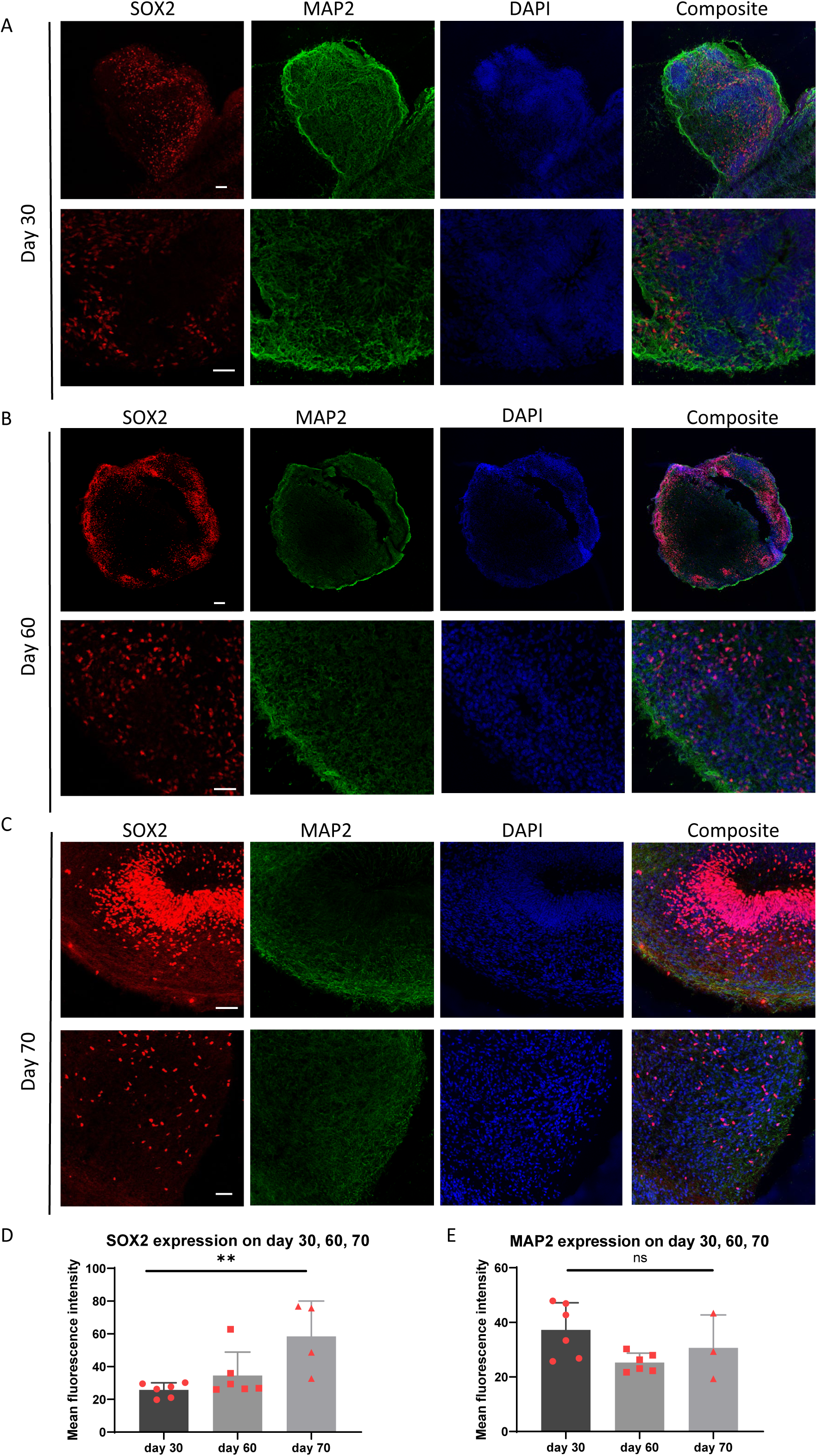
Characterization of cortical organoids from iPSCs during differentiation. (A-C) Cryosectioned organoids from day 30 (A), day 60 (B), and day 70 (C) of differentiation were subjected to immunofluorescent imaging. The staining revealed the presence of the neural progenitor marker SOX2 (red) and the mature neural marker MAP2 (green). Nuclei were counterstained with DAPI (blue). The scale bar represents 100 µm. (D-E) Quantitative measurements were performed to assess the expression levels of SOX2 (D) and MAP2 (E) on day 30, 60, and 70 of differentiation. The mean fluorescence intensity was measured, and statistical analysis was conducted using one-way ANOVA. Significance levels are indicated for P values less than 0.05, with ** indicating P < 0.01. Results not reaching statistical significance are denoted as ns (not significant).

Finally, at day 90, the organoids largely consisted of neurons positive for the mature neural marker NeuN, located in the outer layers (Figure 3A). Small proportions of cells expressed GFAP (Figure 3A), MAP2, and SOX2 (Figure 3B). Cortical pyramidal neuronal markers SATB2 and CTIP2 were found in the middle and upper layers of the organoids (Figure 3C), alongside stratified expressions of oligodendrocyte marker oligodendrocyte transcription factor 2 (OLIG2), astrocyte marker GFAP, and neural marker doublecortin (DCX) (Figure 3D).

**Figure 3:**
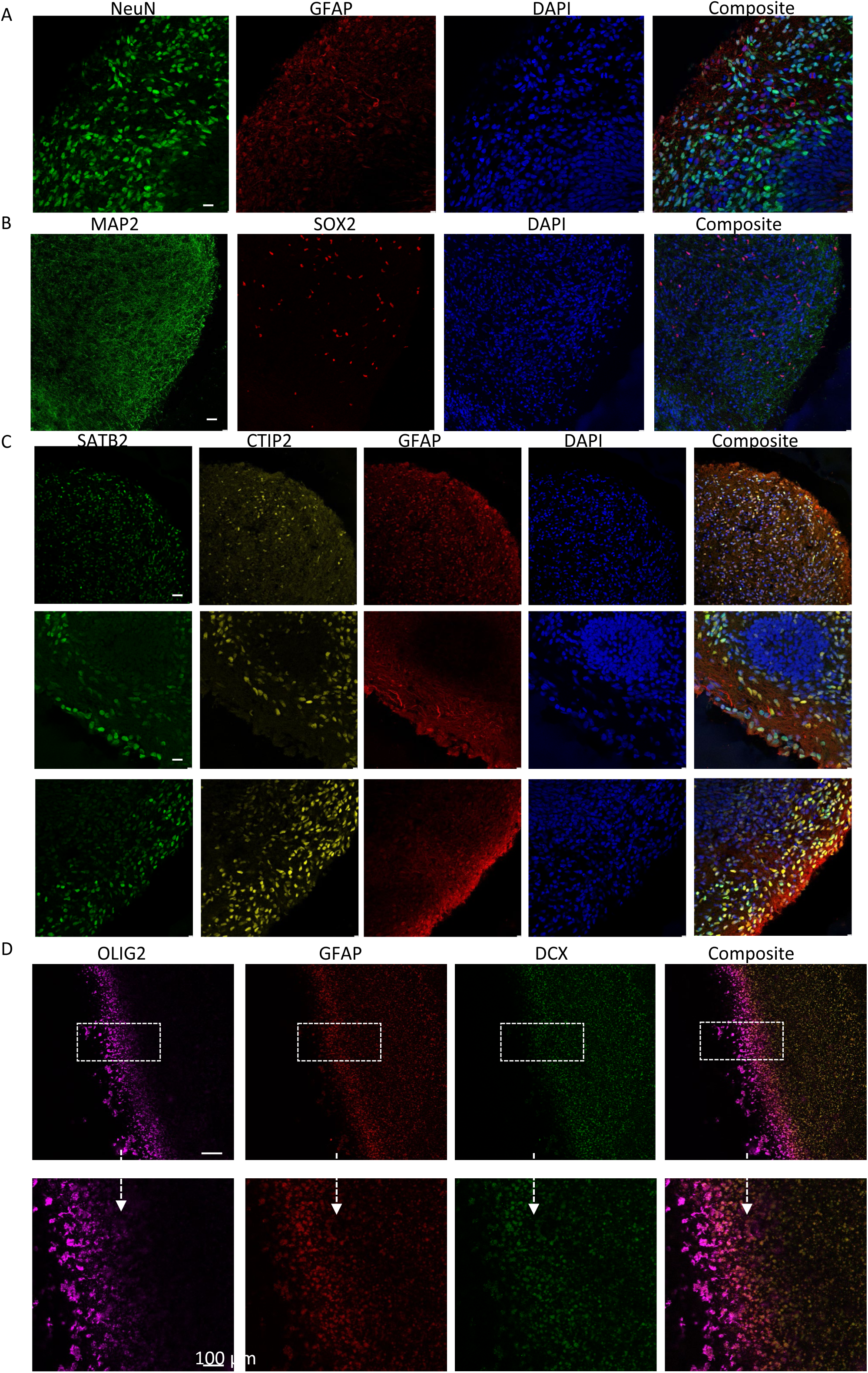
Characterization of cortical organoids from iPSCs at day 90. (A) Cryosectioned organoids on day 90 were subjected to immunofluorescent imaging, revealing the presence of the astrocyte marker GFAP (red) and the mature neural marker NeuN (green). Nuclei were counterstained with DAPI (blue). The scale bar represents 100 µm. (B) Immunofluorescent imaging of cryosectioned organoids on day 90 showed the expression of the mature neural marker MAP2 (green) and the neural progenitor marker SOX2 (red). Nuclei were counterstained with DAPI (blue). The scale bar represents 100 µm. (C) Cryosectioned organoids at day 90 displayed cortical pyramidal neuronal markers SATB2 (green) and CTIP1 (yellow), along with the astrocyte marker GFAP (red). Nuclei were counterstained with DAPI (blue). The scale bar represents 100 µm. (D) Immunofluorescent imaging of cryosectioned organoids at day 90 revealed stratified layers containing the oligodendrocyte marker OLIG2 (purple), the astrocyte marker GFAP (red), and the neural marker DCX (green). Nuclei were counterstained with DAPI (blue). The scale bar represents 100 µm.

We concluded therefore, that this process effectively generated 3D cortical organoids comprising neural tubes, cortical neurons, glial astrocytes, and oligodendrocytes.

### POLG-patient organoids demonstrated significant structural alterations, marked by neuronal loss and an increase in astrocytosis

In contrast with control samples that produced orderly neuroepithelial tissues and subsequently matured into regular cortical organoids, POLG patient-derived samples showed a marked irregularity in their neuroepithelial tissue. These samples matured into structures resembling brain tissue, but without the complete organization seen in control organoids. Aberrant growth was observed throughout the EB development (Figure 4A).

**Figure 4:**
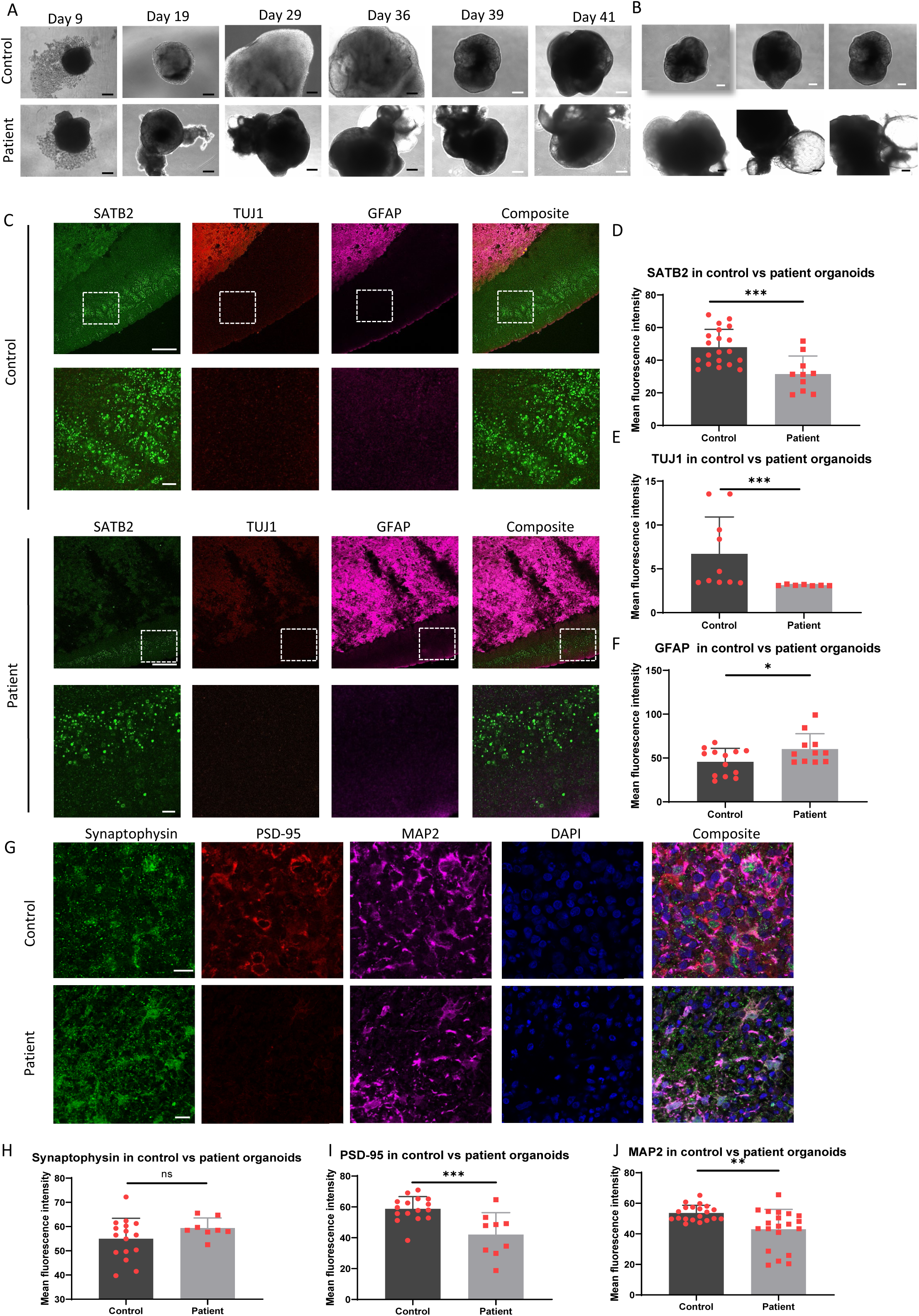
Comparison of lineage markers cortical organoids from control and patient iPSCs. (A) Cell morphology was assessed using phase contrast imaging at day 9, 19, 29, 36, 39, and 41 of differentiation in both control and patient iPSCs. The black scale bar represents 100 µm, and the white scale bar represents 100 µm. (B) Morphology of three individual organoids on day 36 of differentiation was examined in both control and patient iPSCs. The scale bar represents 300 µm. (C) Cryosectioned organoids at 90 days were subjected to immunofluorescent imaging, staining for the cortical pyramidal neuronal marker SATB2 (green), newborn neural marker TUJ1 (green), and reactive astrocyte marker GFAP (red) in both control and patient lines. The scale bar represents 100 µm. (D-F) Quantitative measurements were performed to assess the expression levels of SATB2 (D), TUJ1 (E), and GFAP (F) on day 90 in control and patient cortical organoids. The y-axis represents the mean fluorescence intensity. (G) Immunofluorescent imaging of cryosectioned organoids stained for the presynaptic marker Synaptophysin (green), postsynaptic marker PSD-95 (red), and mature neural marker MAP2 (purple) in both control and patient lines. Nuclei were counterstained with DAPI (blue). The scale bar represents 50 µm. (H-J) Quantitative measurements were conducted to evaluate the expression levels of Synaptophysin (H), PSD-95 (I), and MAP2 (J) in control and patient cortical organoids. The y-axis represents the mean fluorescence intensity. Significance levels are indicated for P values less than 0.05, with * indicating P < 0.05, ** indicating P < 0.01, *** indicating P < 0.001, Results not reaching statistical significance are denoted as ns (not significant).

Specific abnormalities became evident at different developmental stages. At 36 days, peripheral neuroepithelial regions of patient-derived tissue exhibited prominent, fluid-filled cavity (Figure 4B). By day 90, patient organoids demonstrated a significant reduction in ventricle-like structures and neuroepithelial layer thickness compared to control organoids (Figure 4C). Furthermore, staining for cortical neuron marker SATB2 was considerably diminished in patient-derived cortical organoids (Figure 4C and D), as was the case for neuronal marker TUJ1 (Figure 4C and E). In contrast, staining for activated astrocyte marker GFAP was noticeably elevated in patient organoids (Figure 4C and F).

Further analysis was conducted with the presynaptic marker synaptophysin, the postsynaptic marker PSD-95 and mature neural marker MAP2. When compared to controls, patient organoids showed a marked reduction in the levels of PSD-95 (Figure 4G and I) and MAP2 (Figure 4G and J), while synaptophysin levels remained comparable with control samples (Figure 4G and H).

In summary, the POLG patient-derived cortical organoids were distinguished by morphological differences, a reduction in neuronal markers and the excitatory postsynaptic marker, along with an increased presence of astrocytes.

### Patient organoids display altered expressions of mitochondrial proteins

Our prior research highlighted that those patients carrying the same mutations as those modelled in the organoids showed neuronal mtDNA depletion starting in early infancy. This condition is further characterized by progressively increasing levels of mtDNA major arc deletions and respiratory complex I deficiency [34]. This observation was replicated in iPSC-derived 2D NSCs [27] and dopaminergic neurons [13] where we noted both complex I deficiency and mtDNA depletion.

Here, we explored whether these alterations were also manifested in cortical organoids. Complex I levels were examined by conducting multiple immunostainings for the complex I subunit NDUFB10, the mitochondrial mass marker TOMM20, and the mature neuron marker MAP2. We observed a significant decrease in complex I levels in neuronal component of patient organoids compared to controls (Figure 5A and B). However, TOMM20 levels showed no significant change (Figure 5A and C) suggesting that this was not due simply to loss of mitochondrial mass. The NDUFB10/TOMM20 ratio, an indicator of complex I per mitochondrion, was substantially lower in patient samples, signaling a loss of complex I (Figure 5D). Moreover, when we indirectly assessed mtDNA levels using immunostaining for TFAM, patient organoids showed notably lower TFAM levels compared to controls, suggesting that there was indeed mtDNA depletion (Figure 5E and F).

**Figure 5:**
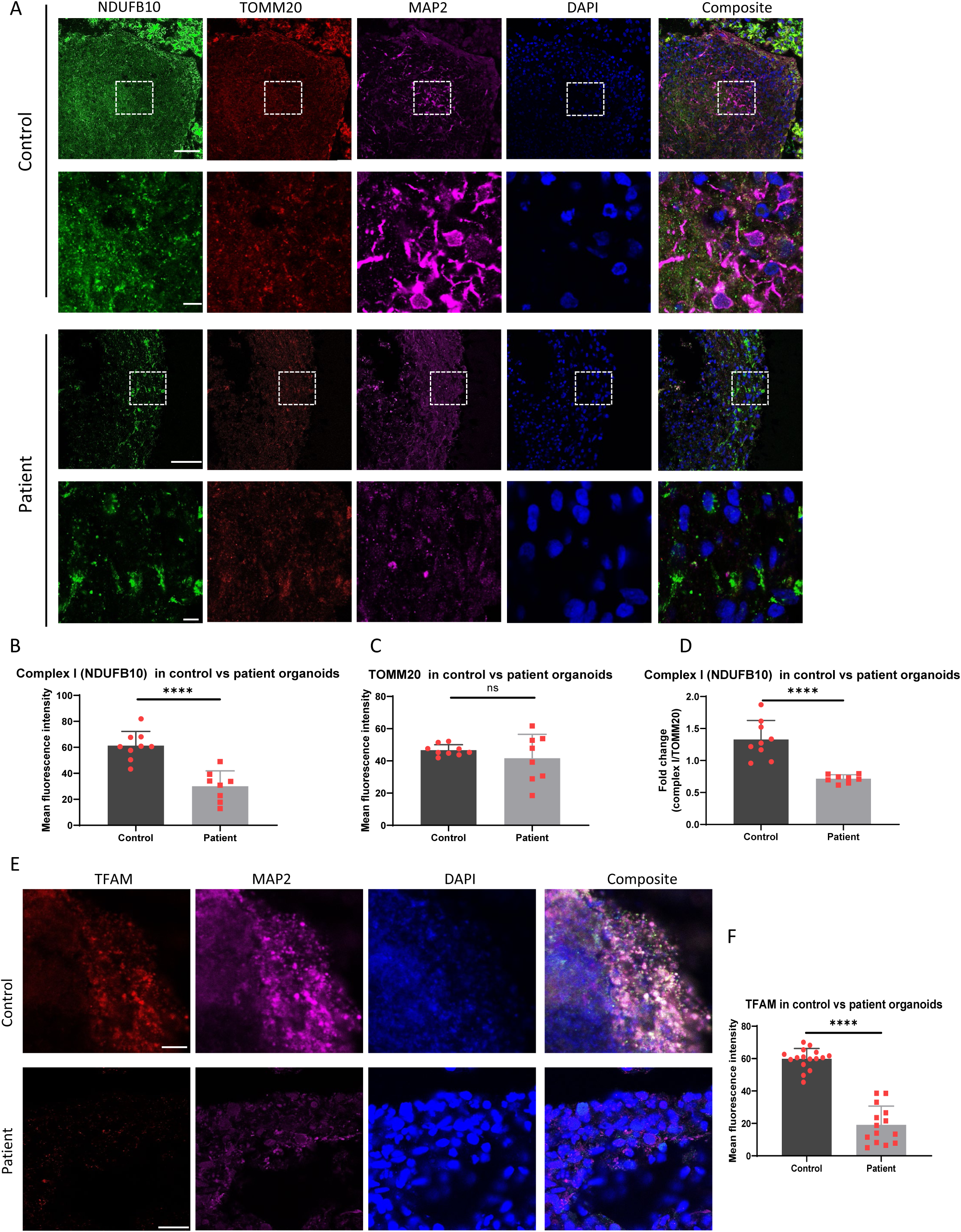
Comparison of mitochondrial related proteins in cortical organoids from control and patient iPSCs. (A) Cryosectioned organoids at 90 days of differentiation were subjected to immunofluorescent imaging, staining for the complex I subunit NDUFB10 (green), outer mitochondrial membrane TOMM20 (red), and mature neural marker MAP2 (purple) in both control and patient iPSC-derived organoids. The scale bar represents 100 µm. (B-D) Quantitative measurements were performed to assess the expression levels of NDUFB10, TOMM20, and MAP2 in control and patient cortical organoids. The y-axis represents the mean fluorescence intensity. (E) Cryosectioned organoids at 90 days of differentiation were subjected to immunofluorescent imaging, staining for the mitochondrial transcription factor A (TFAM), neural progenitor marker SOX2, and mature neural marker MAP2 in both control and patient iPSC-derived organoids. Nuclei were counterstained with DAPI (blue). The scale bar represents 100 µm. (F) Quantitative measurements were conducted to evaluate the expression level of TFAM in control and patient cortical organoids. The y-axis represents the mean fluorescence intensity. Significance levels are indicated for P values less than 0.05, with **** indicating P < 0.000. Results not reaching statistical significance are denoted as ns (not significant).

In conclusion, these findings underscore that iPSC-derived cortical organoids from POLG patients faithfully replicate the key molecular features seen in patient neurons, encompassing neuronal loss, complex I deficiency, and mtDNA depletion.

### Single-cell transcriptomics reveal multiple neuronal cell types in cortical organoids

We employed scRNA-seq to examine the cellular composition and gene expression profiles of the cortical organoids from control iPSCs. We used 4-6 organoids per group for analysis. Our study utilized age and gender-matched control cortical organoids derived from Detroit 551 iPSCs as a comparison group. After applying strict filtering criteria, we analyzed a total of 5,369 cells that expressed 24,576 genes in the control organoid (Supplementary Table 6).

We examined the various cell populations based on cluster gene markers and the expression of known marker genes. The cortical organoids of cell type enrichment analysis in scRNA-seq analysis could be subcategorized further into DA GLU neuron-related genes (*SCG2*, *NSG2*, *MMP3*, *ATP1B1*, *RTN4*, *NEGR1*, *HSP90AA1*, *CELF4*, *LMO3*), dopaminergic neurons (*NEUROD6*, *TMSB10*, *TUBA1A*, *SOX4*, *STMN2*, *PTMA*, *NREP*, *SOX11*, *MLLT11*), ependymal cells (*CA4*, *WLS*, *MGST1*, *PRTG*, *PLS3*, *RSPO2*, *FAM122B*, *WNT2B*, *CDO1*), GABAergic neurons (*RAB8A*, *PIGK*, *ZDHHC5*, *FZD3*, *CCM2*, *MRPL47*, *PNRC1*, *RHBDD2*, *CDC42SE1*), glutaminergic neurons (*MT-CYB*, *MT-ND5*, *NEFM*, *MT-ATP6*, *MT-ND4*, *NEFL*, *MTND3*, *MT-ND2*, *MT-CO3*), and neural progenitor cells (*RPS27*, *FTL*, *RPL21*, *FTH1*, *RPL15*, *RPL13A*, *RPS5*, *CDKN1A*, *VIM*) were also detected, along with radial glial cells (*PTN*, *C1orf61*, *VIM*, *CLU*, *HES1*, *HSPB1*, *MDK*, *DBI*, *LINC01158*) (Supplementary Table 7).

By applying UMAP analysis, we identified eight distinct cell population clusters (Figure 6A-D). The control cortical organoid consisted of 0.02% astrocytes, 14.18% DA GLU neurons, 58.67% dopaminergic neurons, 0.04% ependymal cells, 0.04% GABAergic neurons, 9.05% glutaminergic neurons, 1.60% neural progenitor cells, and 16.40% radial glial cells (Figure 6C and Supplementary Figure 4). Further classification of the neuronal populations revealed that 16.97% were DA GLU neurons, 70.23% were dopaminergic neurons, 0.05% were GABAergic neurons, 10.83% were glutaminergic neurons, and 1.92% were neural progenitor cells (Figure 6D).

**Figure 6:**
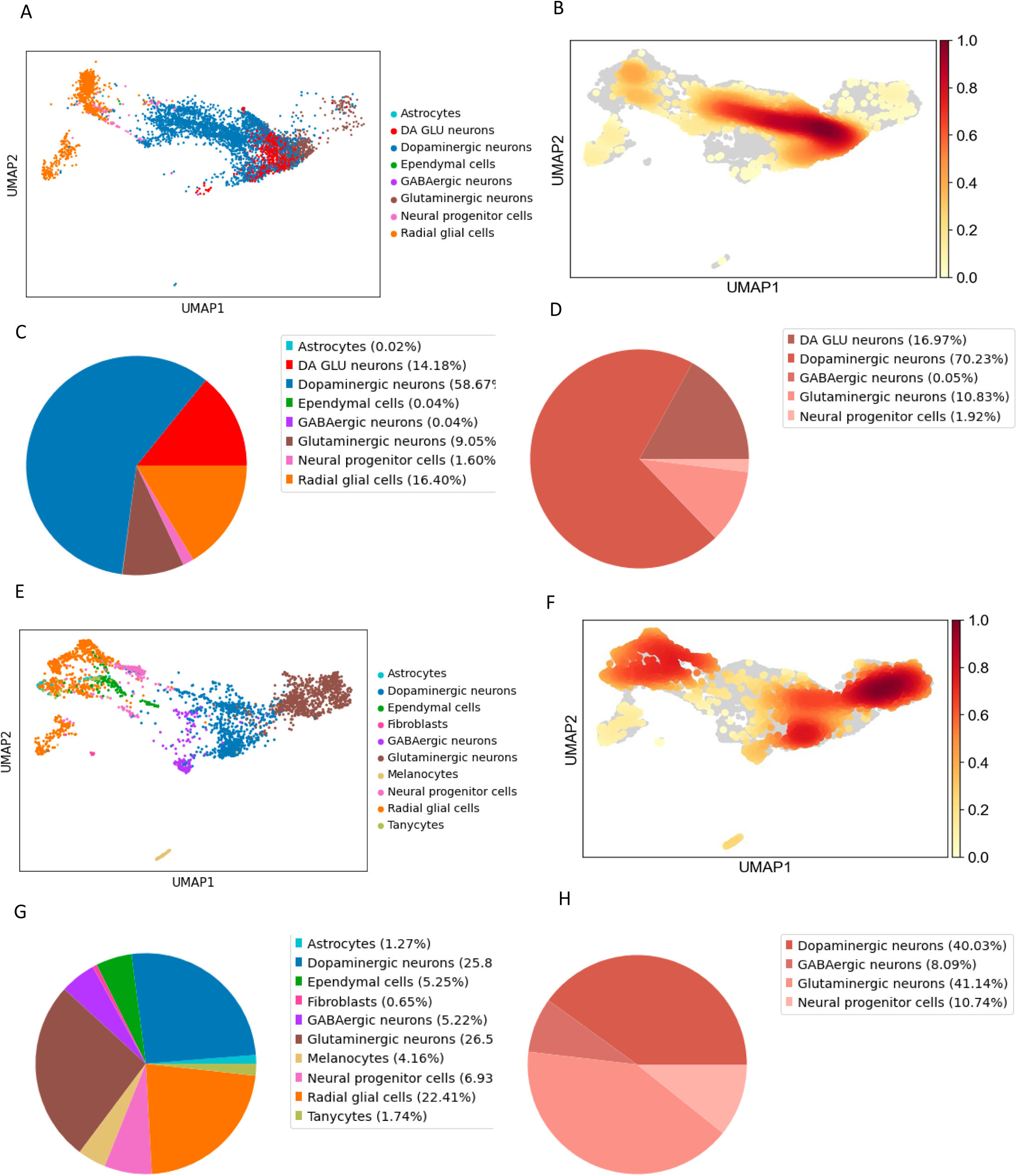
Comparison of Single-cell transcriptomic profiling in cortical organoids from control and patient iPSCs. (A, B) Cell clusters in 3-month-old organoids derived from control iPSCs were visualized using the scMRMA and UMAP algorithm (A), and their density is depicted in (B). (C, D) The percentage distribution of all cell clusters (C) and the neuro population (D) in organoids at 3-months old, which were derived from control iPSCs, were represented with the PCA algorithm. (E, F) Cell clusters in 3-month-old organoids derived from patient iPSCs were visualized using the scMRMA and UMAP algorithm (E), and their density is shown in (F). (G, H) The percentage distribution of all cell clusters (G) and the neuro population (H) in organoids at 3-months old, which were derived from patient iPSCs, were represented with the PCA algorithm.

Pathway enrichment analyses on the neuronal population (DA GLU neurons and other neurons) affirmed the enrichment of gene sets linked to neural development processes, including axon development, neural migration, generation of neurons, regulation of neural differentiation, neural tube development, axonogenesis, and synaptic transmission (Supplementary Figure 5 and 6, and Supplementary Table 8 and 9).

Our results suggest that human iPSC-derived cortical organoids comprise multiple neuronal cell types, encompassing mature neurons and astrocytes.

### Single-cell transcriptomics confirm neuronal loss and astrocytosis in POLG patient organoids and reveal alterations in multiple regulatory pathways

Next, we compared the single-cell transcriptomic profiles between the patient and control samples. For the patient group, cortical organoids derived from compound heterozygous CP2A were utilized. We analyzed a total of 3331 cells from the patient organoid, which expressed 25389 genes (Supplementary Table 6).

The patient cortical organoids of cell type enrichment analysis in scRNA-seq analysis could be subcategorized further into GABAergic neurons (*H3F3B*, *NREP*, *SOX4*, *NR2F2*, *MLLT11*, *EIF4G2, RTN, RBFOX2, GAD1*), glutaminergic neurons (*TCF7L2*, *GAP43, STMN2*, *MAP1B*, *STMN1*, *SCG2*, *NEFL*, *STMN4*, *PGM2L1*), dopaminergic neurons (*RTN1*, *NREP*, *STMN2*, *PTMA*, *NSG2*, *TUBA1A*, *NOVA1*, *MLLT11*, *TMSB10*) and neural progenitor cells (*RPS19*, *RPS27*, *EEF1A1*, *RPL37*, *RPL7*, *RPLP0*, *RPL18A*, *C1orf61*, *RPL13A*). Melanocytes (*MLANA*, *PMEL*, *TYRP1*, *ANXA2*, *GPNMB*, *DCT*, *SAT1*, *QPCT*, *LGALS1*) were also detected, along with radial glial cell (*C1orf61*, *PTN*, *GPM6B*, *VIM*, *CLU*, *PTPRZ1*, *CNN3*, *TTYH1*, *EDNRB*), along with tanycytes (*SPARCL1*, *GPM6B*, *CLU*, *PTN*, *NTRK2*, *ATP1A2*, *PSAT1*, *SLC1A3*, *SPARC*), along with fibroblasts (*LUM*, *COL3A1*, *DCN*, *LGALS1*, *COL5A1*, *RPS18*, *RPL37*, *RPL31*, *RPL13*), ependymal cells (*CLU*, *SPARCL1*, *IFITM3*, *SPARC*, *B2M*, *RPS27L*, *SERF2*, *PLTP*, *MGST1*) and astrocyte-related genes (*ATP1A2*, *NTRK2*, *SLC1A3*, *SPARCL1*, *GPM6B*, *PTN*, *ADGRG1*, *CLU*, *TTYH1*) (Figure 6G and H, and Supplementary Table 10).

Using the UMAP annotation plot, we identified ten distinct cell population clusters (Figure 6E-H), identified through the expression of specific markers (Supplementary Table 10). The patient cortical organoid comprised 1.27% astrocytes, 25.82% dopaminergic neurons, 5.25% ependymal cells, 0.65% fibroblasts, 5.22% GABAergic neurons, 26.54% glutaminergic neurons, 4.16% melanocytes, 6.93% neural progenitor cells, 22.41% radial glial cells, and 1.74% tanycytes (Figure 6E and Supplementary Figure 7). Further classification of the neuronal populations revealed 40.03% dopaminergic neurons, 8.09% GABAergic neurons, 41.14% glutaminergic neurons, and 10.74% neural progenitor cells (Figure 6H).

Patient organoids displayed neurons, radial glial cells, astrocytes, and ependymal cells, similar to controls. However, unique clusters in the patient organoids included melanocytes, tanycytes and fibroblasts (Figure 6E, G, H). When compared to controls, there was a decrease in the percentage of neurons (from 83.54 % in control to 64.51% in patient organoids), with an increase in astrocytes (from 0.02 % in control to 1.27 % in patient organoids) and radial glial cells (from 16.40 % in control to 22.41 % in patient organoids) (Figure 6G and H). A notable finding was the marked reduction of dopaminergic neurons and DA GLU neurons in patient organoids (Figure 6G and H).

Gene expression analysis of neuronal populations in patients and control organoids showed 206 DEGs in patients, with 78 being up-regulated and 128 down-regulated (Figure 7A). Particularly, several mtDNA-encoded genes, such as *MT-ND5, MT-RNR1, MT-RNR2, MT-TL1*, and *MT-TY*, were among the most significantly downregulated genes (Figure 7B). Enrichment analysis for the downregulated genes highlighted functions relating to neuronal differentiation, dendrite formation, iron metabolism, reactive oxygen species, amino acid metabolism, and MAPK signaling (Figure 7C-E, Table 1, 2, Supplementary Figure 8, and Supplementary Table 11). On the other hand, upregulated genes were enriched in pathways tied to neurodegenerative diseases, NOTCH and JAK-STAT signaling pathways (Figure 7F, Table 3).

**Figure 7:**
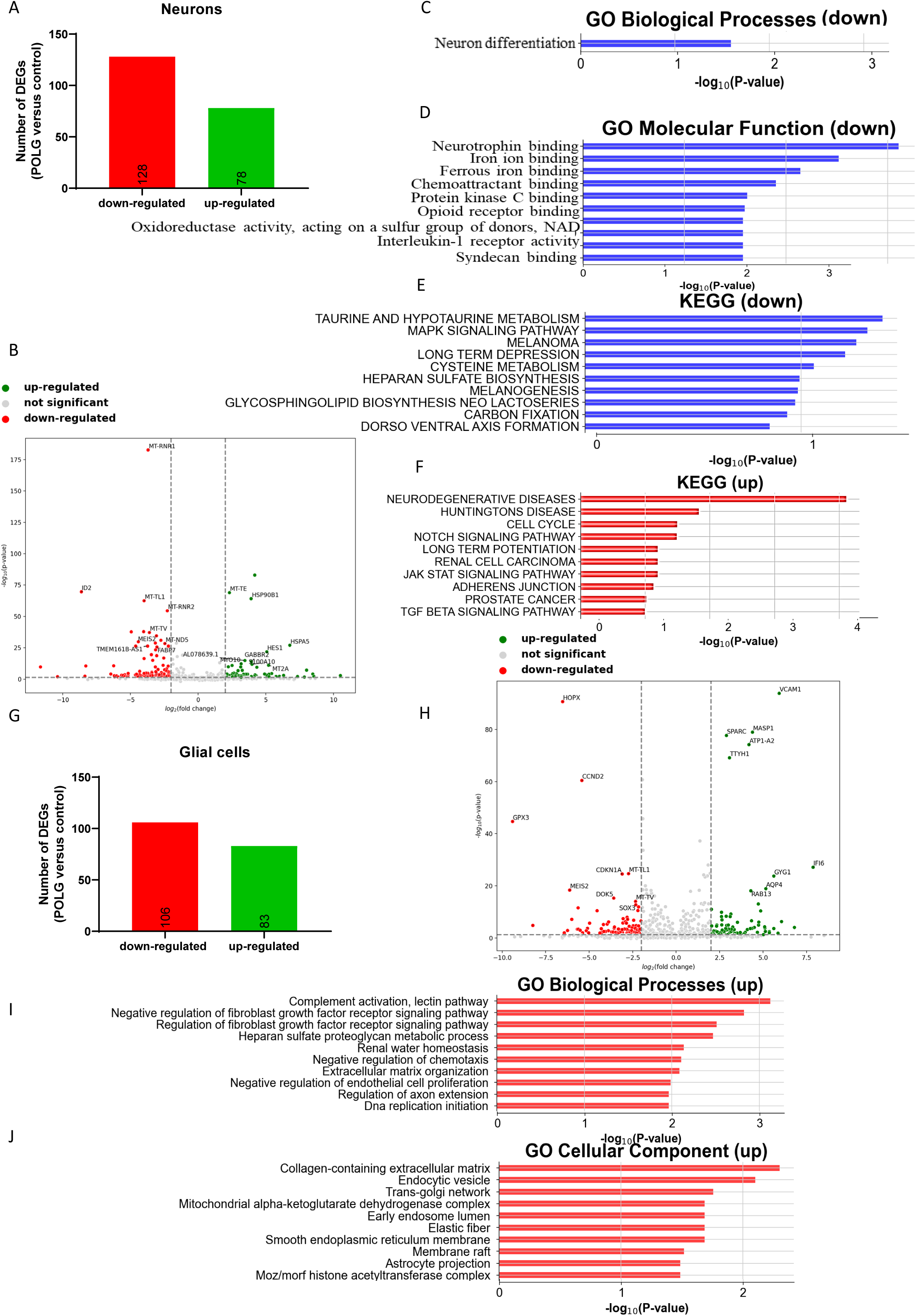
Comparison of molecular pathways in cortical organoids from control and patient iPSCs. (A) The number of DEGs that are upregulated and downregulated in neuron population of patient organoids compared to control organoids. (B) Volcano plot illustrating the DEGs in neuron population of patient organoids compared to control organoids. (C) GO biological progresses enriched for downregulated DEGs in neuron population of patient organoids compared to control organoids. (D) GO molecular functions enriched for downregulated DEGs in the neuron population of patient organoids compared to control organoids. (E) KEGG pathways enriched for downregulated DEGs in neuron population of patient organoids compared to control organoids. (F) KEGG pathways enriched for upregulated DEGs in neuron population of patient organoids compared to control organoids. (G) The number of DEGs that are upregulated and downregulated in glial cell population (astrocytes and radial glial cells) of patient organoids compared to control organoids. (H) Volcano plot illustrating the DEGs in glial cell population (astrocytes and radial glial cells) in patient organoids and control organoids. (I) GO biological processes enriched for upregulated DEGs in glial cell population (astrocytes and radial glial cells) in patient organoids compared to control organoids. (J) GO cellular components enriched for upregulated DEGs in glial cell population (astrocytes and radial glial cells) in patient organoids compared to control organoids.

**Table 1:**
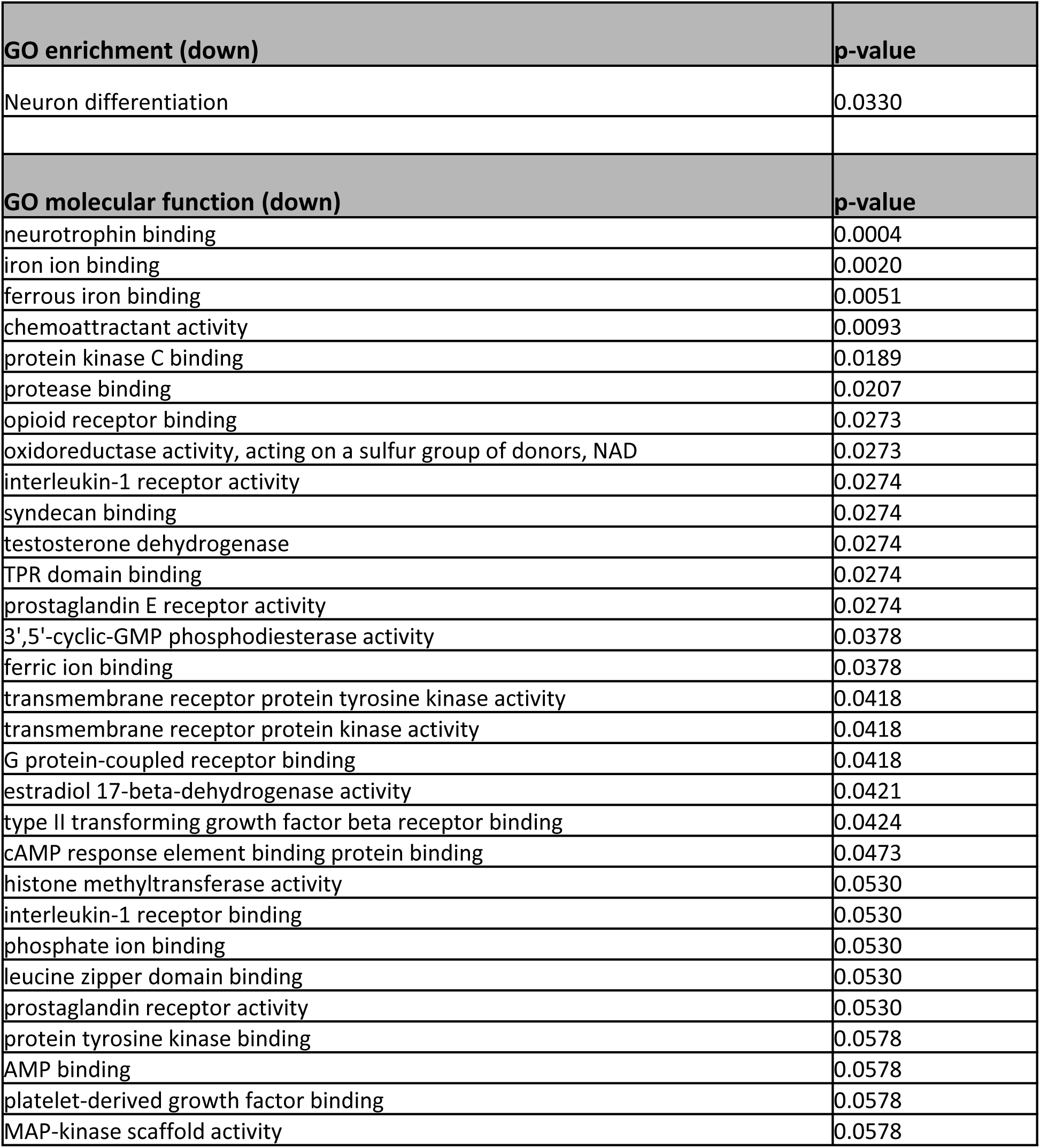
GO enrichment by down-regulated DEGs in neuron population of patient organoids versus controls.

| **GO enrichment (down)** | **p-value** |
| --- | --- |
| Neuron differentiation | 0.0330 |
| **GO molecular function (down)** | **p-value** |
| neurotrophin binding | 0.0004 |
| iron ion binding | 0.0020 |
| ferrous iron binding | 0.0051 |
| chemoattractant activity | 0.0093 |
| protein kinase C binding | 0.0189 |
| protease binding | 0.0207 |
| opioid receptor binding | 0.0273 |
| oxidoreductase activity, acting on a sulfur group of donors, NAD | 0.0273 |
| interleukin-1 receptor activity | 0.0274 |
| syndecan binding | 0.0274 |
| testosterone dehydrogenase | 0.0274 |
| TPR domain binding | 0.0274 |
| prostaglandin E receptor activity | 0.0274 |
| 3',5'-cyclic-GMP phosphodiesterase activity | 0.0378 |
| ferric ion binding | 0.0378 |
| transmembrane receptor protein tyrosine kinase activity | 0.0418 |
| transmembrane receptor protein kinase activity | 0.0418 |
| G protein-coupled receptor binding | 0.0418 |
| estradiol 17-beta-dehydrogenase activity | 0.0421 |
| type II transforming growth factor beta receptor binding | 0.0424 |
| cAMP response element binding protein binding | 0.0473 |
| histone methyltransferase activity | 0.0530 |
| interleukin-1 receptor binding | 0.0530 |
| phosphate ion binding | 0.0530 |
| leucine zipper domain binding | 0.0530 |
| prostaglandin receptor activity | 0.0530 |
| protein tyrosine kinase binding | 0.0578 |
| AMP binding | 0.0578 |
| platelet-derived growth factor binding | 0.0578 |
| MAP-kinase scaffold activity | 0.0578 |

**Table 2:** KEGG pathways enriched by down-regulated DEGs in neuron population of patient organoids versus controls.

| KEGG (down) | p-value |
| --- | --- |
| TAURINE AND HYPOTAURINE METABOLISM | 0.0434 |
| MAPK SIGNALING PATHWAY | 0.0506 |
| MELANOMA | 0.0573 |
| LONG TERM DEPRESSION | 0.0649 |
| CYSTEINE METABOLISM | 0.0900 |
| HEPARAN SULFATE BIOSYNTHESIS | 0.1039 |
| MELANOGENESIS | 0.1069 |
| GLYCOPHINGOLIPID BIOSYNTHESIS NEO LACTOSERIES | 0.1080 |
| CARBON FIXATION | 0.1188 |
| DORSO VENTRAL AXIS FORMATION | 0.1399 |
| THYROID CANCER | 0.1441 |
| CYTOKINE CYTOKINE RECEPTOR INTERACTION | 0.1590 |
| GLUTATHIONE METABOLISM | 0.1897 |
| BLADDER CANCER | 0.2022 |
| LYSINE DEGRADATION | 0.2238 |
| ENDOMETRIAL CANCER | 0.2441 |
| ACUTE MYELOID LEUKEMIA | 0.2484 |
| ANDROGEN AND ESTROGEN METABOLISM | 0.2527 |
| ARACHIDONIC ACID METABOLISM | 0.2505 |
| NON SMALL CELL LUNG CANCER | 0.2520 |
| GLYCAN STRUCTURES BIOSYNTHESIS 2 | 0.2838 |
| FOCAL ADHESION | 0.2921 |
| GLYCOLYSIS AND GLUCONEOGENESIS | 0.2921 |
| GLIOMA | 0.2921 |
| LONG TERM POTENTIATION | 0.3096 |
| RENAL CELL CARCINOMA | 0.3096 |
| VEGF SIGNALING PATHWAY | 0.3150 |
| PANCREATIC CANCER | 0.3252 |
| FC EPSILON RI SIGNALING PATHWAY | 0.3338 |
| CHRONIC MYELOID LEUKEMIA | 0.3377 |

**Table 3:**
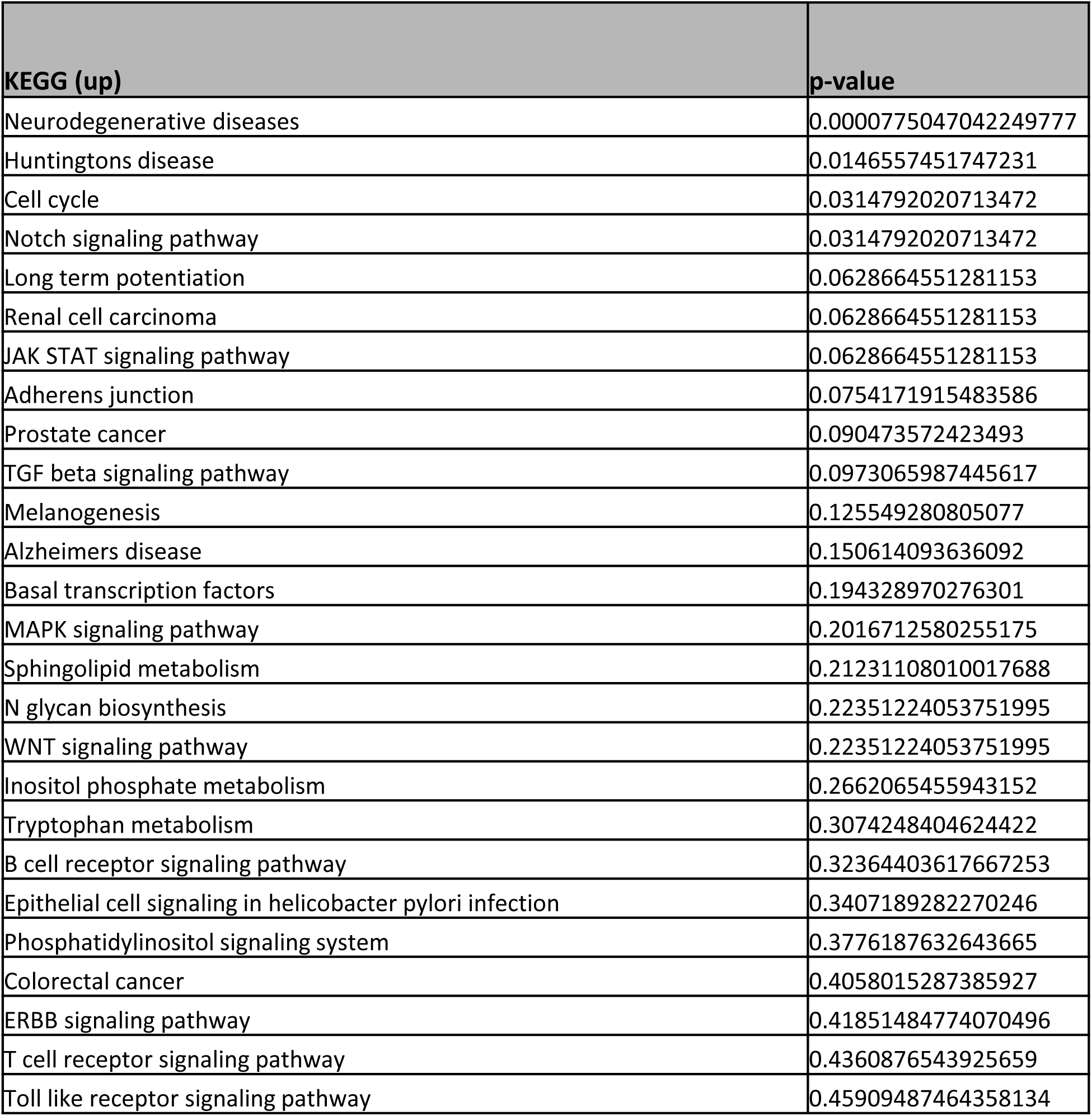
KEGG pathways enriched by up-regulated DEGs in neuron population of patient organoids versus controls.

Looking at the glial cell population, consisting of radial glial cells and astrocytes, we found 189 DEGs when compared to the control organoids, with 83 genes upregulated and 106 genes downregulated in the patient organoids’ glial population (Figure 6G and H). GO analysis on the regulated genes in this population revealed an enrichment in processes including complement activation pathway and astrocyte projection in the patient samples as opposed to controls (Figure 6I and J, Table 4). Conversely, downregulated genes in glial cells were enriched in processes like central nervous system development, axon guidance and nervous system development (Supplementary Figure 9, Supplementary Table 12).

**Table 4:**
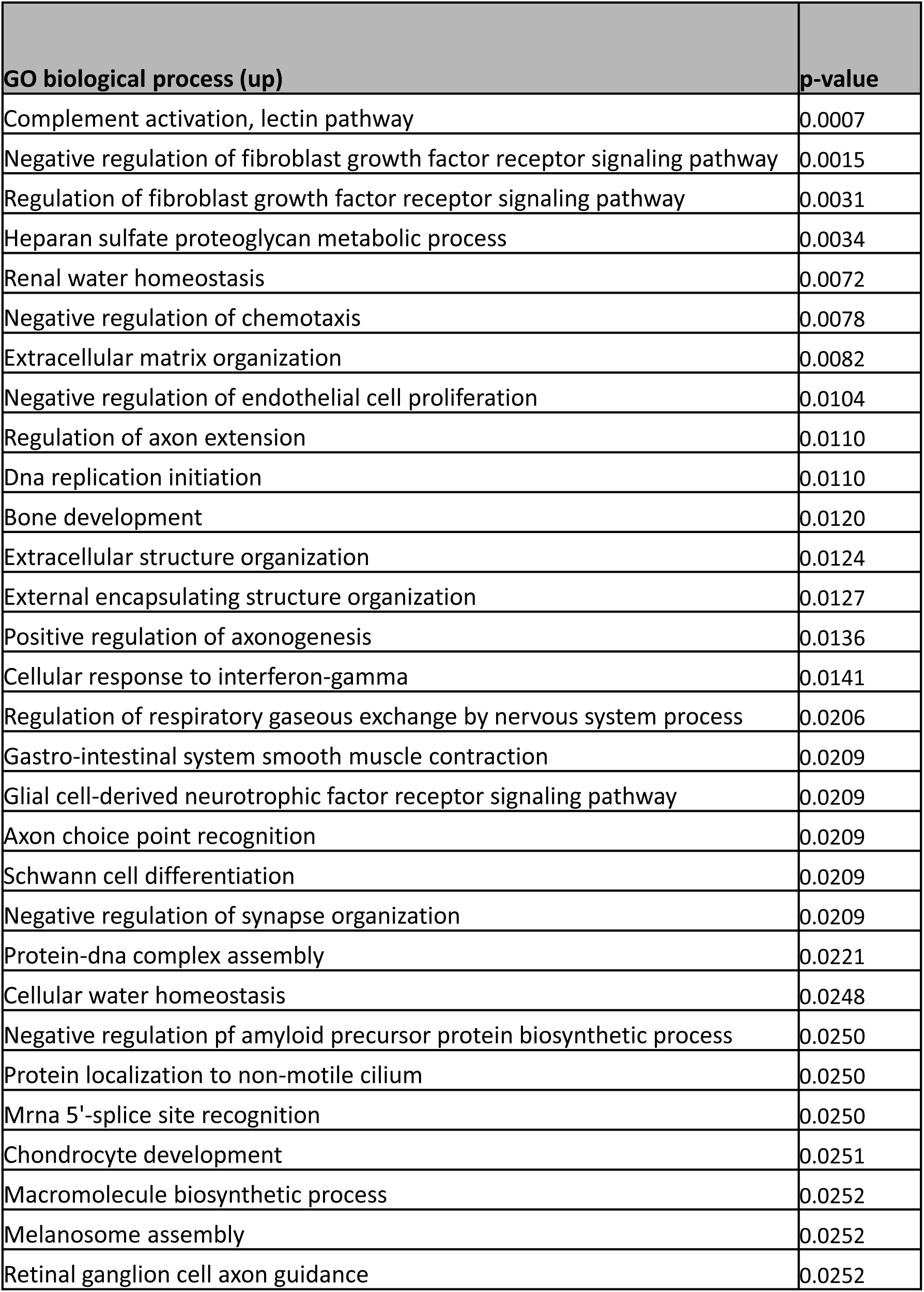
GO enrichment and molecular functions of up-regulated DEGs in glial cell population of patient organoids versus controls.

| GO biological process (up) | p-value |
| --- | --- |
| Complement activation, lectin pathway | 0.0007 |
| Negative regulation of fibroblast growth factor receptor signaling pathway | 0.0015 |
| Regulation of fibroblast growth factor receptor signaling pathway | 0.0031 |
| Heparan sulfate proteoglycan metabolic process | 0.0034 |
| Renal water homeostasis | 0.0072 |
| Negative regulation of chemotaxis | 0.0078 |
| Extracellular matrix organization | 0.0082 |
| Negative regulation of endothelial cell proliferation | 0.0104 |
| Regulation of axon extension | 0.0110 |
| Dna replication initiation | 0.0110 |
| Bone development | 0.0120 |
| Extracellular structure organization | 0.0124 |
| External encapsulating structure organization | 0.0127 |
| Positive regulation of axonogenesis | 0.0136 |
| Cellular response to interferon-gamma | 0.0141 |
| Regulation of respiratory gaseous exchange by nervous system process | 0.0206 |
| Gastro-intestinal system smooth muscle contraction | 0.0209 |
| Glial cell-derived neurotrophic factor receptor signaling pathway | 0.0209 |
| Axon choice point recognition | 0.0209 |
| Schwann cell differentiation | 0.0209 |
| Negative regulation of synapse organization | 0.0209 |
| Protein-dna complex assembly | 0.0221 |
| Cellular water homeostasis | 0.0248 |
| Negative regulation pf amyloid precursor protein biosynthetic process | 0.0250 |
| Protein localization to non-motile cilium | 0.0250 |
| Mrna 5'-splice site recognition | 0.0250 |
| Chondrocyte development | 0.0251 |
| Macromolecule biosynthetic process | 0.0252 |
| Melanosome assembly | 0.0252 |
| Retinal ganglion cell axon guidance | 0.0252 |

**Table 5:** GO cellular component of up-regulated DEGs in glial cell population of patient organoids versus controls.

| **GO cellular component (up)** | **p-value** |
| --- | --- |
| Collagen-containing extracellular matrix | 0.0050 |
| Endocytic vesicle | 0.0080 |
| Trans-golgi network | 0.0175 |
| Mitochondrial alpha-ketoglutarate dehydrogenase complex | 0.0205 |
| Early endosome lumen | 0.0205 |
| Elastic fiber | 0.0207 |
| Smooth endoplasmic reticulum membrane | 0.0207 |
| Membrane raft | 0.0304 |
| Astrocyte projection | 0.0330 |
| Moz/morf histone acetyltransferase complex | 0.0330 |
| Insulin-responsive compartment | 0.0367 |
| H3 histone acetyltransferase complex | 0.0367 |
| Multimeric ribonuclease p complex | 0.0411 |
| Sodium:potassium-exchanging atpase complex | 0.0411 |
| Transcription factor tfiih core complex | 0.0411 |
| Microfibril | 0.0446 |
| Melanosome membrane | 0.0489 |
| Chitosome | 0.0489 |
| Pigment granule membrane | 0.0489 |
| Transcription factor tfiih holo complex | 0.0489 |
| Lysosomal lumen | 0.0498 |
| Beta-catenin-tcf complex | 0.0530 |
| Podosome | 0.0533 |
| Mhc class ii protein complex | 0.0533 |
| Prespliceosome | 0.0609 |
| Trans-golgi network transport vesicle | 0.0615 |
| U2-type spliceosome | 0.0615 |
| Cation-transporting atpase complex | 0.0650 |
| Platelet alpha granule membrane | 0.0684 |
| U1 snrnp | 0.0723 |

In the cohort of dopaminergic neurons, analysis revealed 115 DEGs that were upregulated and 29 that were downregulated in patient cortical organoids compared to those that were control (Supplementary Figure 10). An enrichment analysis of the downregulated DEGs within this dopaminergic neuron population showed an association with several GO molecular function processes and cellular components. Notably, these included NAD+ ADP-ribosyltransferase activity within molecular functions, while cellular components encompassed structures such as the apical dendrite, dendrite, microtubule and cytoskeleton, spindle and spindle microtubule, asymmetric synapse, postsynaptic density, cytoskeleton, and axon (Supplementary Figure 11, 12 and Supplementary Table 13, 14). A KEGG pathway analysis of the downregulated DEGs within this dopaminergic neuron population showed enriched WNT, JAK STAT and MARK signaling pathways (Supplementary Figure 13).

Overall, these findings demonstrate substantial differences in gene expression between patient and control organoids, revealing significant changes in key neuronal and glial cell populations. The dysregulation of numerous genes related to neuronal differentiation, central nervous system development signaling pathways, in conjunction with shifts in the composition of specific neuronal cell types, underscores the complex genetic and cellular landscape of POLG-related disease.

### Metformin ameliorates the phenotype of the POLG cortical organoid

In the realm of therapeutic interventions for POLG-associated disorders, the role of metformin as a treatment option has garnered considerable attention. Building on our previous work, we’ve substantiated its effectiveness through an array of tests [28]. A closer examination of metformin’s potential therapeutic pathways reveals its intimate association with mitochondrial operations and cellular metabolic processes. This connection, in the milieu of POLG-related diseases, might be instrumental in delivering its curative properties. We next explored the potential of our organoid as a platform for preclinical drug trials. According to prior studies, metformin could alleviate neuronal damage/loss, encourage neuronal differentiation, and suppress astrocyte formation in a cerebral ischemia/reperfusion rat model [35]. We, along with others, have found that metformin fosters mitochondrial biogenesis and bolsters mitochondrial activity and function [28, 36]. Thus, we asked if metformin could mitigate the phenotype and molecular profile of the POLG-organoid. We exposed patient organoids to 250 µM metformin from day 6 and observed an enhancement in cortical organoid structure and morphology after a 30-day (Figure 8A). Moreover, metformin-treated organoids showed reduced expression of astrocytic marker GFAP (Figure 8B and D) and elevated levels of neuronal marker TUJ1 (Figure 8B and E), compared to their untreated counterparts, while the cortical neural marker SATB2 remained stable (Figure 8B and C). We also noted an upswing in both synaptophysin (Figure 8F and G), and PSD-95 expressions (Figure 8F and H) compared to the untreated samples.

**Figure 8:**
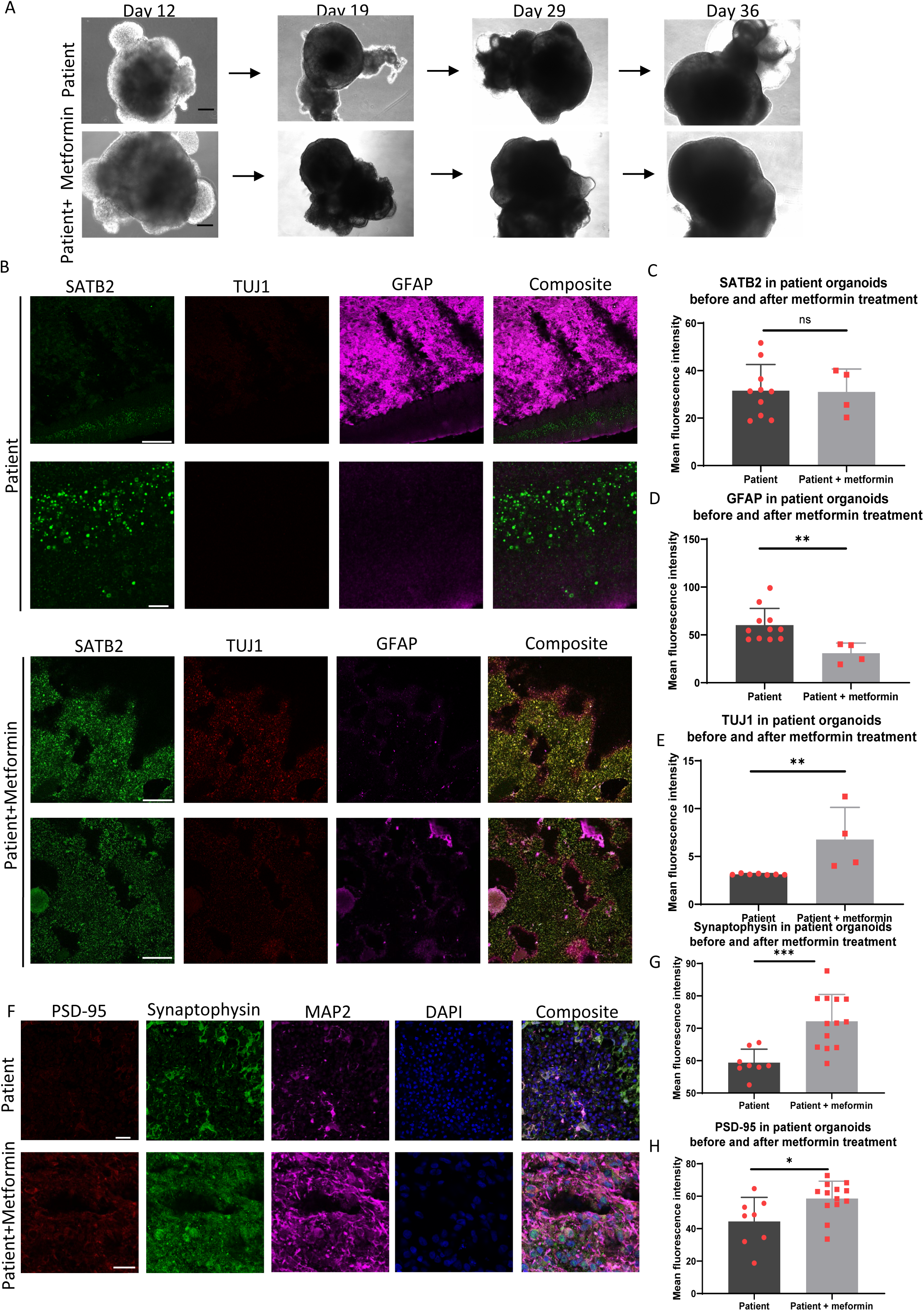
Comparison of lineage markers in patient brain organoid before and after metformin treatment. (A) Representative phase contrast images of the cell morphology of patient brain organoids before and after metformin treatment at day 12, 19, 29, and 36. Black scale bar is 100 µm. (B) Immunofluorescent imaging of cryo-sectioned organoids at 36 days, staining the cortical pyramidal neuronal marker SATB2 (green), newborn neural marker TUJ1 (green), and reactivated astrocyte marker GFAP (red) in patient organoids and patient organoids treated with metformin. Scale bar is 100 µm. (C-E) Quantitative measurements of the level of SATB2 (D), GFAP (E), and TUJ1 (F) expression on day 36 in patient organoids and patient organoids treated with metformin. The Y-axis represents the mean fluorescence intensity. (F) Immunofluorescent imaging of cryo-sectioned organoids, staining the presynaptic marker Synaptophysin (green), postsynaptic marker PSD-95 (red), and mature neural marker MAP2 (purple) in control and patient organoids. Nuclei are stained with DAPI (blue). Scale bar is 100 µm. (G-H) Quantitative measurements of the level of Synaptophysin (G) and PSD-95 (H) expression in control and patient cortical organoids. The Y-axis represents the mean fluorescence intensity. Significance levels are indicated for P values less than 0.05, with * indicating P < 0.05, ** indicating P < 0.01, *** indicating P < 0.001, **** indicating P < 0.000. Results not reaching statistical significance are denoted as ns (not significant).

When we stained complex I (NDUFB10), TOMM20 and MAP2, we detected a marked increase in the levels of TOMM20 (Figure 9A and B), MAP2 (Figure 9A, E and G, Supplementary Figure 14) and total complex I (Figure 9A and C, Supplementary Figure 14) in metformin treated samples compared to untreated ones. On normalizing total complex I (NDUFB10) to the mitochondrial mass (TOMM20), we saw a significantly higher level of NDUFB10/TOMM20 in metformin treated patient organoids (Figure 9A and D). Further, immunostaining of TFAM, SOX2 and MAP2 showed significantly increased levels of TFAM (Figure 9E and F) and MAP2 expression (Figure 9E and G) in the patient organoids suggesting an increase in mtDNA copy number and neural regrowth with metformin treatment compared to untreated samples.

**Figure 9:**
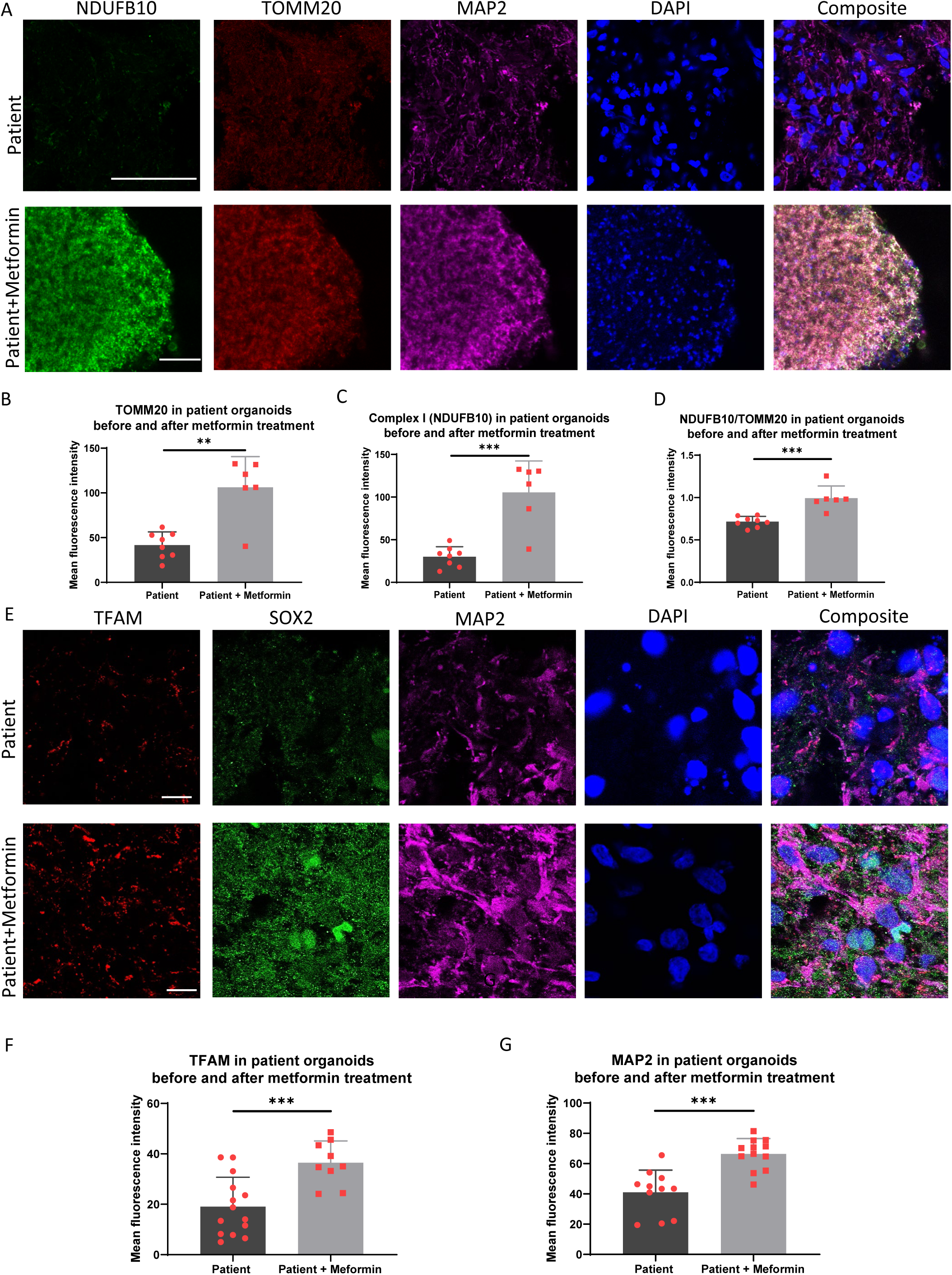
metformin rescues the neuronal damage and mitochondrial defect in POLG patient brain organoid. (A) Immunofluorescent imaging of cryo-sectioned organoids at 90 days, showing the staining of NDUFB10 (green), TOMM20 (red), and MAP2 (purple) in patient organoids and patient organoids treated with metformin. Nuclei are stained with DAPI (blue). Scale bar is 100 µm. (B-D) Quantitative measurements of the level of TOMM20 (B), NDUFB10 (C), and NDUFB10/TOMM20 (D) expression. The Y-axis represents the mean fluorescence intensity. (E) Immunofluorescent imaging of cryo-sectioned organoids at 36 days, staining TFAM (red), SOX2 (green), and MAP2 (purple) in patient organoids and patient organoids treated with metformin. Nuclei are stained with DAPI (blue). Scale bar is 100 µm. (F, G) Quantitative measurements of the level of TFAM (F) and MAP2 (G). The Y-axis represents the mean fluorescence intensity. Significance levels are indicated for P values less than 0.05, with * indicating P < 0.05, ** indicating P < 0.01, *** indicating P < 0.001.

In summary, metformin treatment of POLG organoids led to improved structure and morphology, an increased neuronal marker and reduced astrocytic markers, increased mitochondrial mass and complex I and TFAM expression. These results demonstrate metformin’s potential for mitigating the phenotype and enhancing the molecular profile of POLG organoids, validating the utility of organoid models in preclinical drug trials.

### Single-cell transcriptomics confirm neuronal enrichment and decreased astrocytosis in POLG organoids upon metformin treatment

We next evaluated the impact of metformin on the single-cell transcriptomic profile of POLG organoids by treating the CP2A patient organoids for over 2 months. We identified a total of 8433 cells expressing 26,403 genes (Supplementary Table 6). The patient cortical organoids upon metformin treatment of cell type enrichment analysis in scRNA-seq analysis could be subcategorized further into dopaminergic neurons (MAB21L1, *PTMA*, *RTN1*, *TMSB10*, *MLLT11*, *NREP*, *STMN2*, *H3F3B*, *VSNL1*), ependymal cells (*CLU*, *IFITM3*, *SPARC*, *SPARCL1*, *B2M*, *NPC2*, *SERF2*, *GSTP1*, *CRYAB*), along with fibroblasts (*COL3A1*, *COL1A2*, *COL1A1*, *LGALS1*, *ISLR*, *COL6A3*, *LUM*, *MFAP4*, *POSTN*), GABAergic neurons (*SOX4*, *LHX1*, *NREP*, *ZNF385D*, *H3F3B*, *TFAP2A*, *RTN1*, *BLCAP*, *ETFB*), glutaminergic neurons (*TCF7L2*, *NEFL*, *STMN2*, *GAP43*, *MAP1B*, *STMN1*, *LHX9*, *NEFM*, *WLS*), melanocytes (*TYR*, *TYRP1*, *DCT*, *MLANA*, *PMEL*, *LGALS3*, *PTGDS*, *SAT1*, *CTSB*) and neural progenitor cells (*RPS27L*, *RPS27*, *RPS19*, *FTL*, *GNG5*, *RPLP1*, *RPL7*, *VIM*, *RPS3A*) were also detected, along with radial glial cells (*PTN*, *C1orf61*, *VIM*, *GPM6B*, *GNG5*, *DBI*, *SPARC*, *CNN3*, *TTYH1*), along with tanycytes (*SPARCL1*, *ATP1A2*, *PTN*, *CLU*, *NTRK2*, *MGST1*, *IFITM3*, *PLTP*, *VIM*), and astrocyte-related genes (*ATP1A2*, *SPARCL1*, *SLC1A3*, *PI15*, *QKI*, *PTPRZ1*, *GPM6B*, *PTN*, *NTRK2*) (Figure 10A-D).

**Figure 10:**
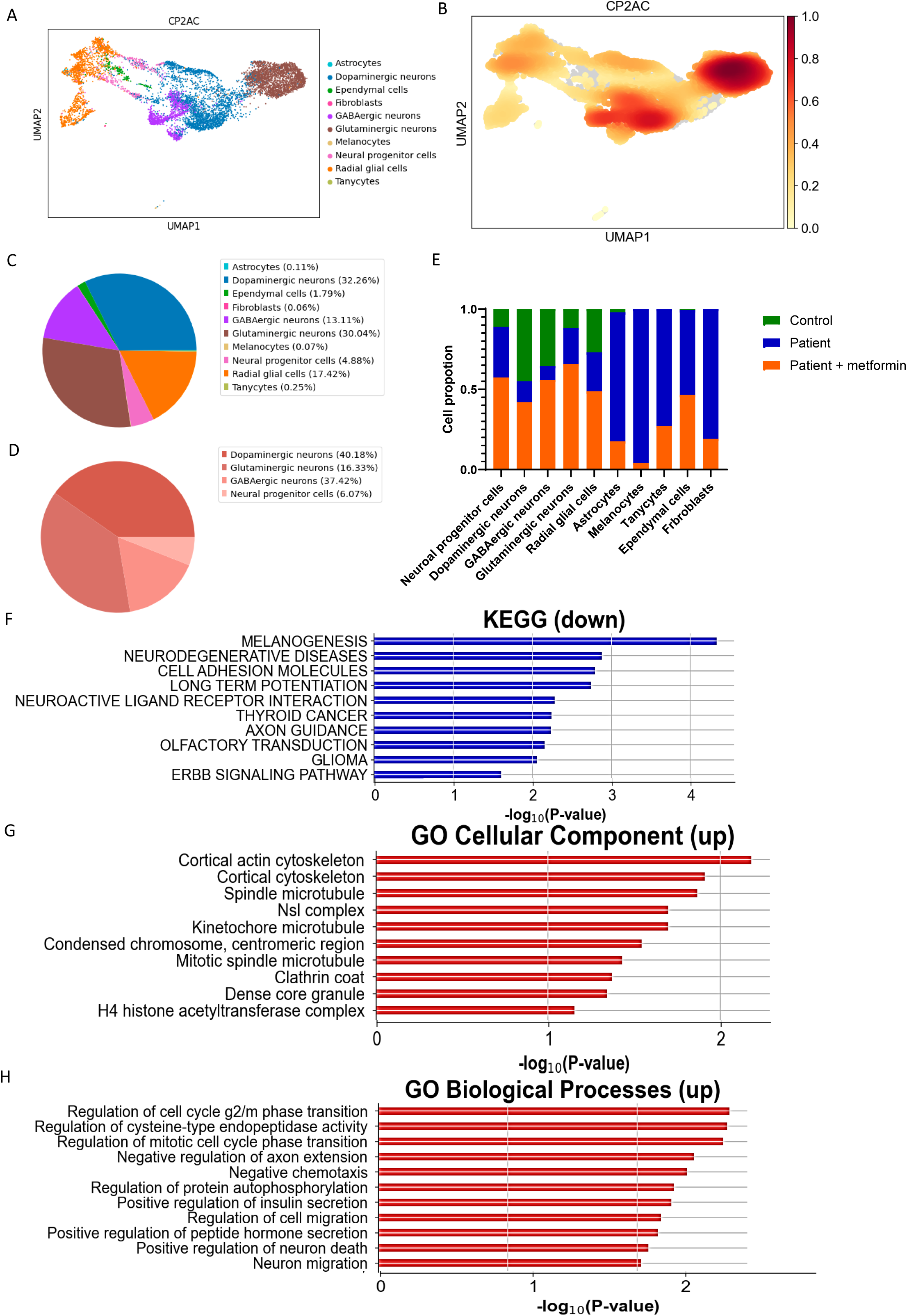
Comparison of molecular pathways in cortical organoids from patient brain organoid before and after metformin treatment. (A, B) Cell clusters in 3-month-old organoids derived from patient iPSCs with metformin treatment were visualized using the scMRMA and UMAP algorithm (A), and their density is depicted in (B). (C, D) The percentage distribution of all cell clusters (C) and the neuro population (D) in patient organoids with metformin treatment, were represented with the PCA algorithm. (E) Percentage histogram showing the distribution of individual cell clusters in control organoids, patient organoids, and patient organoids treated with metformin. (F) KEGG pathways enriched for down-regulated differentially expressed genes (DEGs) in the neuron population of patient organoids compared to patient organoids treated with metformin. (G) GO cellular components enriched for up-regulated DEGs in the neuron population of patient organoids compared to patient organoids treated with metformin. (H) GO biological processes enriched for up-regulated DEGs in the neuron population of patient organoids compared to patient organoids treated with metformin.

The UMAP plot identified eight distinct cell clusters, which included neurons (80.29%), two subtypes of radial glial cells (17.42%), astrocytes (0.11%), ependymal cells (1.79%), melanocytes (0.07%), tanycytes (0.25%) and fibroblasts (0.06%) (Figure 10 A-D and Supplementary Figure 15). Interestingly, even after metformin treatment, we did not detect a DA GLU neuron population in the POLG organoids, indicating that metformin may not alleviate their absence (Figure 10A-D, and Supplementary Table 15).

Further comparison of cell cluster proportions between control organoids, untreated POLG organoids, and metformin-treated POLG organoids indicated shifts in cell populations. Metformin treatment appeared to reduce the astrocyte population while increasing the neuron population when we compared untreated and treated POLG organoids (Figure 10E). Furthermore, metformin seemed to decrease the presence of melanocytes, tanycytes, and fibroblasts (Figure 10E).

Pathway analysis in the neuronal population revealed downregulated genes were enriched for multiple pathways, including those associated with neurodegenerative diseases and axon guidance (Figure 10E, Table 6). On the other hand, upregulated genes were enriched for processes related to cortical actin cytoskeleton, cortical cytoskeleton, neuronal migration, and regulation of neurotransmitter transport (Figure 10F and G, Table 7 and 8, Supplementary Figure 16). Similar patterns were observed in glial cells, where downregulated genes showed enrichment for pathways related to neurodegenerative diseases (Supplementary Figure 17 and Supplementary Table 17).

**Table 6:**
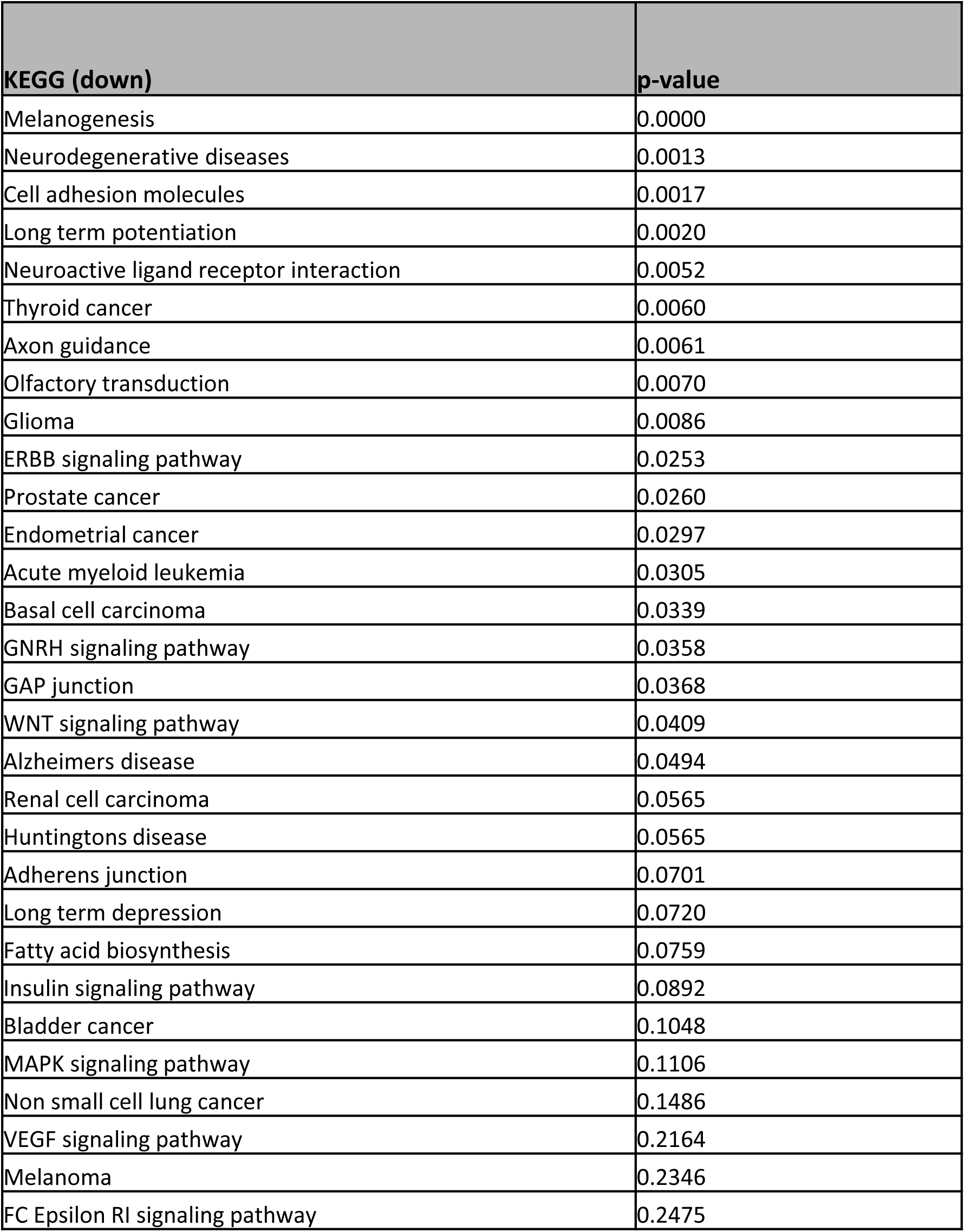
KEGG pathways of down-regulated DEGs in the neuron population in treated patient organoids versus untreated group.

**Table 7:** GO cellular component of up-regulated DEGs in the neuron population in treated patient organoids versus untreated group.

| **GO cellular component (up)** | **p-value** |
| --- | --- |
| Cortical actin cytoskeleton | 0.0065285211411632085 |
| Cortical cytoskeleton | 0.0122016834288158 |
| Spindle microtubule | 0.01347812964903343 |
| Nsl complex | 0.019783188827670047 |
| Kinetochore microtubule | 0.019783188827670047 |
| Condensed chromosome, centromeric region | 0.028627941656276728 |
| Mitotic spindle microtubule | 0.03750369637859683 |
| Clathrin coat | 0.042621588289367775 |
| Dense core granule | 0.0444782967604326 |
| H4 histone acetyltransferase complex | 0.06716770257616217 |
| Nuclear inner membrane | 0.07853472518961592 |
| Condensed chromosome | 0.14065272420579772 |
| Tertiary granule lumen | 0.14677992675719528 |
| Caveola | 0.1575908014459229 |
| Specific granule lumen | 0.16445587118522992 |
| Clathrin-coated endocytic vesicle membrane | 0.17909621999960726 |
| Cell-cell junction | 0.18165997883083781 |
| Voltage-gated potassium channel complex | 0.1868981223255205 |
| Endoplasmic reticulum lumen | 0.19783188827091158 |
| Lipid droplet | 0.20353634681624722 |
| Potassium channel complex | 0.20644996557731154 |
| Plasma membrane raft | 0.209405292733057 |
| Clathrin-coated endocytic vesicle | 0.21544346899478176 |
| Clathrin-coated vesicle membrane | 0.2248287530720051 |
| Azurophil granule lumen | 0.22804717220560994 |
| Actin cytoskeleton | 0.23131166294517203 |
| Secretory granule lumen | 0.23131166294517203 |
| Intracellular non-membrane-bounded organelle | 0.24138820728065213 |
| Integral component of plasma membrane | 0.248348607010951 |
| Microtubule cytoskeleton | 0.2448436746722709 |

**Table 8:** GO biological processes of up-regulated DEGs in the neuron population in treated patient organoids versus untreated group.

| **GO biological processes (up)** | **p-value** |
| --- | --- |
| Regulation of cell cycle g2/m phase transition | 0.0020 |
| Regulation of cysteine-type endopeptidase activity | 0.0021 |
| Regulation of mitotic cell cycle phase transition | 0.0022 |
| Negative regulation of axon extension | 0.0036 |
| Negative chemotaxis | 0.0042 |
| Regulation of protein autophosphorylation | 0.0052 |
| Positive regulation of insulin secretion | 0.0056 |
| Regulation of cell migration | 0.0066 |
| Positive regulation of peptide hormone secretion | 0.0071 |
| Positive regulation of neuron death | 0.0081 |
| Neuron migration | 0.0094 |
| Regulation of g2/m transition of mitotic cell cycle | 0.0094 |
| Regulation of cytokine production | 0.0100 |
| Heart morphogenesis | 0.0106 |
| Negative regulation of viral genome replication | 0.0106 |
| Positive regulation of interleukin-1 beta production | 0.0119 |
| Positive regulation of interleukin-1 production | 0.0139 |
| Negative regulation of endopeptidase activity | 0.0139 |
| Negative regulation of metallopeptidase activity | 0.0139 |
| Cellular response to heparin | 0.0147 |
| Lipid transport across blood-brain barrier | 0.0147 |
| Positive regulation of amino acid transport | 0.0150 |
| Cellular response to type I interferon | 0.0150 |
| Type I interferon signaling pathway | 0.0150 |
| Regulation of viral genome replication | 0.0166 |
| Histamine metabolic process | 0.0172 |
| Negative regulation of amino acid transport | 0.0172 |
| Regulation of macromolecule biosynthetic process | 0.0176 |
| Regulation of hormone biosynthetic process | 0.0176 |
| Positive regulation of tau-protein kinase activity | 0.0176 |

In the population of dopaminergic neurons, a total of 39 DEGs were identified as upregulated, while 251 were downregulated in cortical organoids from patients treated with metformin, as compared to untreated organoids (Supplementary Figure 17). When analyzing the upregulated DEGs within the Dopaminergic neuron population, several GO cellular components were found to be enriched. These included the integral and intrinsic component of mitochondrial outer membrane, integral component of mitochondrial membrane, mitochondrial outer membrane and synaptic membrane (Supplementary Figure 19 and Supplementary Table 18). KEGG analysis showed the upregulation of MARK and MTOR signaling pathways (Supplementary Figure 20).

Our single-cell transcriptomic analysis of metformin treated POLG organoids revealed changes in the proportions of different cell types and shifts in gene expression profiles, particularly in neuronal and glial populations, suggesting that metformin can influence neurodegenerative disease pathways and cellular components.

## Discussion

In our study, we generated a detailed cellular and molecular profile of cortical organoids developed from a patient harboring known pathogenic *POLG* mutations. The organoids recapitulated key aspects of the disease process, providing insights into disease mechanisms and a robust platform for drug testing. Immunostaining and scRNA-seq in our novel, human derived organoid model of POLG encephalopathy confirmed the presence of multiple types of neuronal and glial cells, consistent with differentiation into brain-like tissue. The scRNA-seq data also indicated the presence of radial glial cells, however, indicating that the maturation of the organoids is not complete. This is a common technical limitation of brain organoid research [15, 37]. Nevertheless, our model reliably recapitulated important hallmark tissue and molecular changes we have observed in our postmortem studies of patients with POLG encephalopathy, including neuronal loss affecting both dopaminergic and DA GLU neuron populations, astrocytosis, complex I deficiency and mtDNA depletion [38].

Our results demonstrated successful generation of cortical organoids from iPSCs using a neural induction protocol involving dual SMAD inhibition and canonical Wnt inhibition. The organoids exhibited progressive development, with the appearance of neural identity markers and the formation of cortical brain regions. At different stages of development, we observed distinct characteristics in the organoids, including the expression of neural markers and the presence of different neuronal cell types. By day 90, the organoids consisted predominantly of mature neurons positive for the marker NeuN, along with smaller populations of cells expressing GFAP, MAP2, and SOX2. Furthermore, the cortical pyramidal neuronal markers SATB2 and CTIP2 were observed in specific layers of the organoids, and stratified expressions of oligodendrocyte marker OLIG2, astrocyte marker GFAP, and neural marker DCX were detected. Importantly, the patient-derived organoids exhibited significant structural alterations compared to control organoids, characterized by neuronal loss, and increased astrocytosis. The patient organoids showed irregular neuroepithelial tissue organization, reduced ventricle-like structures, and diminished expression of cortical neuron markers SATB2 and TUJ1. In contrast, there was a notable elevation in the expression of the activated astrocyte marker GFAP. These findings suggest that *POLG* mutations contribute to aberrant growth and altered neuronal development in cortical organoids.

To gain insights into the cellular composition and gene expression dynamics of the cortical organoids, we performed scRNA-seq analysis. Our scRNA-seq analysis revealed eight distinct cell population clusters in the control organoids, including astrocytes, dopaminergic neurons, glutaminergic neurons, GABAergic neurons, neural progenitor cells, ependymal cells, radial glial cells, and DA GLU neurons. The DA GLU neurons were further classified into subpopulations based on gene expression profiles. Pathway enrichment analyses on the neuronal population confirmed the enrichment of gene sets associated with neural development processes, such as axon development, neural migration, generation of neurons, and synaptic transmission. Results from both immunostaining and scRNA-seq revealed changes in cell type number and composition in cortical organoids from patients compared to healthy controls. Neuronal loss is one of the most severe phenotypic manifestations in the spectrum of POLG diseases[39–41] and we found significant neuronal loss in patient cortical organoids, including cortical neurons and GABAergic DA GLU neurons. The loss of DA GLU neurons is particularly interesting since postmortem studies have identified loss of this neuronal subtype [42] as a feature of severe POLG related disease.

Our study not only revealed the loss of neurons but also highlighted the presence of astrocyte activation and reactive gliosis in patient-derived organoids. The observed astrocyte activation is particularly significant as it has been shown in our study [43] and previous research [44] to be associated with neuronal toxicity and may contribute to the neural loss observed in POLG-related disorders. Astrocytes play crucial roles in supporting neuronal function and maintaining brain homeostasis. However, under pathological conditions, such as *POLG* mutations, astrocytes can become activated and undergo reactive gliosis. This reactive response is characterized by changes in astrocyte morphology, gene expression, and secretion of various signaling molecules. The activation of astrocytes in POLG patient organoids suggests a potential role for these cells in contributing to the disease pathology. Studies have demonstrated that activated astrocytes can release pro-inflammatory factors, reactive oxygen species, and other neurotoxic substances, which can lead to neuronal dysfunction and degeneration. These toxic effects on neurons may contribute to the observed neuronal loss in POLG disease. Moreover, the presence of reactive gliosis in patient-derived organoids further supports the idea that these iPSC-derived brain models can recapitulate key pathological features of POLG disease. The activation of astrocytes and the subsequent gliosis response are hallmarks of various neurodegenerative disorders, including POLG-related disorders. This similarity in pathological features between patient-derived organoids and actual disease conditions strengthens the validity and relevance of these organoid models for studying POLG disease mechanisms. By recapitulating the pathological changes observed in patient brains, iPSC-derived brain organoids offer a valuable platform for investigating the underlying mechanisms of POLG disease. These models provide an opportunity to explore the interactions between different cell types, such as neurons and astrocytes, and the impact of these interactions on disease progression. Furthermore, they enable the testing of potential therapeutic strategies aimed at modulating astrocyte activation and mitigating the neurotoxic effects associated with POLG-related disorders [45, 46].

In our study, we made an intriguing observation that patient organoids with *POLG* mutations exhibited a loss of both neurons and markers associated with mature neuronal identity. Neurons are typically post-mitotic cells that form during embryonic development and maintain their specialized characteristics, functionality, and resilience throughout an individual’s lifespan. However, in certain circumstances, cell dedifferentiation can occur during developmental processes or in response to stress or injury [47]. Recent evidence also suggests that loss of cellular identity is associated with neuronal vulnerability and neurogenesis [48–51]. The loss of mature neuronal markers suggests that POLG disease neurons may dedifferentiate, thereby reactivating the developing neural cell cycle. Evidence from familial AD-iPSC-derived neurons and postmortem AD brains also supports this phenomenon [21, 52–54]. Additionally, our findings indicate that patient organoids exhibited a loss of postsynaptic markers, although the specific marker is not mentioned in the provided text. In vivo studies have revealed that postsynaptic cells play a critical role in initiating presynaptic differentiation and stabilizing immature synaptic contacts, enabling them to mature and respond to signals that promote synaptic maturation. Thus, our findings suggest that POLG disease either leads to blocked postsynaptic differentiation [55], or that postsynaptic markers are lost as part of the dedifferentiation process, while presynaptic markers remain.

At the transcriptomic level, scRNA-seq highlighted cell-type-specific variations in POLG organoids. In neuronal populations, we observed a notable downregulation of pathways tied to neuronal differentiation. This observation supports our results showing loss of mature neuronal markers that suggested a state of dedifferentiation. Loss of established neuronal identity and reversion to a more progenitor-like state has, moreover, been observed in other neurodegenerative diseases [56]. Intriguingly, we noticed upregulation of the NOTCH and JAK-STAT signaling pathways. The NOTCH signaling pathway plays a crucial role in maintaining neural progenitor cells and inhibiting neuronal differentiation, which aligns with our observation of impaired neuronal maturation [57]. The JAK-STAT pathway is primarily associated with cellular responses to cytokines and growth factors and has been implicated in neuroinflammatory responses and glial cell activation [57]. This upregulation could potentially reflect the observed astrocytosis in our model, indicating a response to neuronal damage or that astrocytes are playing a more active role in the disease process. These results exemplify the complex, multifaceted cellular changes resulting from *POLG* mutations and the potential interplay between genetic changes, cellular differentiation status, and intercellular signaling in disease progression.

We assayed the mitochondrial consequences of *POLG* mutations in ‘living organoid cells’ and observed loss of complex I and mtDNA copies, consistent with our findings in patient postmortem brain tissue [38] and 2D neuronal cell models [12, 13]. The simultaneous loss of neurons in patient organoids made it attractive to postulate a causal relationship between complex I loss and neural cell death, however, complex I loss was found to affect the whole brain in PD [58], not just the substantia nigra [59, 60], suggesting that loss of this complex may not itself be the primary cause of neuronal loss, but a compensatory response [61]. Irrespective of whether neuronal loss of complex I a pathological event or compensatory response is, this feature is clearly important and associated with the POLG disease process in both in patients and our cortical organoids. Following changes in level of both complex I and mtDNA may, therefore, be useful for monitoring treatment or other interventions.

Another key molecular consequence of *POLG* mutations is the presence of mtDNA depletion [34, 62]. We have shown that this is present in neurons from infants under 1 year of age and a stable finding in surviving neurons of patients from all age groups [34]. Here, using the patient cortical organoids, we also observed lowered mtDNA levels using an indirect method based on flow cytometric measurement of TFAM. While this method relies on an indirect assessment, TFAM binds mtDNA in molar quantities and we suggest that it can be used in live cells as a surrogate measure of mtDNA level [13, 26, 27]. The presence of mtDNA depletion appear to impair cell growth, as in the case of patient [25]. The presence of mtDNA depletion has important implications for cellular function and growth. In the case of the patient we studied, impaired cell growth was observed, which is consistent with previous reports linking mtDNA depletion to cellular dysfunction and compromised cell viability. The reduction in mtDNA levels may disrupt mitochondrial function, as mtDNA encodes essential genes involved in oxidative phosphorylation and ATP production. This impairment in energy production can negatively impact cell growth and survival. Efforts to increase mtDNA levels in cells with *POLG* mutations have been explored as a potential therapeutic strategy. One approach involves the use of mitochondrial-targeted nucleotides or nucleotide analogs to enhance mtDNA replication and restore mtDNA levels [63]. These compounds can bypass the impaired mtDNA replication machinery caused by *POLG* mutations and promote the synthesis of new mtDNA copies. However, it should be noted that the development of effective and safe therapies targeting mtDNA depletion in POLG-related disorders is still in the early stages and requires further investigation.

Our results indicate that the POLG organoid model is suitable as a platform for drug testing and screening. Parameters such as cellularity and cell type complements, complex I quantity, mitochondrial mass as measured by TOMM20 quantity, and mtDNA copy number measured indirectly by assessing TFAM, can serve as sensitive readouts to assess drug effects. As proof of concept, we show that exposure to metformin, a drug shown to boost mitochondrial biogenesis, largely mitigates the impact of *POLG* mutation, indicating therapeutic potential. Although metformin can inhibit mitochondrial complex I activity at low dosage, data from our previous study [28] and other reports suggests that higher doses (250 μM - 1000 μM) of metformin increase complex I activity [64]. We tested metformin and found that 250 μM for 2 months improved the mitochondrial deficit and partly mitigated the neuronal loss. These results nominate metformin as a potential candidate for clinical trials in POLG patients, subject to further preclinical validation.

Our findings must be interpreted in the light of the following limitation: we used healthy, age-matched control iPSCs and patient-specific iPSCs, which were differentiated into cortical organoids. We are fully aware that the current state-of-the-art in iPSC research is the use of genetically corrected isogenic controls. However, it has been reported that the high efficiency of genome cleavage and repair makes the introduction of heterozygous alleles by standard CRISPR/Cas9 technology difficult [13, 27, 65, 66]. Thus, we felt that the presence of compound heterozygous mutations, such as were present in the patient used in this study, made the generation of isogenic controls impractical. We feel, nevertheless, that our results are important and if we look only at the patient organoids, internally consistent. They show a clear abnormality and response to metformin that brings them in line with what is seen in controls.

Despite the advances represented by our POLG organoid model, there are limitations to be addressed. In our paradigm, the organoid culture method suppressed mesoderm-derived immune cell/microglia presence [67] in order to achieve a more consistent forebrain identity and cell composition. This allowed us to address initial astroglia and neuron dependent pathologies without immune or secondary inflammatory responses. However, in future, the supplementation of POLG organoid models with microglia/allogeneic immune cells could allow us to address any immune-related pathogenesis in POLG-related disease. Evidence suggests that neuroinflammation and mitochondrial dysfunction may trigger a vicious circle: dysfunctional mitochondria induce inflammation, and inflammation induces mitochondrial dysfunction [68]. It would be exciting, therefore, to address how immune cells affected by the *POLG* mutation contribute to cell vulnerability and disease pathogenesis in POLG organoids.

Our study highlights the unparalleled benefits of using the 3D cerebral organoid model. This model excels in differentiating and producing a variety of brain cell types commonly found in the human brain, such as neurons, astrocytes, and neural stem cells. But it’s not just about cellular differentiation: our in-depth histological and functional analyses reveal that these 3D cerebral organoids closely mimic the human brain in both structure and functionality, closely resembling genuine brain tissue. What sets our study apart is our emphasis on the differentiation of POLG cerebral organoids. These specialized 3D constructs not only display molecular markers indicative of POLG-related diseases but also shed light on the distinct physiological and functional characteristics of these conditions. This provides an unparalleled resource for studying POLG-linked disorders, enabling research to encompass interactions among a variety of cell types, rather than just focusing on individual cells. By harnessing the potential of these 3D cerebral organoids, we are well-positioned to explore the complex relationships between different brain cells, uncover the molecular underpinnings of diseases, and lead the charge in drug testing and validation. Most importantly, this model offers a research platform that truly reflects the human physiological scenario, which not only boosts the biological significance of our work but also improves the likelihood of success in drug discovery and validation.

## Data availability

The scRNA-seq data is stored in the external repository GEO GSE234659. The remaining data is available upon request.

## Acknowledgements

We would like to express our gratitude to the staff of the Center for Neuro-SysMed, Department of Neurology, Haukeland University Hospital, for their support and provision of the necessary personnel and infrastructure for this study. This work was supported by funding from the University of Bergen Meltzers Høyskolefonds (#103517133), Norwegian Research Council (#229652, #62613) and Rakel og Otto Kr.Bruuns legat (#809432, #103816102). The funders had no role in study design, data collection, data analysis, data interpretation, or writing of the report.

## Author contributions

KL and AC conceptualized the study. KL, AC and YH were involved in methodology. KL, AC, YH, TY, BCL conducted the investigation. KL and AC wrote the original draft. KL, AC, YH, TY, BCL, GJS, CT and LAB contributed to writing, review, and editing. KL, LAB and GJS acquired funding. KL and LAB provided resources. KL supervised the study. All authors have agreed to authorship.

---

**Supplementary Figure 1:**
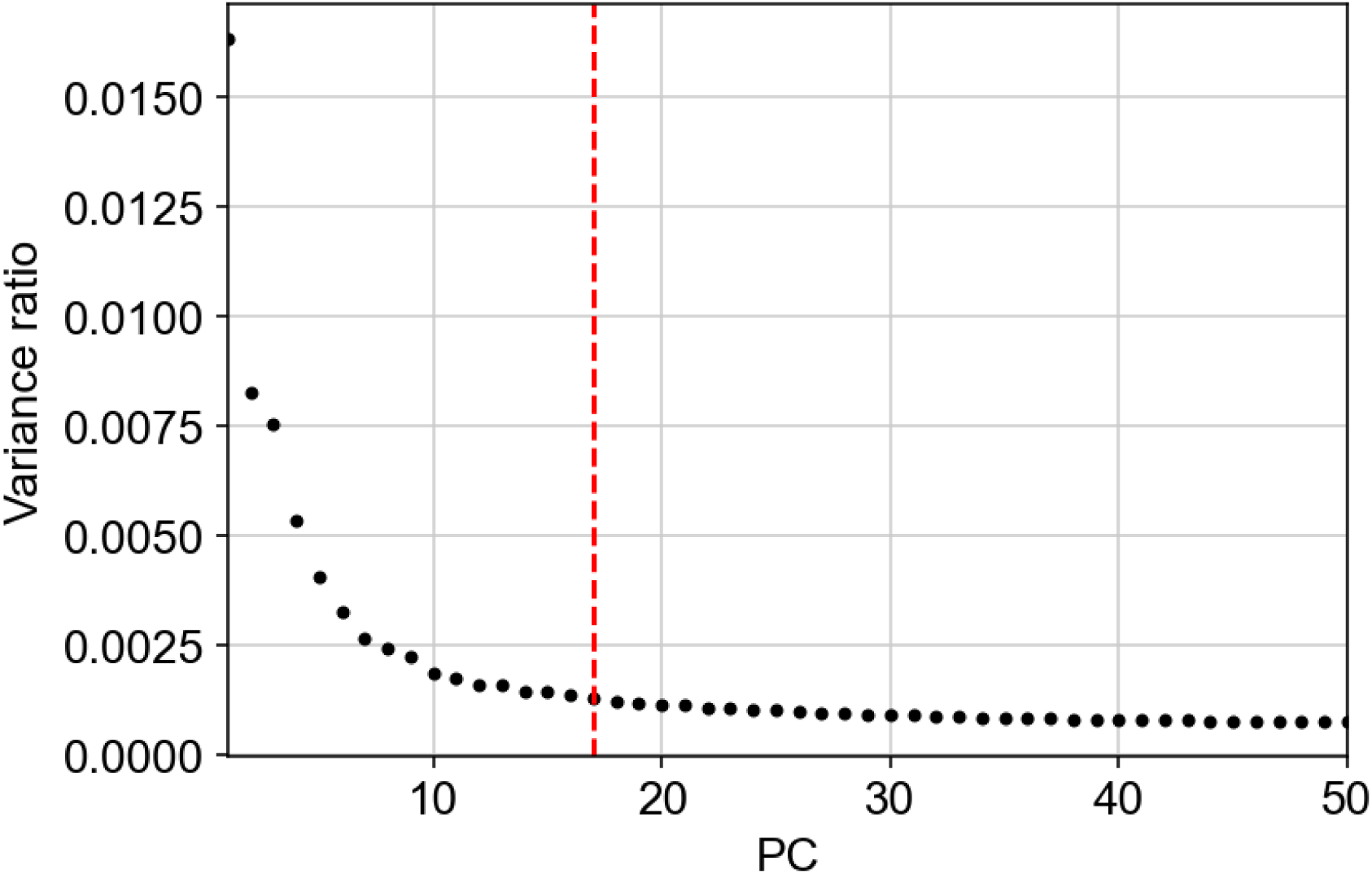
Detection of number of principal components before reaching the plateau. The plateau is reached at the 17th principal component (shown with a dashed red line).

**Supplementary Figure 2:**
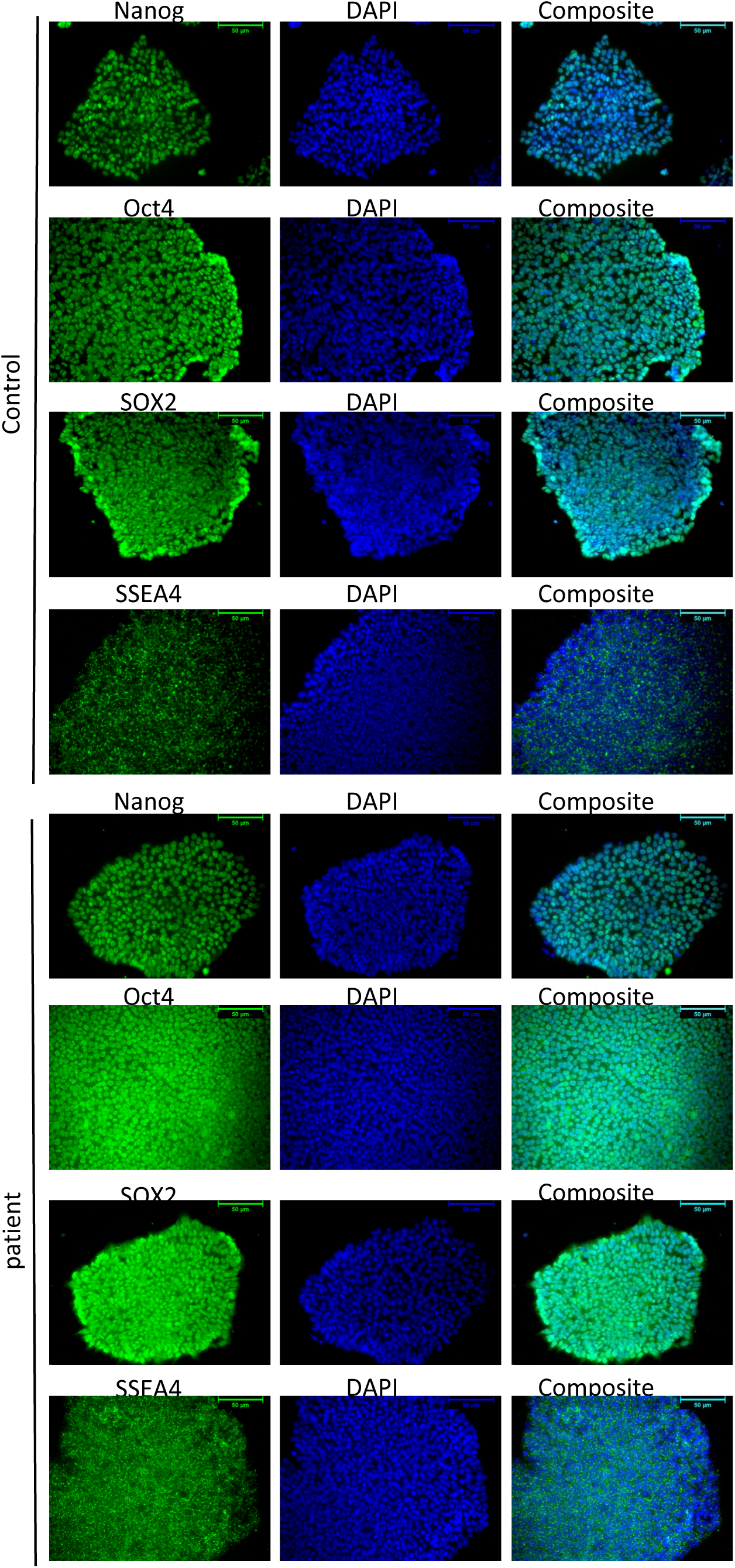
Panel of pluripotency marker expression in control and patient iPSCs.

**Supplementary Figure 3:**
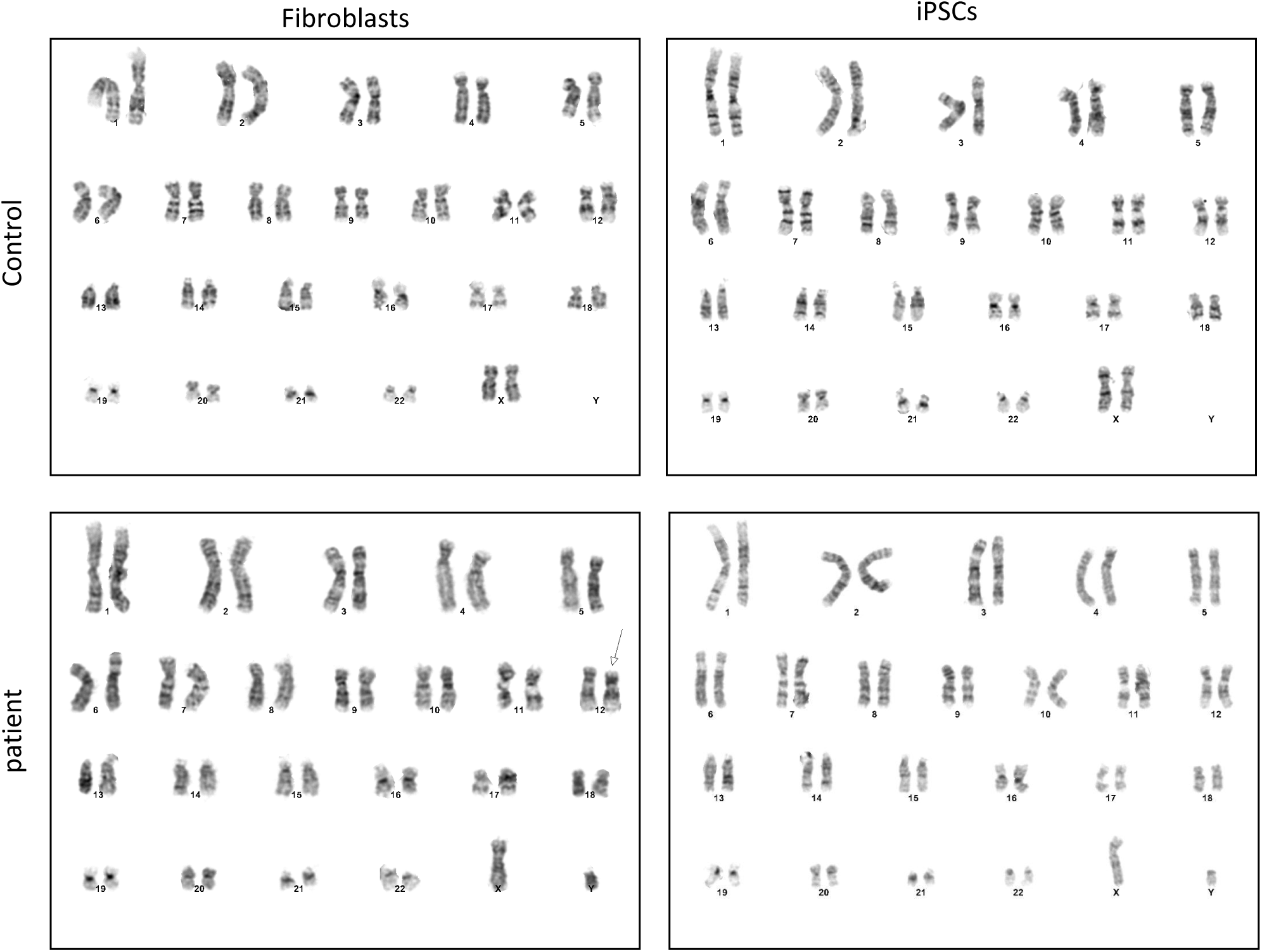
Representative karyotypes for control and patient POLG fibroblasts and reprogrammed iPSC lines.

**Supplementary Figure 4.**
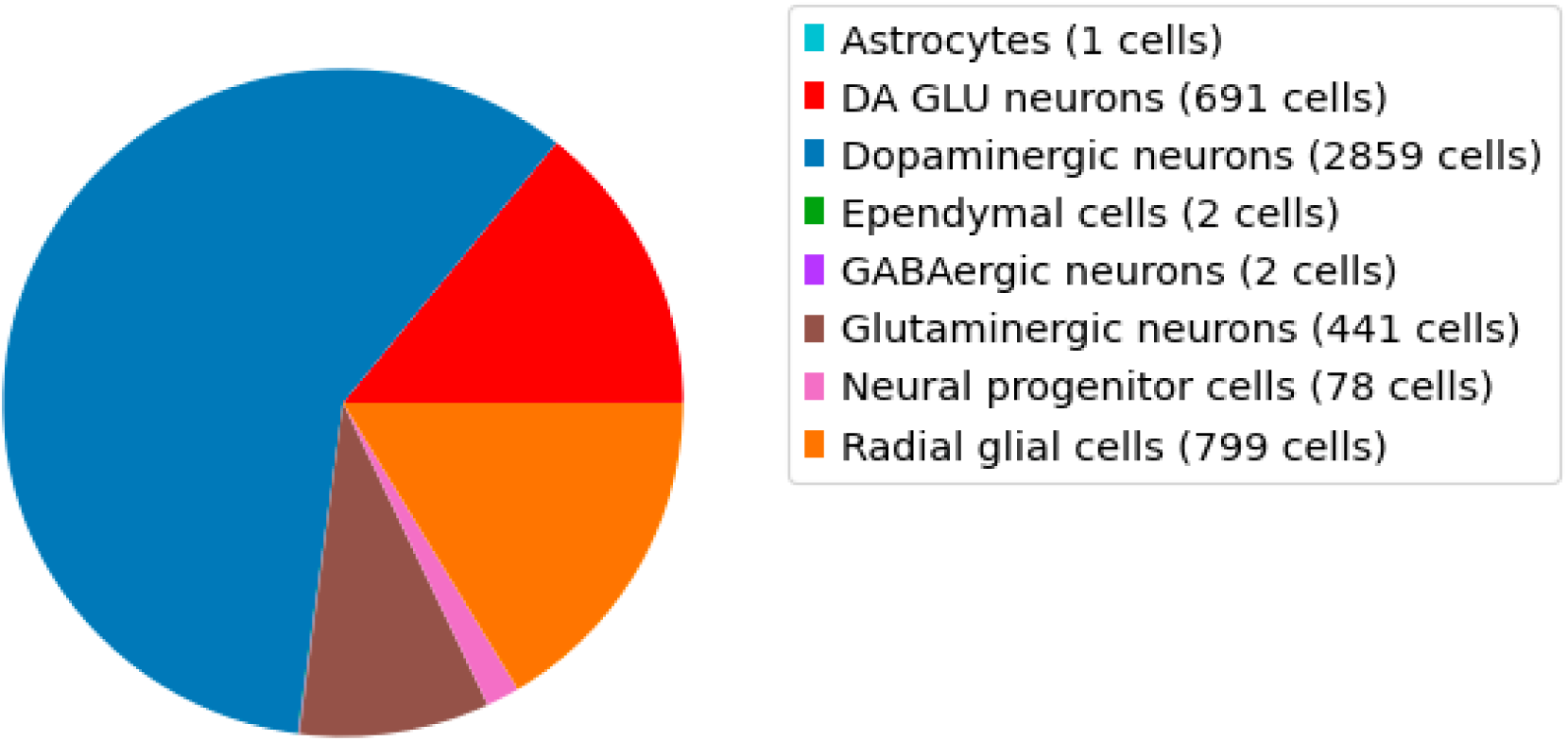
Number of each cell cluster in scRNA-seq of control cortical organoids.

**Supplementary Figure 5:**
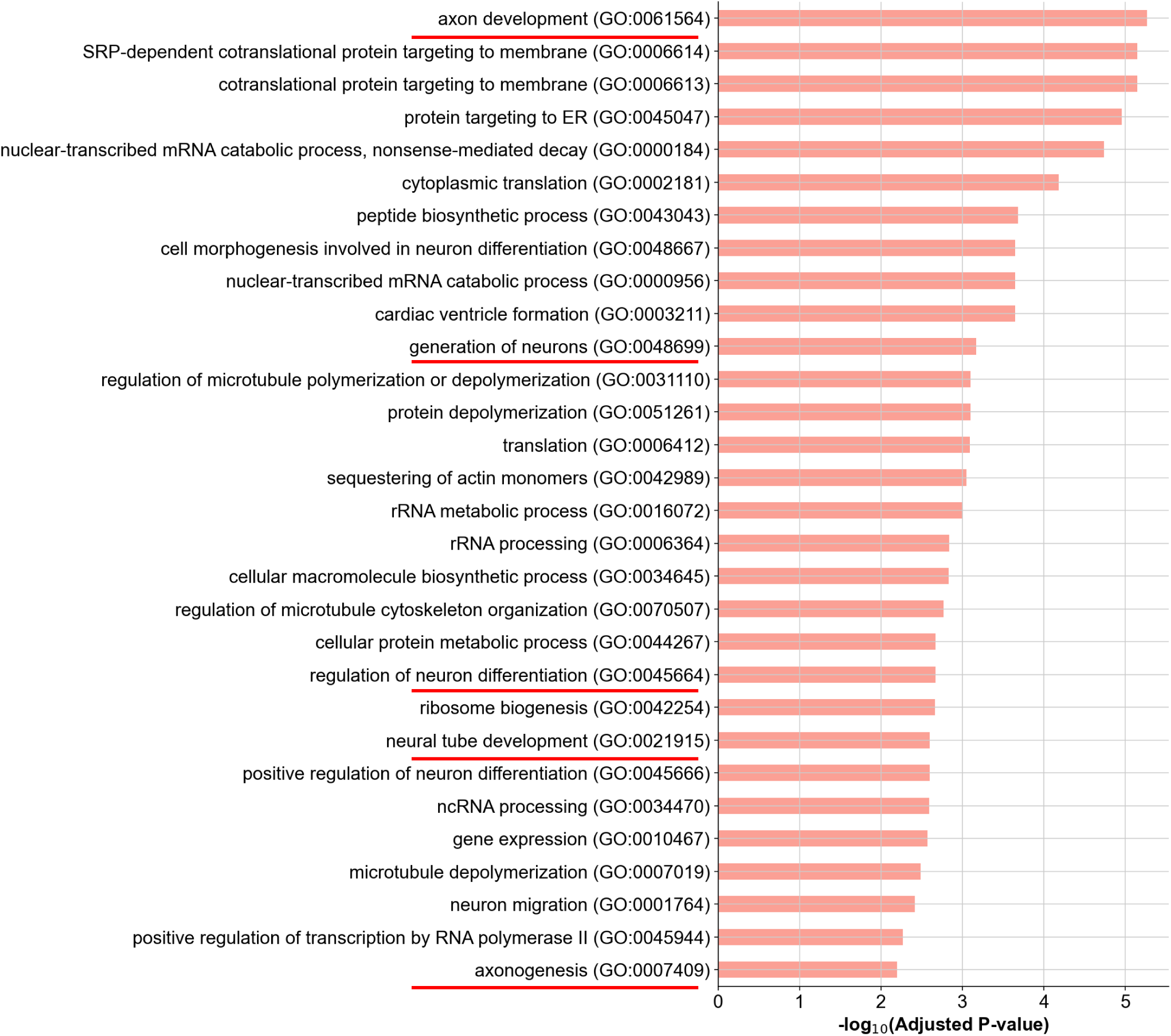
GO terms in neuron population enriched in the analysis of control cortical organoids.

**Supplementary Figure 6:**
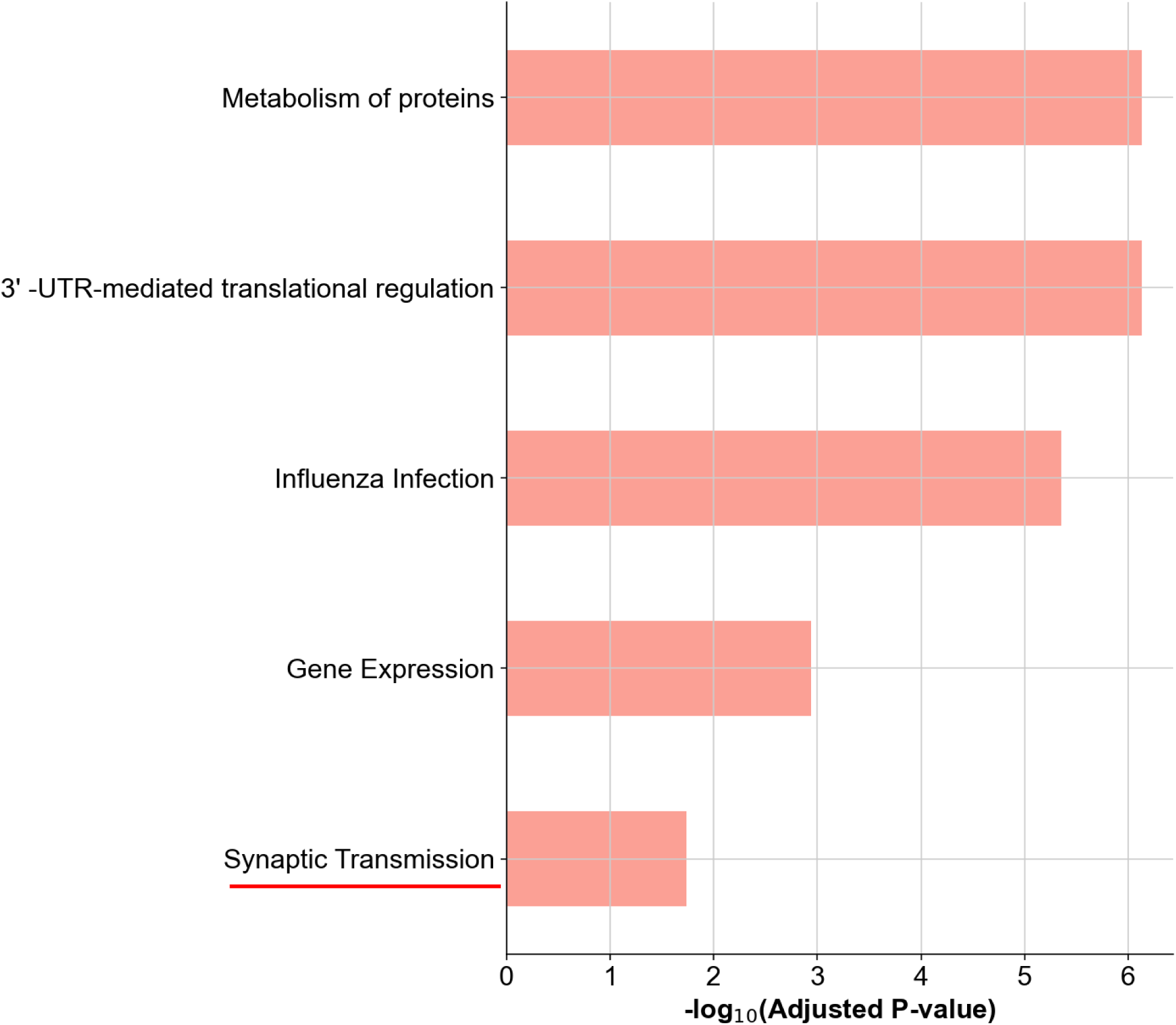
Reactome terms enriched in the analysis of neuron population in control cortical organoids.

**Supplementary Figure 7.**
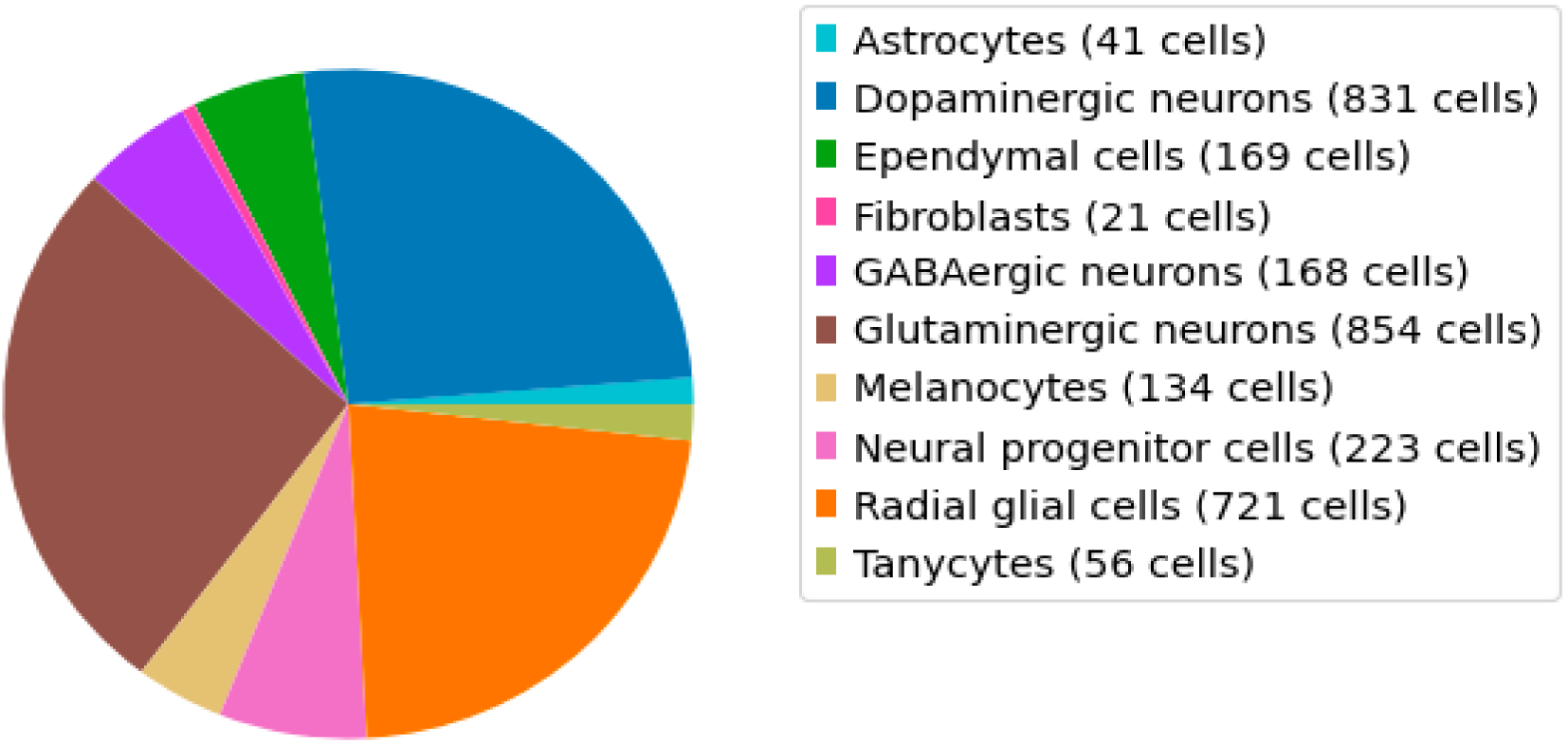
Number of each cell cluster in scRNA-seq of patient cortical organoids.

**Supplementary Figure 8:**
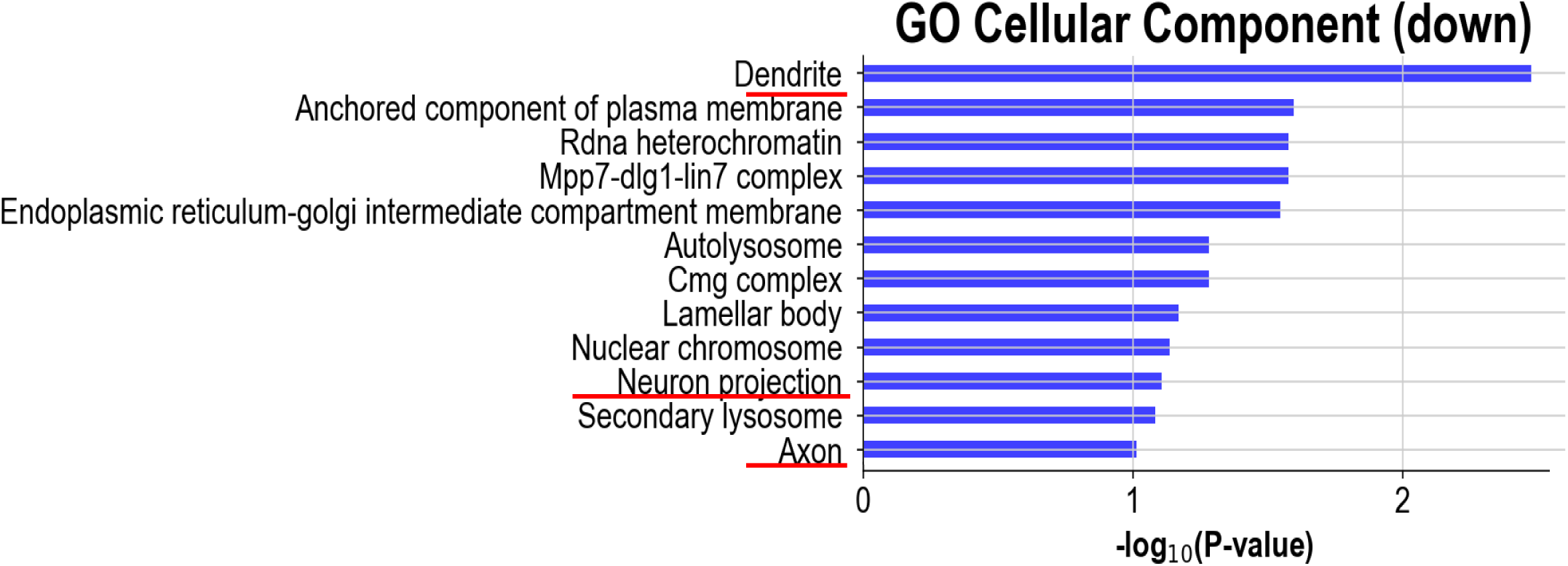
GO cellular components enriched in the analysis of the downregulated DEGs of the neuron population in patient cortical organoids versus control.

**Supplementary Figure 9:**
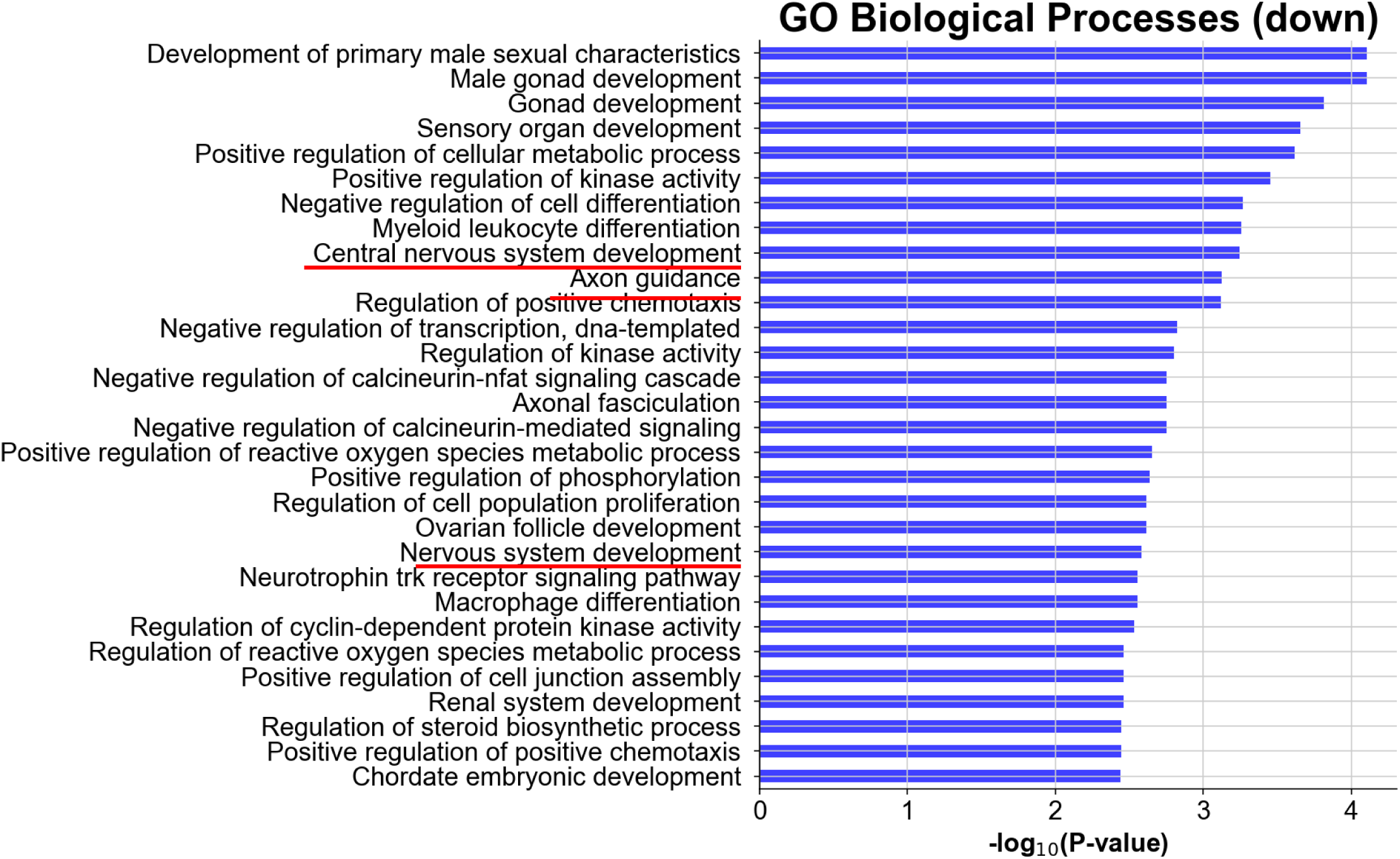
GO biological processes enriched in the analysis of the downregulated DEGs of the glial population in patient cortical organoids versus control.

**Supplementary Figure 10:**
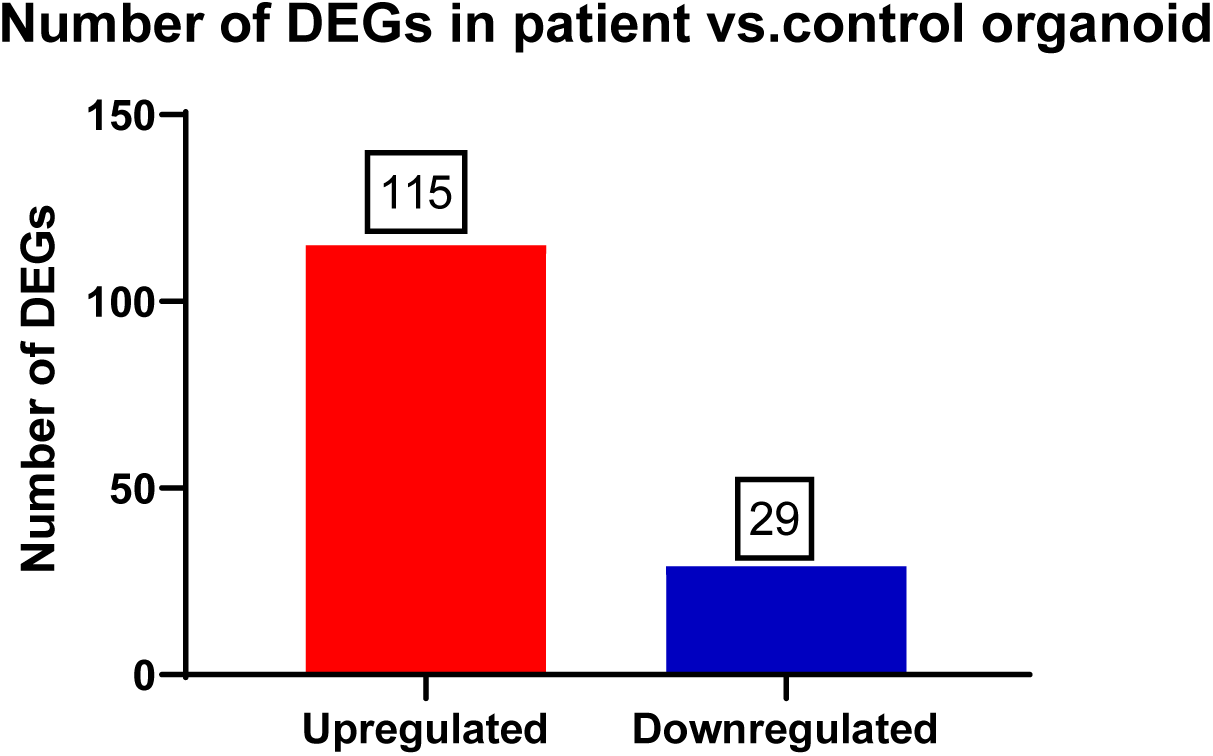
Numbers of up- and downregulated DEGs pooled for the dopaminergic neuron population in patient cortical organoids versus control organoids.

**Supplementary Figure 11:**
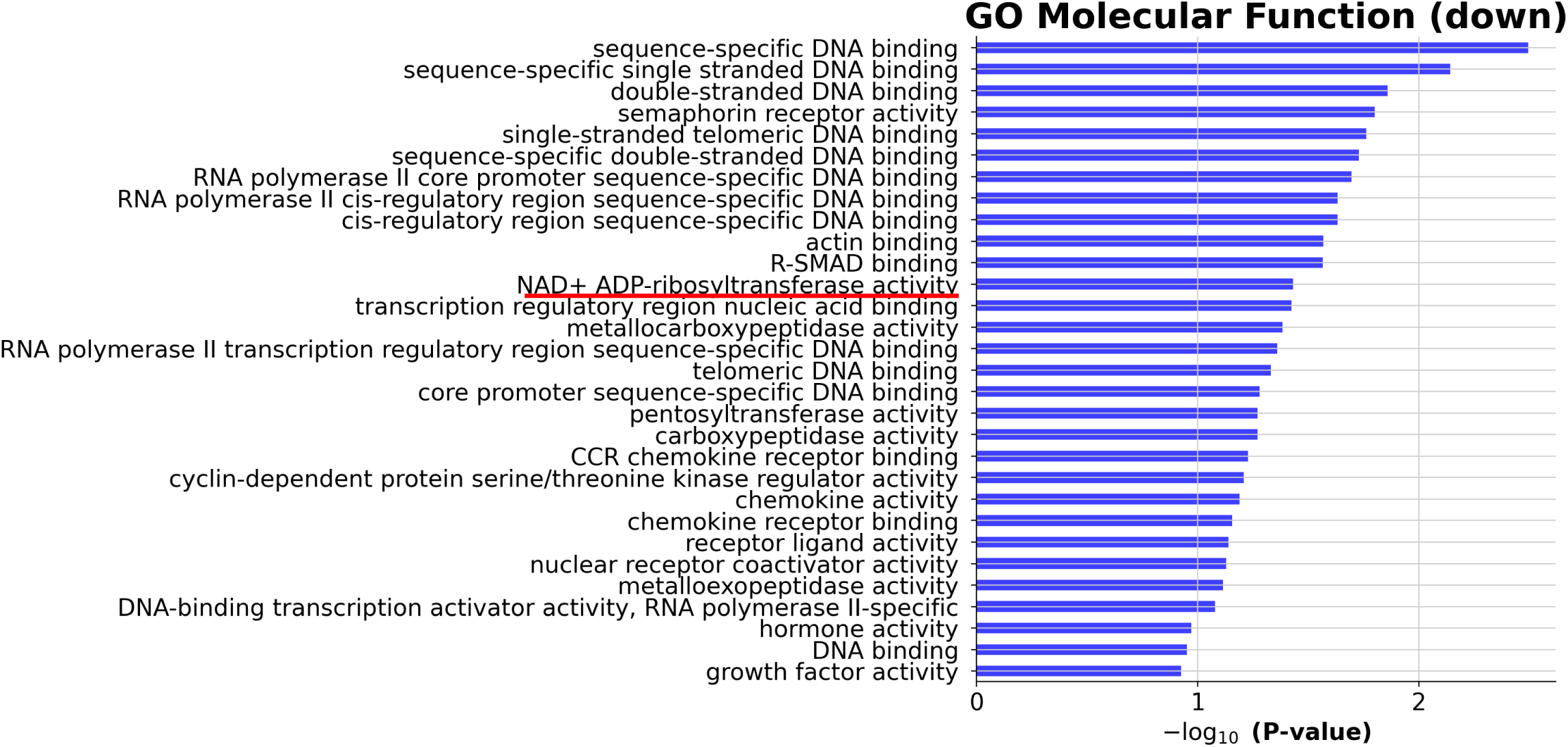
GO molecular function processes enriched in the analysis of the downregulated DEGs of the dopaminergic neuron population in patient cortical organoids versus control.

**Supplementary Figure 12:**
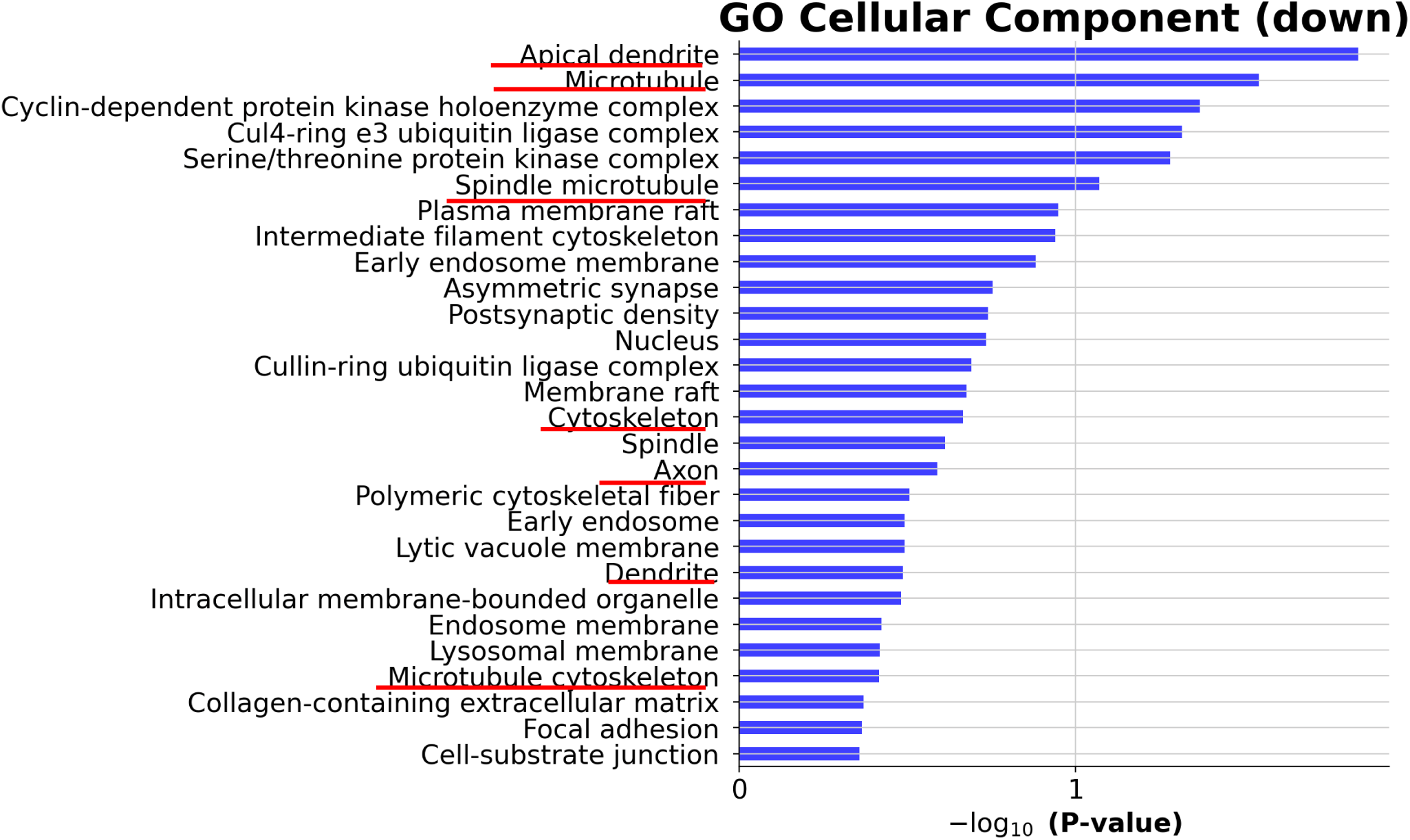
GO cellular components enriched in the analysis of downregulated DEGs of the dopaminergic neuron population in patient cortical organoids versus control.

**Supplementary Figure 13:**
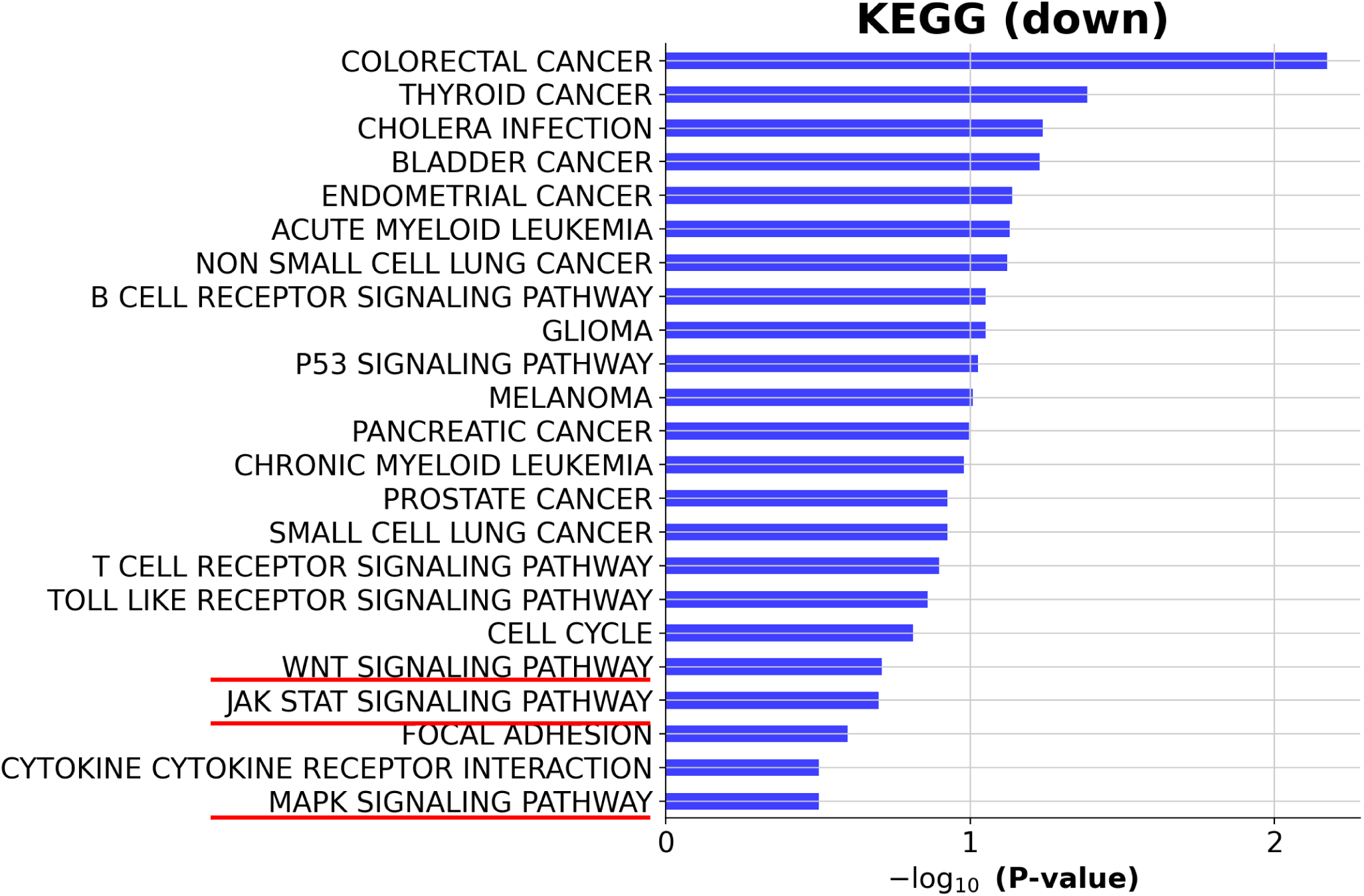
KEGG pathways enriched in the analysis of downregulated DEGs of the dopaminergic neuron population in patient cortical organoids versus control.

**Supplemental Figure 14.**
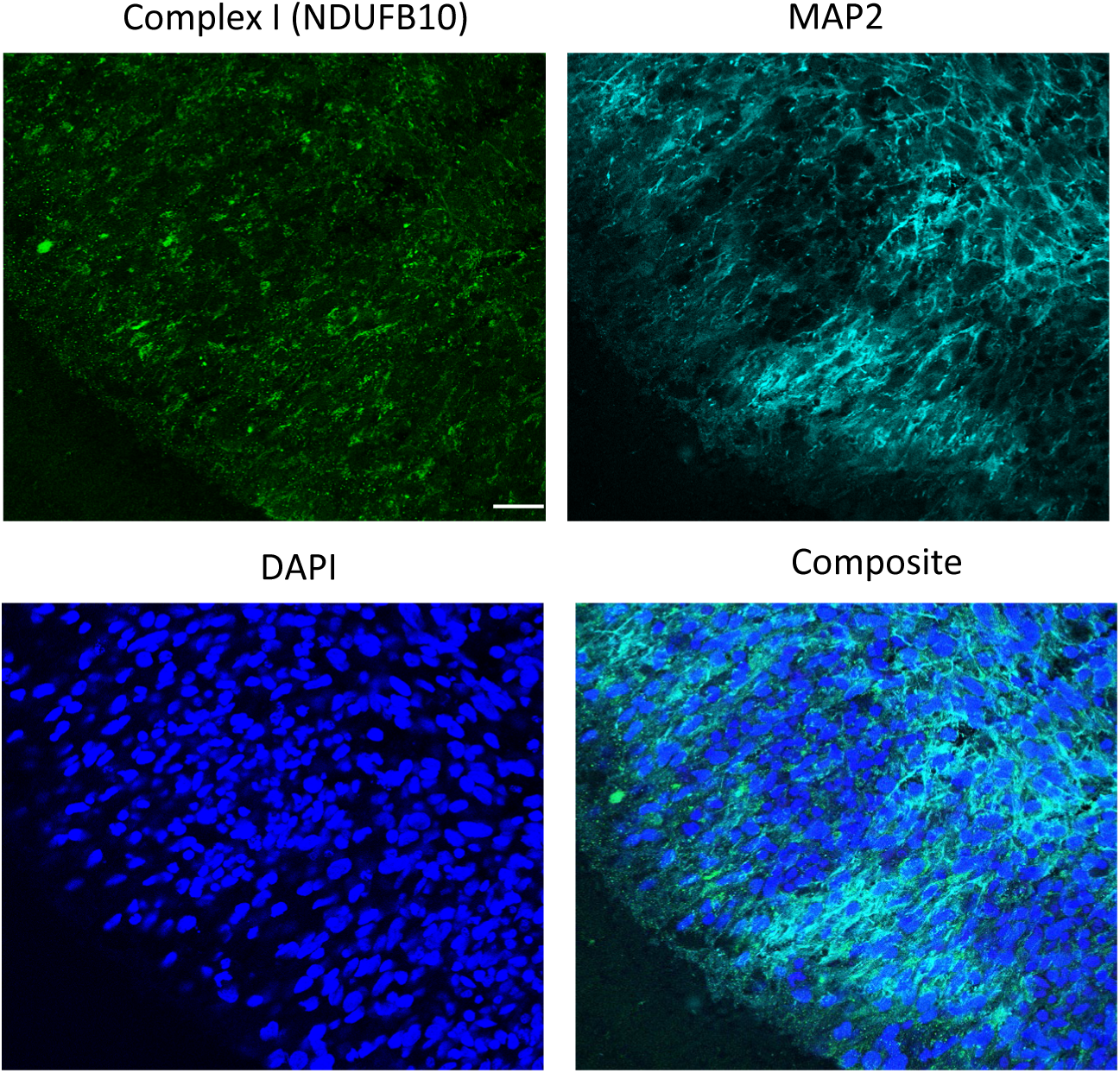
Immunofluorescent imaging of cryo-sectioned organoids at 90 days, showing the staining of NDUFB10 (green), GAD65 (red), and MAP2 (purple) in patient organoids and patient organoids treated with metformin. Nuclei are stained with DAPI (blue). Scale bar is 50 µm.

**Supplementary Figure 15.**
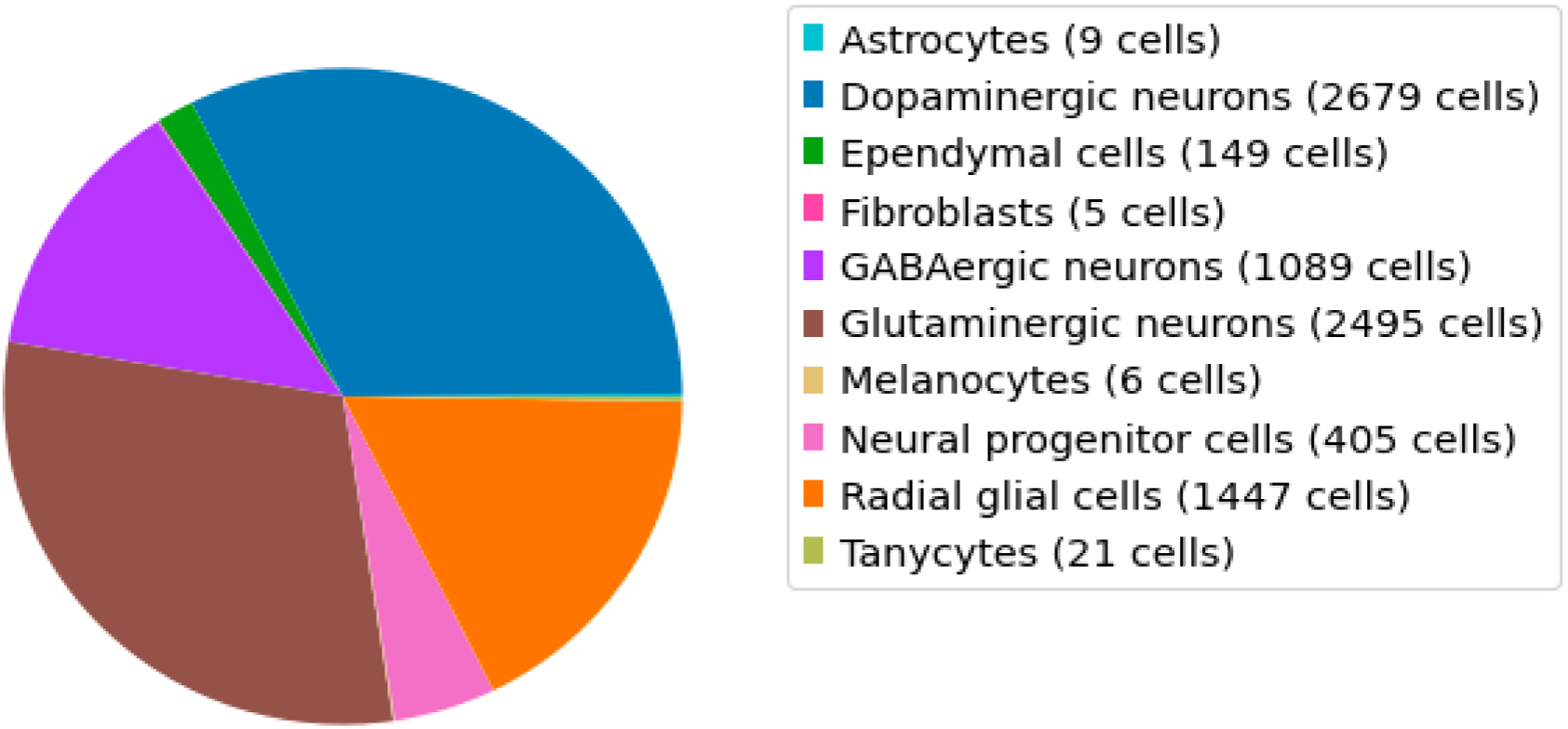
Number of each cell cluster in scRNA-seq of patient cortical organoids treated with metformin.

**Supplementary Figure 16:**
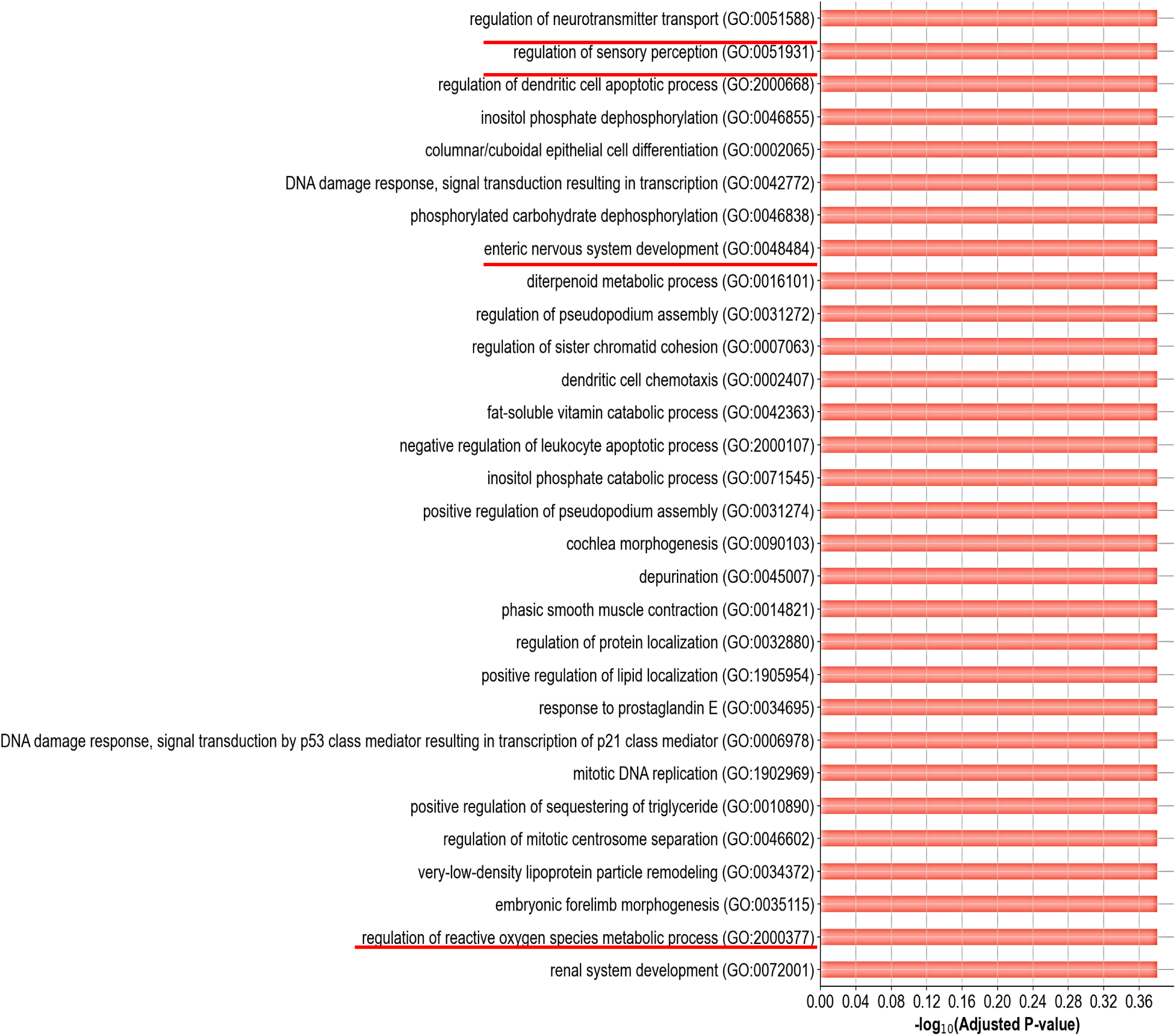
GO terms enriched in the analysis of the upregulated DEGs of the neuron population in patient cortical organoids with metformin treatment versus untreated organoids.

**Supplementary Figure 17:**
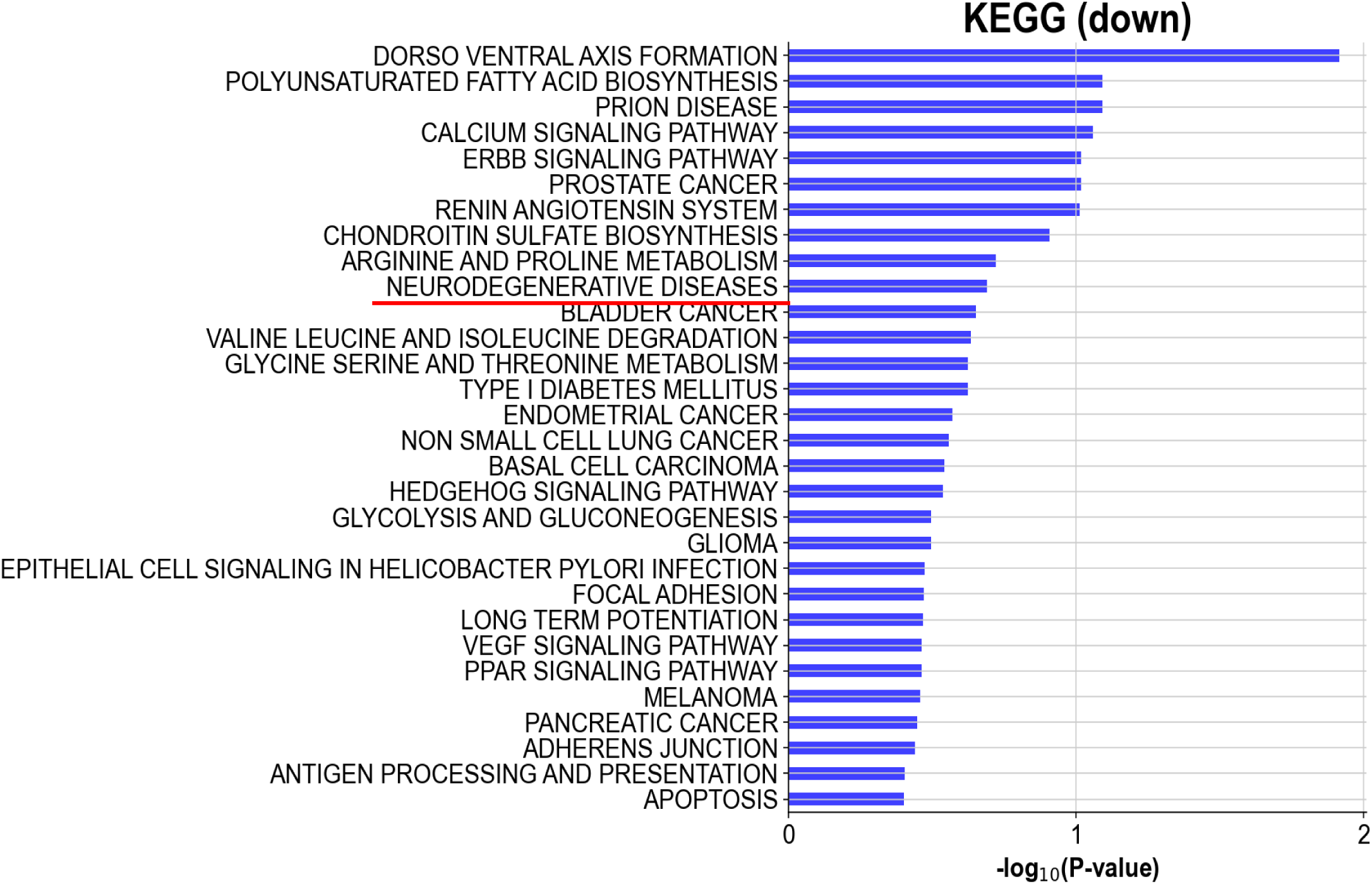
GO terms enriched in the analysis of the downregulated DEGs of the glial population in patient cortical organoids with metformin treatment versus untreated organoids.

**Supplementary Figure 18:**
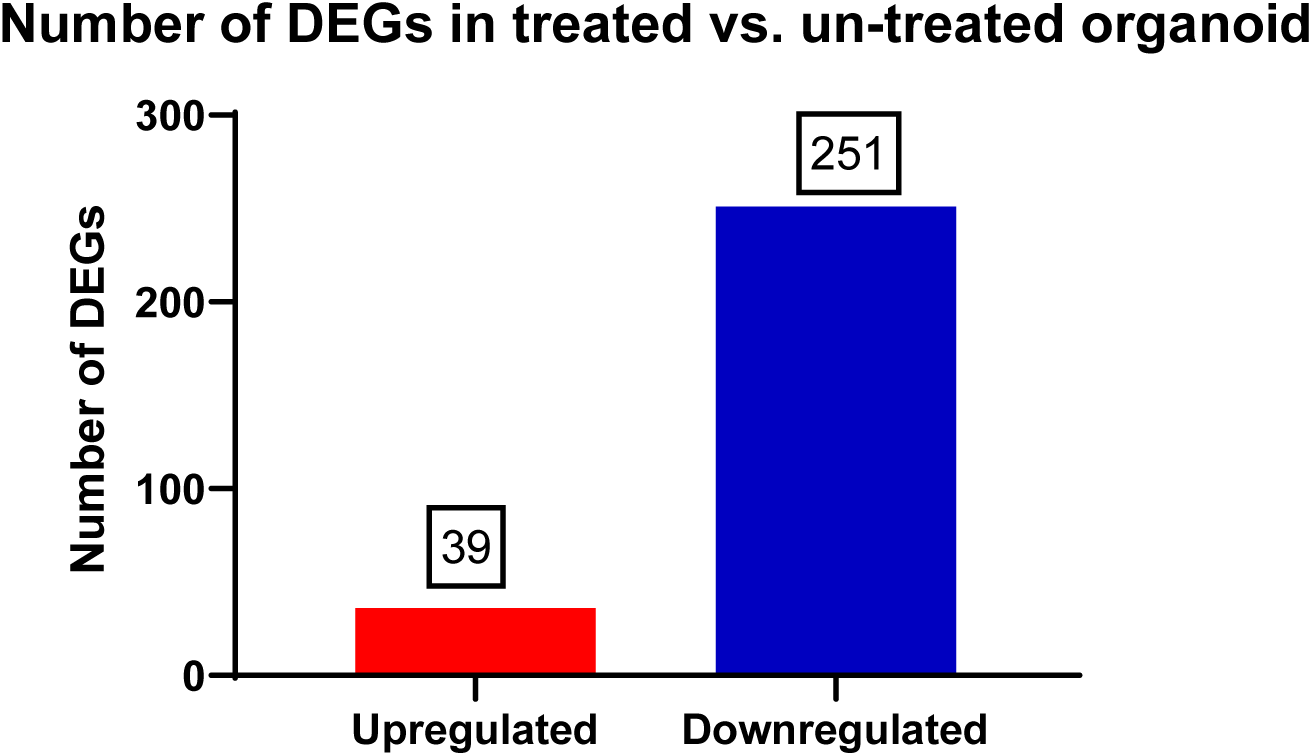
Numbers of up- and downregulated DEGs pooled for the dopaminergic neuron population in metformin treated patient cortical organoids versus untreated organoids.

**Supplementary Figure 19:**
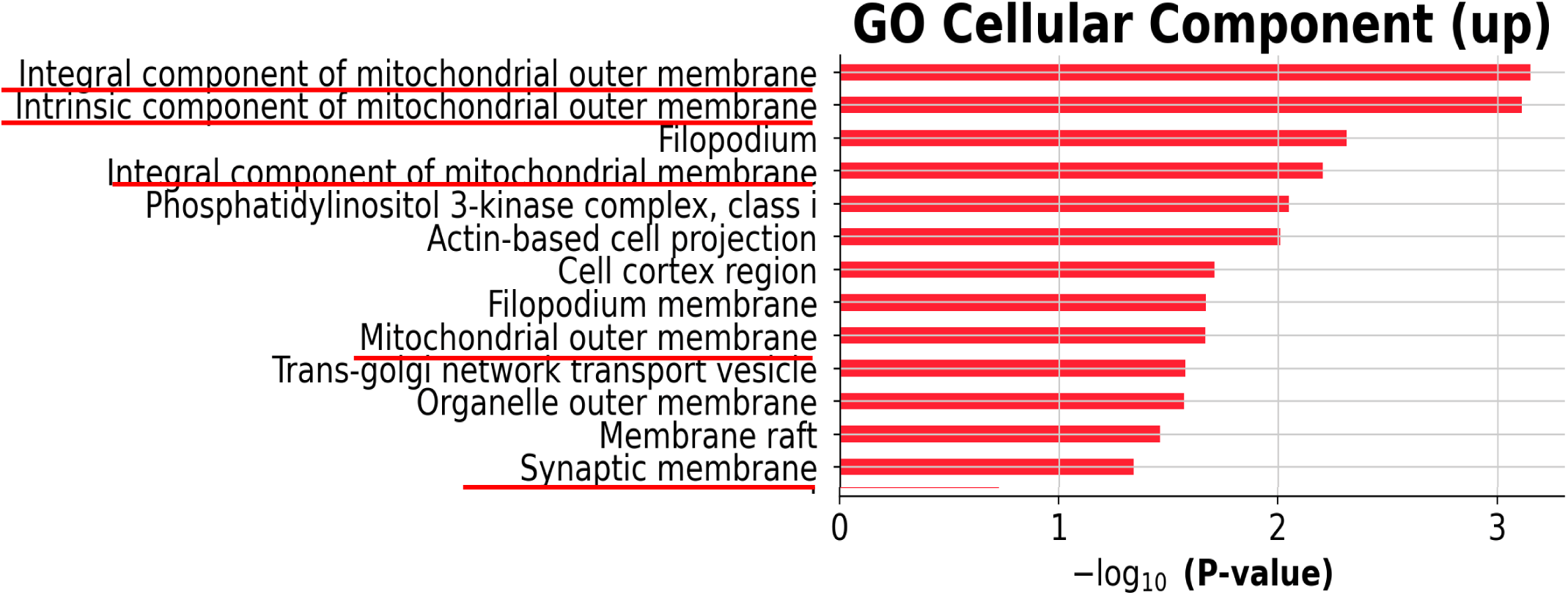
GO cellular components enriched in the analysis of the upregulated DEGs of the dopaminergic neuron population in patient cortical organoids with metformin treatment versus untreated organoids.

**Supplementary Figure 20:**
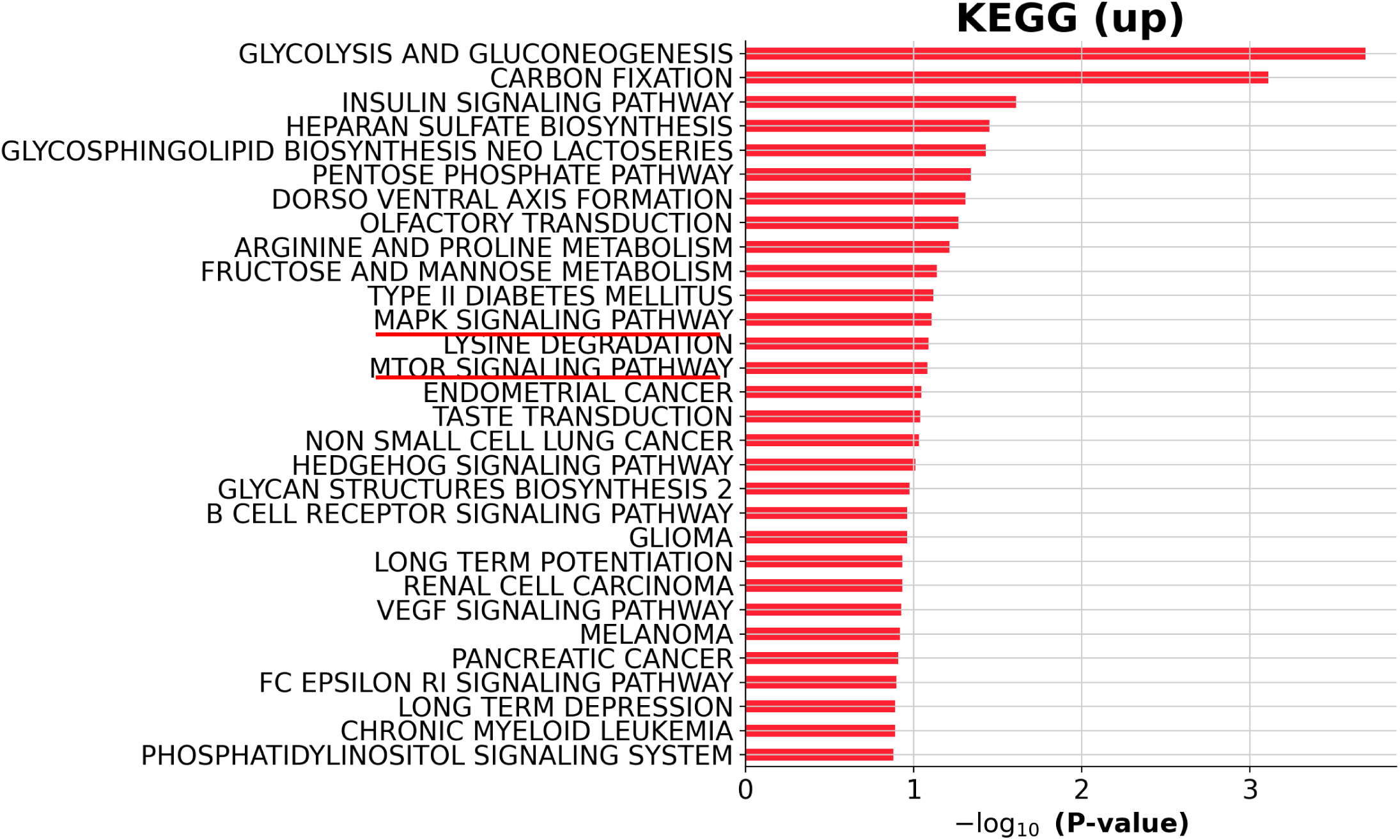
KEGG pathways enriched in the analysis of the upregulated DEGs of the dopaminergic neuron population in patient cortical organoids with metformin treatment versus untreated organoids.

**Supplementary Table 1:** List of the information for the neural induction medium.

| Component | Concentration | Supplier | Catalog number |
| --- | --- | --- | --- |
| DMEM/F-12 |  | Life Technologies | 11330057 |
| Knockout serum replacement |  | Life Technologies | 10828028 |
| MEM-NEAA | 1% (v/v) | Life Technologies | 35050 |
| β-Mercaptoethanol | 100 μM | Sigma-Aldrich | M7522 |
| LDN-193189 | 100 nM | Sigma-Aldrich | SML0559 |
| SB431542 | 10 μM | Abcam | ab120163 |
| XAV939 | 2 μM | Sigma-Aldrich | X3004 |

**Supplementary Table 2:** List of the information for the neural differentiation medium minus vitamin A.

| Component | Concentration | Supplier | Catalog number |
| --- | --- | --- | --- |
| DMEM/F-12 |  | Life Technologies | 11330057 |
| Neurobasal medium |  | Life Technologies | 2110349 |
| Insulin | 0.025% (v/v) | Sigma-Aldrich | I9278 |
| MEM-NEAA | 0.5% (v/v) | Life Technologies | 11140050 |
| Glutamax supplement | 1% (v/v) | Life Technologies | 35050 |
| Penicillin/Streptomycin | 1% (v/v) | Life Technologies | 15140-122 |
| N2 supplement | 0.5% (v/v) | Life Technologies | 17502-048 |
| B27 supplement without vitamin A | 1% (v/v) | Life Technologies | 12587010 |
| β-Mercaptoethanol | 50 μM | Sigma-Aldrich | M7522 |

**Supplementary Table 3:** List of the information for the neural differentiation medium minus vitamin A.

| Component | Concentration | Supplier | Catalog number |
| --- | --- | --- | --- |
| DMEM/F-12 |  | Life Technologies | 11330057 |
| Neurobasal medium |  | Life Technologies | 2110349 |
| Insulin | 0.025% (v/v) | Sigma-Aldrich | I9278 |
| MEM-NEAA | 0.5% (v/v) | Life Technologies | 11140050 |
| Glutamax supplement | 1% (v/v) | Life Technologies | 35050 |
| Penicillin/Streptomycin | 1% (v/v) | Life Technologies | 15140-122 |
| N2 supplement | 0.5% (v/v) | Life Technologies | 17502-048 |
| B27 supplement | 1% (v/v) | Life Technologies | 17504-04 |
| β-Mercaptoethanol | 50 μM | Sigma-Aldrich | M7522 |
| BDNF | 20 ng/ml | Pepro Tech | 450-02 |
| Ascorbic acid | 200 μM | Sigma-Aldrich | A92902 |

**Supplementary Table 4:** List of the information on the software used for scRNA-seq analysis.

| Analysis | Software | Version |
| --- | --- | --- |
| Pre-processing tool | CeleScope | 1.7.1 |
| Quality Control of reads | fastqc | 0.11.8 |
| Trimming | cutadapt | 1.17 |
| Alignment | STAR | 2.6.1a |
| Quantification | featureCounts | 2.0.1 |
| Downstream analysis | R | 4.1.3 |
|  | python | 3.9.12 |
|  | Seurat | 3.1.2 |
|  | scanpy | 1.9.1 |
|  | anndata | 0.8.0 |
|  | leidenalg | 0.8.10 |
|  | scvelo | 0.2.2 |
|  | Monocle2 | 2.4.0 |
| Enrichment analysis | gseapy | 0.10.8 |
| Gene annotation | scMRMA | 1.0 |

**Supplementary Table 5:**
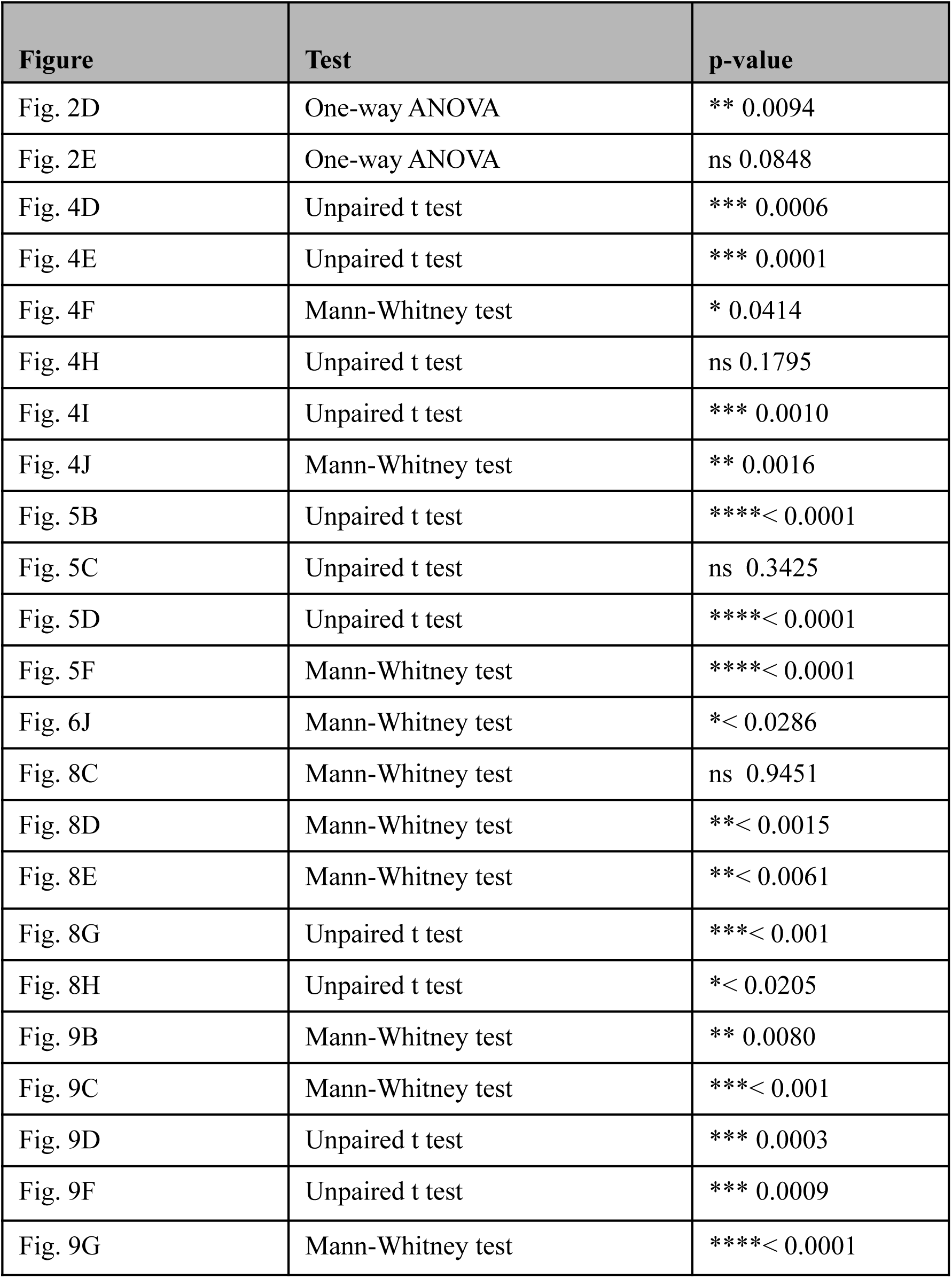
The list of statistical tests and p-values in the figures.

| **Figure** | **Test** | **p-value** |
| --- | --- | --- |
| Fig. 2D | One-way ANOVA | ** 0.0094 |
| Fig. 2E | One-way ANOVA | ns 0.0848 |
| Fig. 4D | Unpaired t test | *** 0.0006 |
| Fig. 4E | Unpaired t test | *** 0.0001 |
| Fig. 4F | Mann-Whitney test | * 0.0414 |
| Fig. 4H | Unpaired t test | ns 0.1795 |
| Fig. 4I | Unpaired t test | *** 0.0010 |
| Fig. 4J | Mann-Whitney test | ** 0.0016 |
| Fig. 5B | Unpaired t test | ****< 0.0001 |
| Fig. 5C | Unpaired t test | ns 0.3425 |
| Fig. 5D | Unpaired t test | ****< 0.0001 |
| Fig. 5F | Mann-Whitney test | ****< 0.0001 |
| Fig. 6J | Mann-Whitney test | *< 0.0286 |
| Fig. 8C | Mann-Whitney test | ns 0.9451 |
| Fig. 8D | Mann-Whitney test | **< 0.0015 |
| Fig. 8E | Mann-Whitney test | **< 0.0061 |
| Fig. 8G | Unpaired t test | ***< 0.001 |
| Fig. 8H | Unpaired t test | *< 0.0205 |
| Fig. 9B | Mann-Whitney test | ** 0.0080 |
| Fig. 9C | Mann-Whitney test | ***< 0.001 |
| Fig. 9D | Unpaired t test | *** 0.0003 |
| Fig. 9F | Unpaired t test | *** 0.0009 |
| Fig. 9G | Mann-Whitney test | ****< 0.0001 |

**Supplementary Table 6:** List of the information on the samples analyzed by scRNA-seq.

| Sample | Estimated Number of Cells | Mean Reads per Cell | Median UMI per Cell | Total Gene | Median Gene per Cell |
| --- | --- | --- | --- | --- | --- |
| Control organoid | 5,369 | 49,258 | 1,882 | 24,576 | 1,010 |
| patient organoid | 3,331 | 69,973 | 3,542 | 25,389 | 1,648 |
| patient organoid + metformin | 8,433 | 27,269 | 1,919 | 26,403 | 1,071 |

**Supplementary Table 7:** The list of top 10 genes for Figure 6A-D in cortical organoids of cell type enrichment analysis in scRNA-seq analysis.

| DA<br>GLU<br>neurons | Dopami<br>nergic<br>neurons | Ependym<br>al cells | GABAer<br>gic<br>neurons | Glutaminer<br>gic neurons | Neural<br>progenitor<br>cells | Radial glia<br>cells |
| --- | --- | --- | --- | --- | --- | --- |
| *SCG2* | *NEURO*<br>*D6* | *CA4* | *RAB8A* | *MT-CYB* | *RPS27* | *PTN* |
| *NSG2* | *TMSB10* | *WLS* | *PIGK* | *MT-ND5* | *FTL* | *C1orf61* |
| *MMP3* | *TUBA1*<br>*A* | *MGST1* | *ZDHHC5* | *NEFM* | *RPL21* | *VIM* |
| *ATP1B1* | *SOX4* | *PRTG* | *FZD3* | *MT-ATP6* | *FTH1* | *CLU* |
| *RTN4* | *STMN2* | *PLS3* | *CCM2* | *MT-ND4* | *RPL15* | *HES1* |
| *NEGR1* | *PTMA* | *RSPO2* | *MRPL47* | *NEFL* | *RPL13A* | *HSPB1* |
| *HSP90A*<br>*A1* | *NREP* | *FAM122B* | *PNRC1* | *MTND3* | *RPS5* | *MDK* |
| *CELF4* | *SOX11* | *WNT2B* | *RHBDD2* | *MT-ND2* | *CDKN1A* | *DBI* |
| *LMO3* | *MLLT11* | *CDO1* | *CDC42S*<br>*E1* | *MT-CO3* | *VIM* | *LINC01158* |

**Supplementary Table 8:** The list of GO terms and p-values in Supplementary Figure 5.

| **GO terms** | **p-value** |
| --- | --- |
| Axon development | 0.0000 |
| SRP-dependent cotranslational protein targeting to membrane | 0.0000 |
| Cotranslational protein targeting to membrane | 0.0000 |
| Protein targeting to ER | 0.0000 |
| Nuclear-transcribed mRNA catabolic process, nonsense-mediated decay | 0.0000 |
| Cytoplasmic translation | 0.0001 |
| Peptide biosynthesis process | 0.0002 |
| Cell morphogenesis involved in neuron differentiation | 0.0002 |
| Nuclear-transcribed mRNA catabolic process | 0.0002 |
| Cardiac ventricle formation | 0.0002 |
| Generation of neurons | 0.0007 |
| Regulation of microtubule polymerization or depolymerization | 0.0008 |
| Protein depolymerization | 0.0008 |
| Translation | 0.0008 |
| Sequestering of actin monomers | 0.0009 |
| rRNA metabolic process | 0.0010 |
| rRNA processing | 0.0015 |
| Cellular macromolecule biosynthetic process | 0.0015 |
| Regulation of microtubule cytoskeleton organization | 0.0017 |
| Cellular protein metabolic process | 0.0021 |
| Regulation of neuron differentiation | 0.0021 |
| Ribosome biogenesis | 0.0022 |
| Neural tube development | 0.0026 |
| Positive regulation of neuron differentiation | 0.0026 |
| ncRNA processing | 0.0026 |
| Gene expression | 0.0026 |
| Microtubule depolymerization | 0.0032 |
| Neuron migration | 0.0039 |

**Supplementary Table 9:**
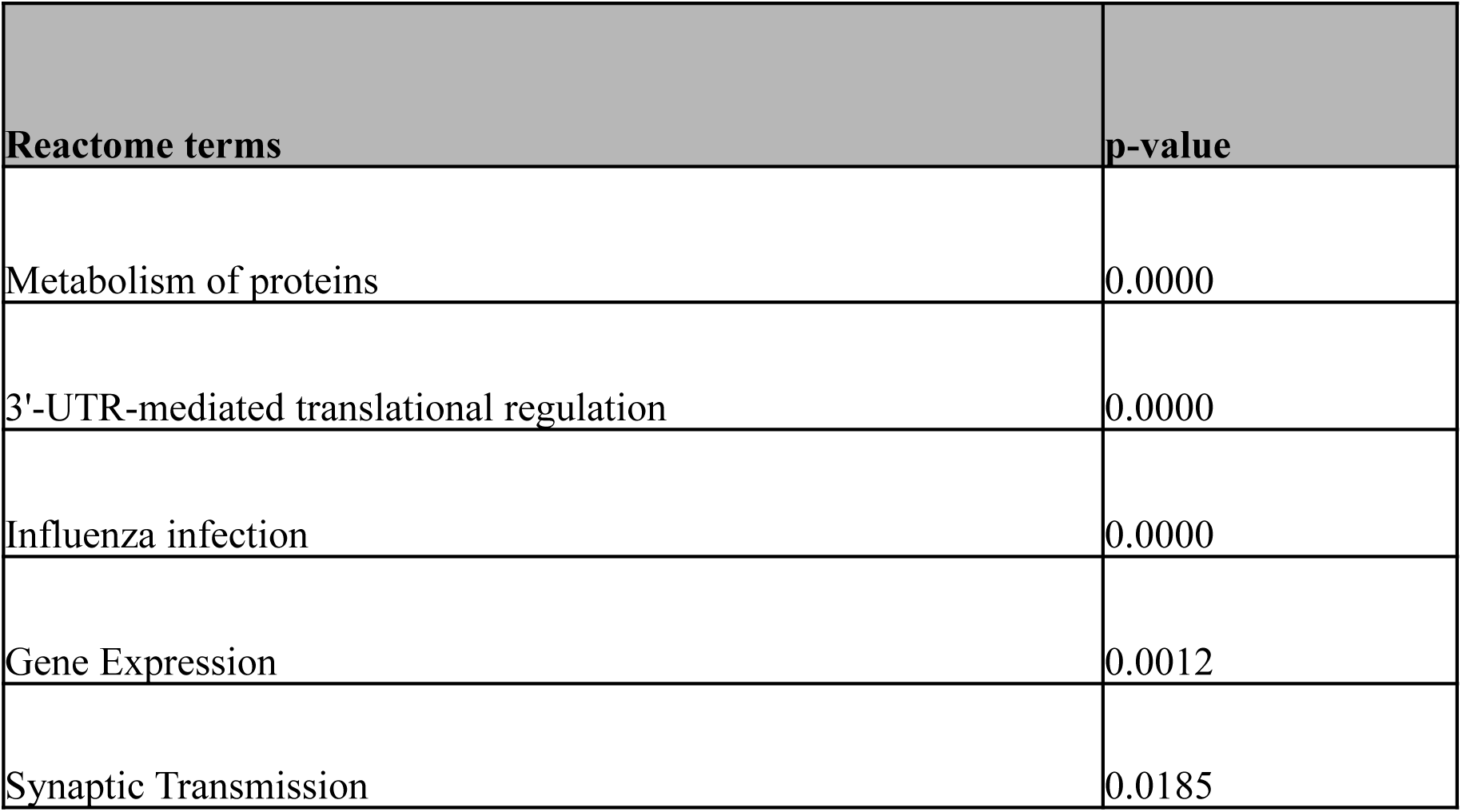
The list of Reactome terms and p-values in Supplementary Figure 6.

| Reactome terms | p-value |
| --- | --- |
| Metabolism of proteins | 0.0000 |
| 3'-UTR-mediated translational regulation | 0.0000 |
| Influenza infection | 0.0000 |
| Gene Expression | 0.0012 |
| Synaptic Transmission | 0.0185 |

**Supplementary Table 10:** The list of top 10 genes for Figure 6E-H in patient cortical organoids of cell type enrichment analysis in scRNA-seq analysis.

| **GABA**<br>**ergic**<br>**neuron**<br>**s** | **Glutam**<br>**inergic**<br>**neuron**<br>**s** | **Dopami**<br>**nergic**<br>**neurons** | **Neural**<br>**progen**<br>**itor**<br>**cells** | **Melan**<br>**ocytes** | **Radia**<br>**l glial**<br>**cell** | **Tanycy**<br>**tes** | **Fibrob**<br>**lasts** | **Epend**<br>**ymal**<br>**cells** | **Astroc**<br>**ytes** |
| --- | --- | --- | --- | --- | --- | --- | --- | --- | --- |
| *H3F3B* | *TCF7L2* | *RTN1* | *RPS19* | *MLAN*<br>*A* | *C1orf*<br>*61* | *SPARC*<br>*L1* | *LUM* | *CLU* | *ATP1A*<br>*2* |
| *NREP* | *GAP43* | *NREP* | *RPS27* | *PMEL* | *PTN* | *GPM6*<br>*B* | *COL3A*<br>*1* | *SPARC*<br>*L1* | *NTRK2* |
| *SOX4* | *STMN2* | *STMN2* | *EEF1A*<br>*1* | *TYRP*<br>*1* | *GPM*<br>*6B* | *CLU* | *DCN* | *IFITM*<br>*3* | *SLC1A*<br>*3* |
| *NR2F2* | *MAP1B* | *PTMA* | *RPL37* | *ANXA*<br>*2* | *VIM* | *PTN* | *LGALS*<br>*1* | *SPARC* | *SPARC*<br>*L1* |
| *MLLT1*<br>*1* | *STMN1* | *NSG2* | *RPL7* | *GPN*<br>*MB* | *CLU* | *NTRK2* | *COL5A*<br>*1* | *B2M* | *GPM6*<br>*B* |
| *EIF4G*<br>*2* | *SCG2* | *TUBA1A* | *RPLP0* | *DCT* | *PTPR*<br>*Z1* | *ATP1A*<br>*2* | *RPS18* | *RPS27*<br>*L* | *PTN* |
| *RTN* | *NEFL* | *NOVA1* | *RPL18*<br>*A* | *SAT1* | *CNN3* | *PSAT1* | *RPL37* | *SERF2* | *ADGR*<br>*G1* |
| *RBFOX*<br>*2* | *STMN4* | *MLLT11* | *C1orf6*<br>*1* | *QPCT* | *TTYH*<br>*1* | *SLC1A*<br>*3* | *RPL31* | *PLTP* | *CLU* |
| *GAD1* | *PGM2L*<br>*1* | *TMSB10* | *RPL13*<br>*A* | *LGAL*<br>*S1* | *EDNR*<br>*B* | *SPARC* | *RPL13* | *MGST1* | *TTYH1* |

**Supplementary Table 11:** The list of GO cellular component terms and p-values in Supplementary Figure 8.

| GO cellular component | p-value |
| --- | --- |
| Dendrite (GO:0030425) | 0.0033 |
| Anchored component of plasma membrane (GO:0005886) | 0.0254 |
| Rdna heterochromatin (GO:0000781) | 0.0265 |
| Mpp7-dlg1-lin7 complex (GO:0030054) | 0.0265 |
| Endoplasmic reticulum-golgi intermediate compartment membrane (GO:0033116) | 0.0286 |
| Autolysosome (GO:0045324) | 0.0525 |
| Cmg complex (GO:1990429) | 0.0527 |
| Lamellar body (GO:0042628) | 0.0674 |
| Nuclear chromosome (GO:0000228) | 0.0731 |
| Neuron projection (GO:0043005) | 0.0781 |
| Secondary lysosome (GO:0005764) | 0.0838 |
| Axon (GO:0030424) | 0.0977 |

**Supplementary Table 12:**
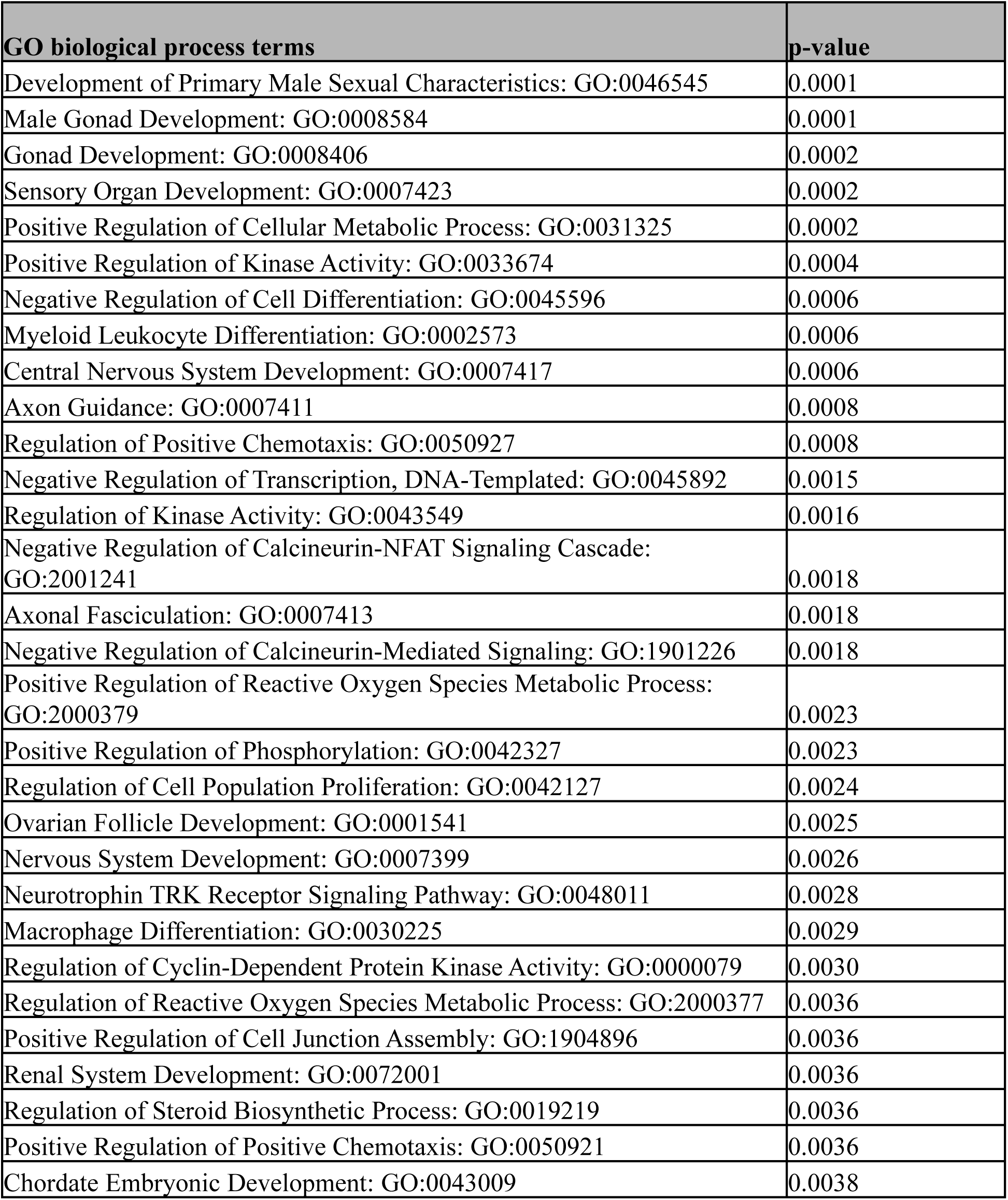
The list of GO biological process terms and p-values in Supplementary Figure 9.

**Supplementary Table 13:** The list of GO molecular function processes and p-values in Supplementary Figure 11.

| **GO molecular function processes** | **p-value** |
| --- | --- |
| Sequence-specific DNA binding (GO:0043565) | 0.003193 |
| Sequence-specific single stranded DNA binding (GO:0098847) | 0.00723 |
| Double-stranded DNA binding (GO:0003690) | 0.013839 |
| Semaphorin receptor activity (GO:0017154) | 0.015839 |
| Single-stranded telomeric DNA binding (GO:0043047) | 0.017266 |
| Sequence-specific double-stranded DNA binding (GO:1990837) | 0.018649 |
| RNA polymerase II core promoter sequence-specific DNA binding (GO:0000979) | 0.020116 |
| RNA polymerase II cis-regulatory region sequence-specific DNA binding (GO:0000978) | 0.023298 |
| Cis-regulatory region sequence-specific DNA binding (GO:0000987) | 0.023298 |
| Actin binding (GO:0003779) | 0.027039 |
| R-SMAD binding (GO:0070412) | 0.027205 |
| NAD<sup>+</sup> adp-ribosyltransferase activity (GO:0003950) | 0.037047 |
| Transcription regulatory region nucleic acid binding (GO:0001067) | 0.037639 |
| Metallocarboxypeptidase activity (GO:0004181) | 0.041235 |
| RNA polymerase II transcription regulatory region sequence-specific DNA binding (GO:0000977) | 0.043676 |
| Telomeric DNA binding (GO:0042162) | 0.046793 |
| Core promoter sequence-specific DNA binding (GO:0001046) | 0.052319 |
| Pentosyltransferase activity (GO:0016763) | 0.053695 |
| Carboxypeptidase activity (GO:0004180) | 0.053695 |
| CCR chemokine receptor binding (GO:0048020) | 0.059183 |
| Cyclin-dependent protein serine/threonine kinase regulator activity (GO:0016538) | 0.061915 |
| Chemokine activity (GO:0008009) | 0.06464 |
| Chemokine receptor binding (GO:0042379) | 0.070066 |
| Receptor ligand activity (GO:0048018) | 0.072694 |
| Nuclear receptor coactivator activity (GO:0030374) | 0.074115 |
| Metalloexopeptidase activity (GO:0008235) | 0.076806 |
| Dna-binding transcription activator activity, RNA polymerase ii-specific (GO:0001228) | 0.083621 |
| Hormone activity (GO:0005179) | 0.107207 |
| DNA binding (GO:0003677) | 0.111491 |
| Growth factor activity (GO:0008083) | 0.118838 |

**Supplementary Table 14:** The list of GO cellular component and p-values in Supplementary Figure 12.

| **GO cellular component** | **p-value** |
| --- | --- |
| Apical dendrite (GO:0097440) | 0.014409 |
| Microtubule (GO:0005874) | 0.028466 |
| Cyclin-dependent protein kinase holoenzyme complex (GO:0000307) | 0.042628 |
| Cul4-ring E3 ubiquitin ligase complex (GO:0080008) | 0.048177 |
| Serine/threonine protein kinase complex (GO:1902554) | 0.052319 |
| Spindle microtubule (GO:0005876) | 0.084831 |
| Plasma membrane raft (GO:0044853) | 0.112394 |
| Intermediate filament cytoskeleton (GO:0045111) | 0.114977 |
| Early endosome membrane (GO:0031901) | 0.13159 |
| Asymmetric synapse (GO:0032279) | 0.176037 |
| Postsynaptic density (GO:0014069) | 0.182034 |
| Nucleus (GO:0005634) | 0.184057 |
| Cullin-ring ubiquitin ligase complex (GO:0031461) | 0.20444 |
| Membrane raft (GO:0045121) | 0.211391 |
| Cytoskeleton (GO:0005856) | 0.215757 |
| Spindle (GO:0005819) | 0.244172 |
| Axon (GO:0030424) | 0.257348 |
| Polymeric cytoskeletal fiber (GO:0099513) | 0.311926 |
| Early endosome (GO:0005769) | 0.321968 |
| Lytic vacuole membrane (GO:0098852) | 0.322965 |
| Dendrite (GO:0030425) | 0.325945 |
| Intracellular membrane-bounded organelle (GO:0043231) | 0.330228 |
| Endosome membrane (GO:0010008) | 0.378398 |
| Lysosomal membrane (GO:0005765) | 0.382966 |
| Microtubule cytoskeleton (GO:0015630) | 0.383876 |
| Collagen-containing extracellular matrix (GO:0062023) | 0.4269 |
| Focal adhesion (GO:0005925) | 0.432804 |
| Cell-substrate junction (GO:0030055) | 0.438649 |

**Supplementary Table 15:**
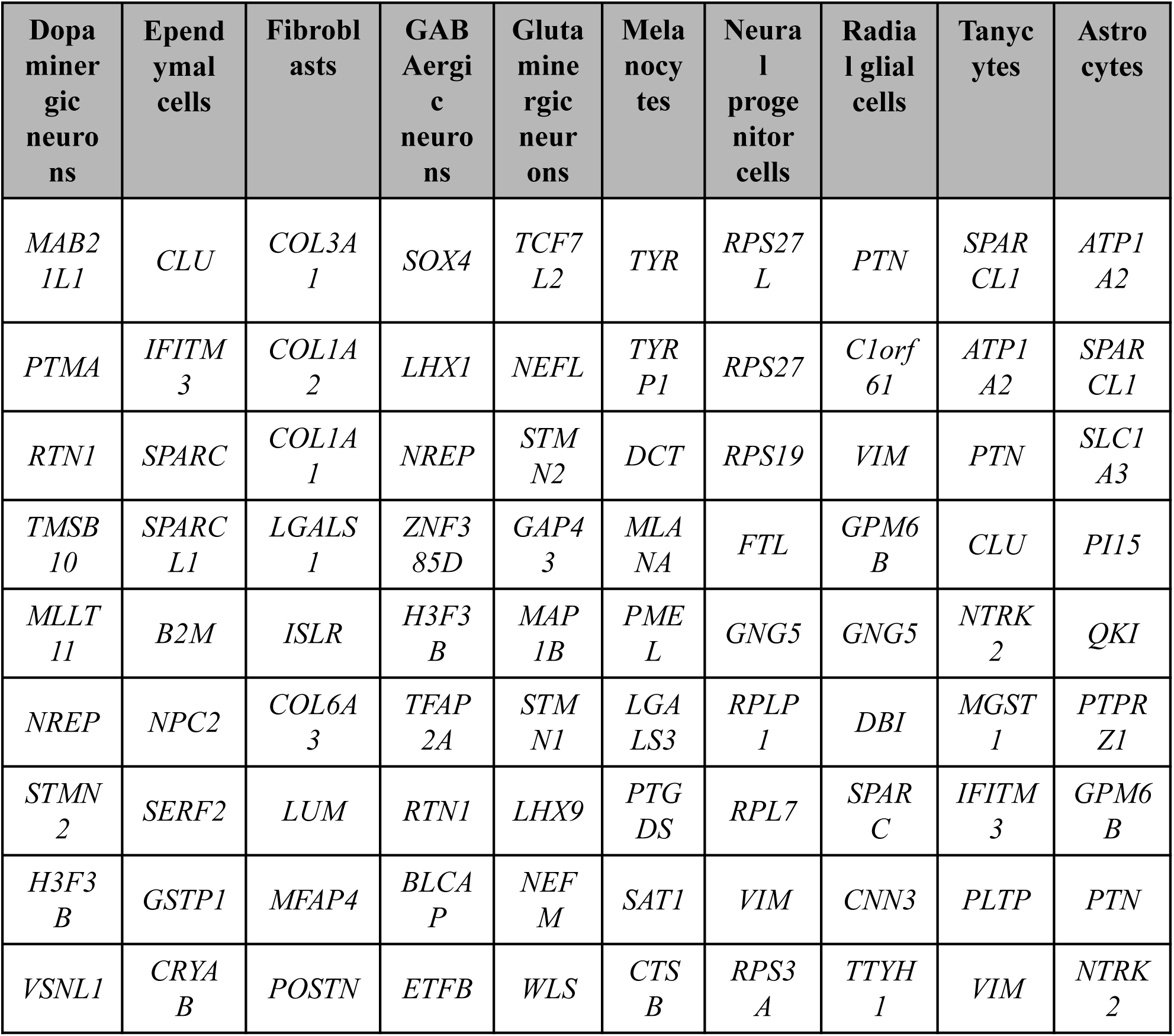
The list of top 10 genes for Figure 10A-D in patient cortical organoids with metformin treatment of cell type enrichment analysis in scRNA-seq analysis.

**Supplementary Table 16:** The list of GO terms and p-values in Supplementary Figure 16.

| **GO terms** | **p-value** |
| --- | --- |
| Regulation of neurotransmitter transport: GO:1902476 | 0.41704493 |
| Regulation of sensory perception: GO:0051960 | 0.41704494 |
| Regulation of dendritic cell apoptotic process: GO:2000252 | 0.41704495 |
| Inositol phosphate dephosphorylation: GO:0043648 | 0.41704496 |
| Columnar/cuboidal epithelial cell differentiation: GO:0002064 | 0.41704497 |
| DNA damage response, signal transduction resulting in transcription: GO:0006978 | 0.41704498 |
| Phosphorylated carbohydrate dephosphorylation: not a specific GO term | 0.41704499 |
| Enteric nervous system development: GO:0048484 | 0.41704500 |
| Diterpenoid metabolic process: GO:0016103 | 0.41704501 |
| Regulation of pseudopodium assembly: GO:0031274 | 0.41704502 |
| Regulation of sister chromatid cohesion: GO:0031135 | 0.41704503 |
| Dendritic cell chemotaxis: GO:1990736 | 0.41704504 |
| Fat-soluble vitamin catabolic process: GO:0042364 | 0.41704505 |
| Negative regulation of leukocyte apoptotic process: GO:2000271 | 0.41704506 |
| Inositol phosphate catabolic process: GO:0016312 | 0.41704507 |
| Positive regulation of pseudopodium assembly: GO:0031275 | 0.41704508 |
| Cochlea morphogenesis: GO:0090102 | 0.41704509 |
| Depurination: GO:0006284 | 0.41704510 |
| Phasic smooth muscle contraction: not a specific GO term | 0.41704511 |
| Regulation of protein localization: GO:0032880 | 0.41704512 |
| Positive regulation of lipid localization: GO:1905953 | 0.41704513 |
| Response to prostaglandin E: GO:0071404 | 0.41704514 |
| DNA damage response, signal transduction by p53 class mediator resulting in transcription of p21 class mediator: GO:0006979 | 0.41704515 |
| Mitotic DNA replication: GO:1902275 | 0.41704516 |
| Positive regulation of sequestering of triglyceride: not a specific GO term | 0.41704517 |
| Regulation of mitotic centrosome separation: GO:0051298 | 0.41704518 |
| Very-low-density lipoprotein particle remodeling: GO:0034376 | 0.41704519 |
| Embryonic forelimb morphogenesis: GO:0035115 | 0.41704520 |
| Regulation of reactive oxygen species metabolic process: GO:2000377 | 0.41704521 |
| Renal system development: GO:0072001 | 0.41704522 |

**Supplementary Table 17:** The list of KEGG pathways and p-values in Supplementary Figure 17.

| **KEGG pathways** | **p-value** |
| --- | --- |
| Dorso ventral axis formation | 0.0120 |
| Polyunsaturated fatty acid biosynthesis | 0.0806 |
| Prion disease | 0.0813 |
| Calcium signaling pathway | 0.0878 |
| ERBB signaling pathway | 0.0952 |
| Prostate cancer | 0.0952 |
| Renin angiotensin system | 0.0970 |
| Chondroitin sulfate biosynthesis | 0.1237 |
| Arginine and proline metabolism | 0.1908 |
| Neurodegenerative diseases | 0.2050 |
| Bladder cancer | 0.2222 |
| Valine leucine and isoleucine degradation | 0.2331 |
| Glycine serine and threonine metabolism | 0.2387 |
| Type I diabetes mellitus | 0.2387 |
| Endometrial cancer | 0.2702 |
| Non small cell lung cancer | 0.2754 |
| Basal cell carcinoma | 0.2861 |
| Hedgehog signaling pathway | 0.2916 |
| Glycolysis and gluconeogenesis | 0.3207 |
| Glioma | 0.3207 |
| Epithelial cell signaling in helicobacter pylori infection | 0.3396 |
| Focal adhesion | 0.3412 |
| Long term potentiation | 0.3412 |
| VEGF signaling pathway | 0.3445 |
| PPAR signaling pathway | 0.3445 |
| Melanoma | 0.3511 |
| Pancreatic cancer | 0.3596 |
| Adherens junction | 0.3682 |
| Antigen processing and presentation | 0.3936 |
| Apoptosis | 0.3974 |

**Supplementary Table 18:** The list of GO cellular component and p-values in Supplementary Figure 19.

| GO cellular component | P-value |
| --- | --- |
| Integral component of mitochondrial outer membrane (GO:0031307) | 0.000711 |
| Intrinsic component of mitochondrial outer membrane (GO:0031306) | 0.000778 |
| Filopodium (GO:0030175) | 0.004888 |
| Integral component of mitochondrial membrane (GO:0032592) | 0.006285 |
| Phosphatidylinositol 3-kinase complex, class I (GO:0097651) | 0.008968 |
| Actin-based cell projection (GO:0098858) | 0.009783 |
| Cell cortex region (GO:0099738) | 0.019627 |
| Filopodium membrane (GO:0031527) | 0.021393 |
| Mitochondrial outer membrane (GO:0005741) | 0.021573 |
| Trans-golgi network transport vesicle (GO:0030140) | 0.026671 |
| Organelle outer membrane (GO:0031968) | 0.026938 |
| Membrane raft (GO:0045121) | 0.034704 |
| Synaptic membrane (GO:0097060) | 0.04579 |

